# Admixture mapping reveals evidence for multiple mitonuclear incompatibilities in swordtail fish hybrids

**DOI:** 10.1101/2025.01.30.635158

**Authors:** Nemo V. Robles, Benjamin M. Moran, María José Rodríguez-Barrera, Gaston I. Jofre, Theresa Gunn, Erik N.K. Iverson, Sofia Beskid, JJ Baczenas, Alisa Sedghifar, Peter Andolfatto, Daniel L. Powell, Yaniv Brandvain, Justin C. Havird, Gil G Rosenthal, Molly Schumer

## Abstract

How barriers to gene flow arise between closely related species is one of the oldest questions in evolutionary biology. Classic models in evolutionary biology predict that negative epistatic interactions between variants in the genomes of diverged lineages, known as hybrid incompatibilities, will reduce viability or fertility in hybrids. The genetic architecture of these interactions and the evolutionary paths through which they arise have profound implications for the efficacy of hybrid incompatibilities as barriers to gene flow between species. While these questions have been studied using theoretical approaches for several decades, only recently has it become possible to genetically map larger numbers of hybrid incompatibilities. Here, we use admixture mapping in natural hybrid populations of swordtail fish (*Xiphophorus*) to identify hybrid incompatibilities involving genetic interactions between the mitochondrial and nuclear genomes. We find that at least nine regions of the genome are involved in mitonuclear incompatibilities. These incompatibilities involve interactions between the nuclear genome and the *X. malinche* mitochondria, the *X. birchmanni* mitochondria, or both. Moreover, they vary in the strength of selection they experience, and the degree to which they limit gene flow in natural hybrid populations. Our results build a deeper understanding of the complex architecture of selection against incompatibilities in naturally hybridizing species and highlight an important role of mitonuclear interactions in the evolution of reproductive barriers between closely related species.

## Introduction

As lineages diverge, mutations that differentiate them will arise and ultimately fix due to the action of genetic drift or natural selection. One of the foundational theories in evolutionary biology is that combinations of these distinct variants in hybrids can lead to ‘incompatible’ genetic interactions that reduce hybrid viability or fertility. Since this idea was first proposed by Dobzhansky and Muller^1,2^, scores of studies have now mapped negative epistatic interactions that lead to reduced viability or fertility in hybrids^3–11^. Decades of theoretical work have established the importance of hybrid incompatibilities as barriers to gene flow and mechanisms through which species can become and remain reproductively isolated^12–20^. More recently, advances in genomics have fueled the identification of dozens of individual genes involved in hybrid incompatibilities^21–24^. However, only a small subset of these studies have successfully mapped hybrid incompatibilities in naturally hybridizing species^6,22,25,26^. As a result, few studies have examined the importance of hybrid incompatibilities as barriers to gene flow in nature^21–24^.

Thus, despite their predicted importance in the formation and maintenance of species, many open questions remain about the evolution of hybrid incompatibilities and their consequences in natural populations. In part due to the experimental challenges of precisely mapping the genes involved in hybrid incompatibilities^27^, theoretical work has vastly outpaced empirical work in this area. Classic theoretical research in evolutionary biology predicts that hybrid incompatibilities may be more likely to arise between rapidly evolving genes, as these will be the first to accumulate functionally important substitutions that differ between species^12,17^. Some existing empirical results are consistent with this interpretation, with several known incompatibility genes showing elevated rates of amino acid substitutions or high rates of structural evolution^6,22,28^.

Since these initial theoretical results in population genetics, advances in systems biology have led to new predictions about the nature of genetic interactions that might lead to genetic breakdown in hybrids and the mechanisms through which this could evolve. Experimental approaches have generated comprehensive maps of gene interactions in model species such as *Saccharomyces* and *C. elegans*, and have led to the realization that the majority of genes have few genetic interactions, while others act as “hubs,” with many interacting partners^29–32^. At the same time, theoretical and empirical advances^33,34^ have indicated that conserved traits and pathways can diverge in their developmental and genetic underpinnings over evolutionary timescales, a process known as developmental systems drift^33,34^. Several newly mapped hybrid incompatibilities have been implicated in conserved developmental processes whose molecular basis appears to have diverged over long evolutionary timescales (e.g. ^35,36^) and thus cause dysfunction in hybrids. Notably, compensatory coevolution in interacting proteins, where substitutions that impact function in one protein are restored through changes in an interacting protein, has been thought to be an important mechanism underlying developmental systems drift (though empirical evidence is mixed^37^).

Among protein complexes, some researchers have suggested that interactions between the mitochondrial and nuclear genome may be especially prone to evolving differences that generate incompatibilities in hybrids^38^. Nearly 1,500 nuclear-encoded proteins localize to the mitochondria in vertebrates, and more than one hundred form physical complexes with mitochondria-encoded proteins. At the same time, different error-correction machinery used by the mitochondrial genome often leads to higher substitution rates in mitochondrial-encoded genes (up to ∼20X in vertebrates^39^). This is often mirrored by elevated substitution rates in nuclear-encoded genes that must interact intimately with their mitochondrial partners^40^. Other features of mitochondrial biology, including the lack of meiotic recombination and potential for sexual conflict may also impact dynamics of evolution and coevolution in mitochondrial and nuclear genes. Thus, interactions between mitochondrial- and nuclear-encoded proteins may be predisposed to the evolution of hybrid incompatibilities due to their molecular and evolutionary properties.

While this hypothesis has been challenging to evaluate since only a few mitonuclear hybrid incompatibilities have been precisely mapped^25,41–43^, the broad predictions of this model are well supported. Hybrids between many species often show viability or fertility effects that depend on the maternal parent in the cross, consistent with a role for the mitochondrial genome in hybrid fitness (as well as other mechanisms^44^). Physiological approaches have also highlighted widespread mitochondrial dysfunction in hybrids that may point to suboptimal interactions between the mitochondrial- and nuclear-derived proteins of the parent species^42,45–51^. With an improved understanding of the genetic architecture of selection on mitonuclear interactions in hybrids, and epistatic interactions in hybrids more generally, researchers can better understand the consequences of hybridization between species from both a genetic and evolutionary perspective.

Swordtail fish of the genus *Xiphophorus* have become a model system for the study of speciation. Several pairs of species in this genus naturally hybridize^23,52–54^, providing unique datasets to study the impacts of hybridization. Among these, some of the best studied natural hybrid populations are those that have formed between sister species *Xiphophorus birchmanni* and *X. malinche* (Fig. 1A)*. X. birchmanni* and *X. malinche* are native to the Sierra Madre Oriental of Mexico and hybridize in multiple river systems where their ranges overlap (Fig. 1B). Despite diverging only an estimated 250,000 generations before the present^55^ (∼0.4% pairwise sequence divergence), these species have multiple known hybrid incompatibilities^6,25^, with evidence for perhaps dozens more from population genetic and cross data^23,27,56^.

**Fig. 1.**
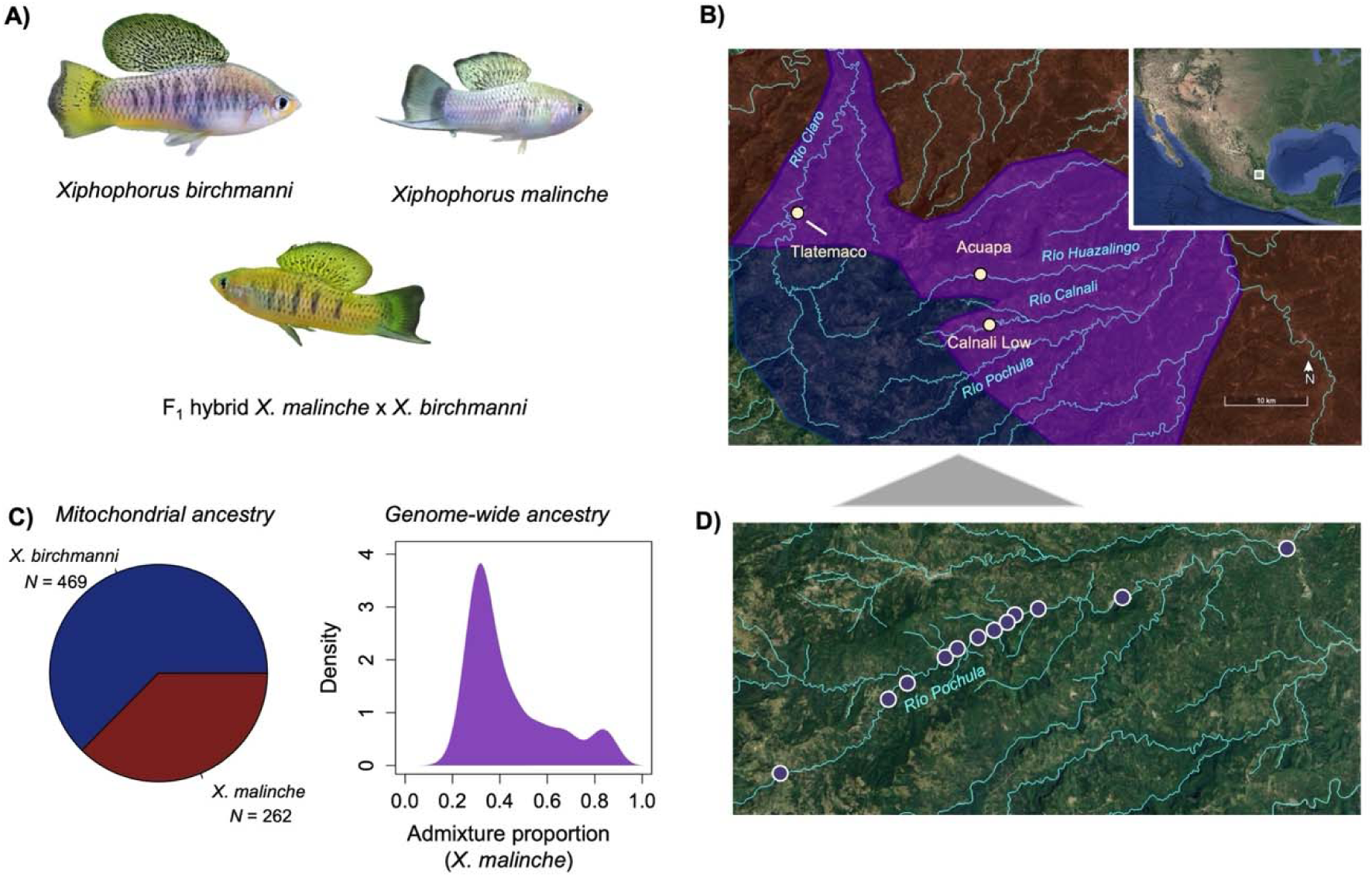
**A)** Photograph of *X. birchmanni* male, *X. malinche* male and an F_1_ hybrid male. **B)** Sampling locations where *X. birchmanni* and *X. malinche* overlap and hybridize that were used for admixture mapping (Calnali Low hybrid population) and local ancestry analyses (Tlatemaco and Acuapa hybrid populations). Rivers from which data was included in this study are labeled in blue on the map. Shading on the map corresponds to the range of *X. malinche* (red), *X. birchmanni* (blue), and hybrids (purple). **C)** Summary of mitochondrial and genome-wide ancestry for individuals used in admixture mapping, which were sampled from the Calnali Low hybrid population on the Río Calnali. Left – proportion of individuals in the admixture mapping population with *X. birchmanni* or *X. malinche* mitochondrial haplotypes. Right – distribution of genome-wide admixture proportion across individuals used in admixture mapping in this study. Admixture proportion is summarized as the proportion of the genome derived from the *X. malinche* parent species. Samples shown here reflect data from 731 samples collected from 2018-2022. **D)** Map of sampling locations of populations included in the cline analysis along the Río Pochula.

In previous work, we used a combination of admixture mapping and analysis of segregation distortion in F_2_ hybrids to fine-map two nuclear-encoded genes involved in a lethal mitonuclear hybrid incompatibility between *X. birchmanni* and *X. malinche*: one on chromosome 6 involving an interaction between *X. birchmanni ndufa13* and the *X. malinche* mitochondria and one on chromosome 13 involving an interaction between either species’ version of *ndufs5* and the mitochondria of the other species^25^. Both of these genes encode proteins in Complex I of the mitochondrial electron transport chain, and physiological and proteomic data indicated that ancestry mismatch at these loci results in Complex I dysfunction. Ultimately this dysfunction leads to embryonic lethality or mortality shortly after birth^25^. We also detected a mitonuclear incompatibility involving chromosome 15 but were unable to localize the driver of this signal^25^.

Here, we perform a higher-powered scan for genes involved in mitonuclear incompatibilities between *X. birchmanni* and *X. malinche* by doubling the sample size of our initial study^25^. We replicate previously detected incompatibilities and find strong evidence for several additional mitonuclear incompatibilities including incompatibilities that are physically linked to those we previously mapped (Table 1). In some cases, we are able to fine-map these interactions to the genes likely to be involved, document evidence of physiological consequences on mitochondrial function, and identify selection on these regions in natural hybrid populations. Our results further underscore the importance of mitonuclear interactions in the evolution of hybrid incompatibilities between closely related species.

**Table 1.**
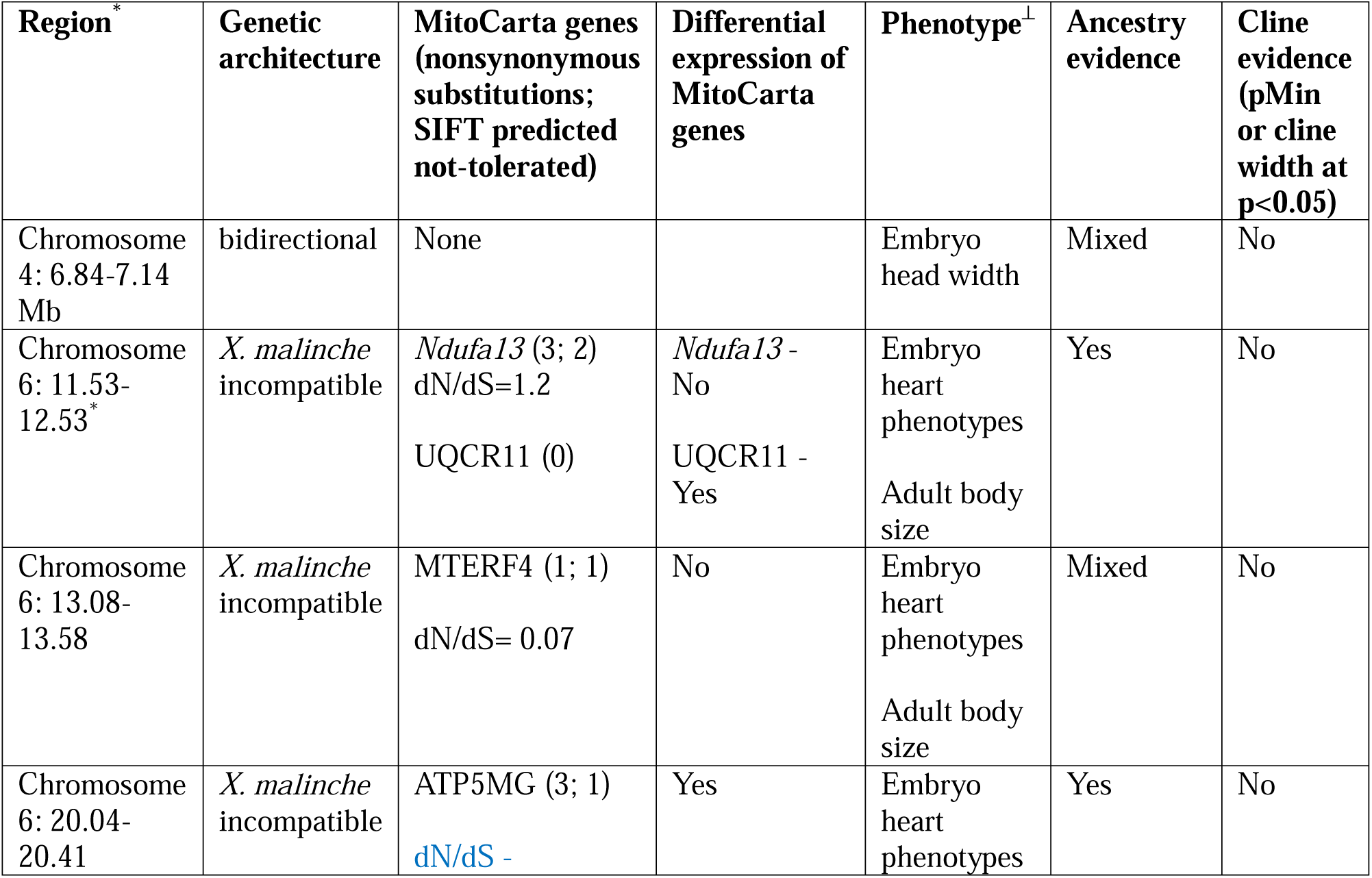
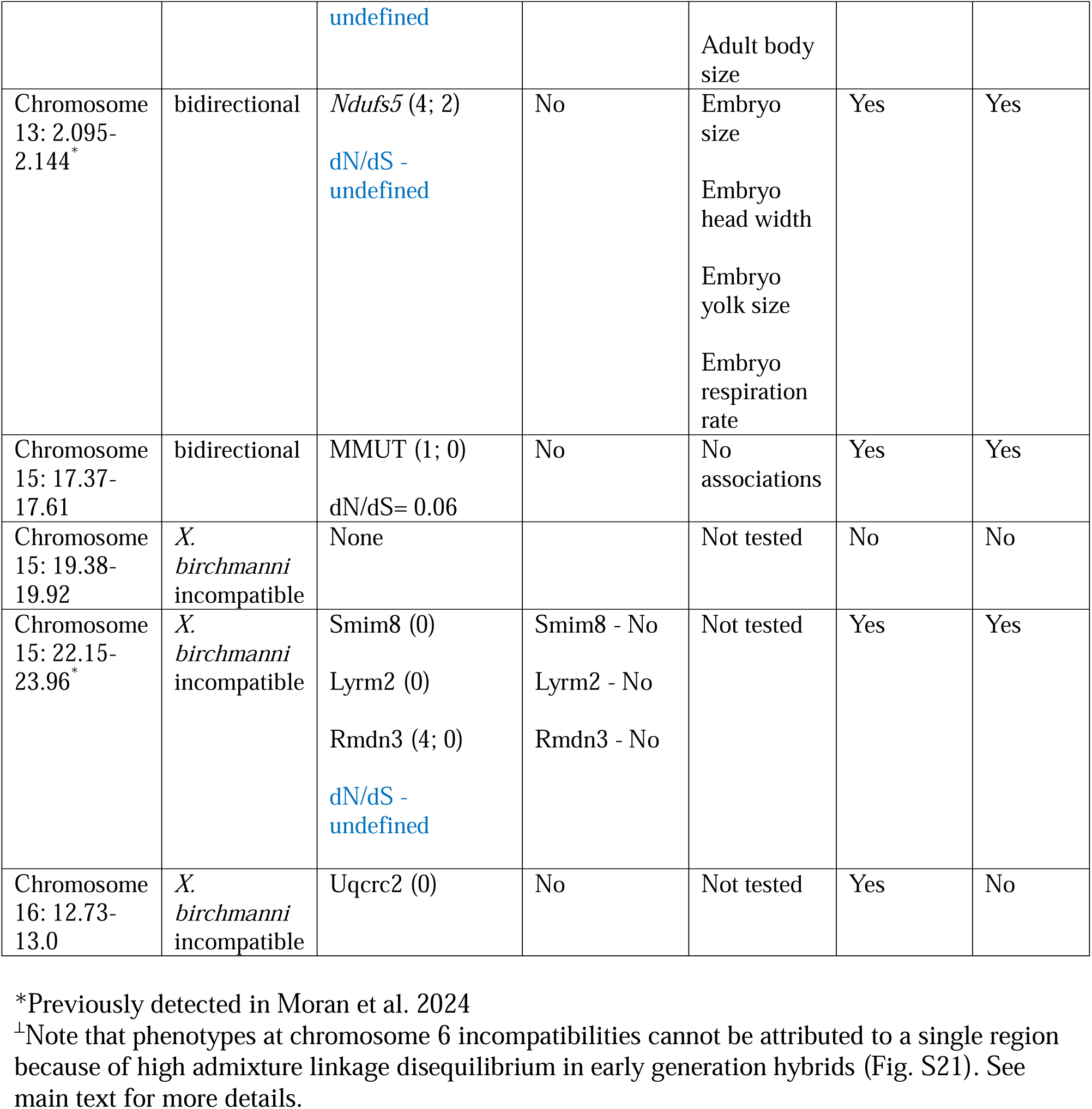
Summary of results for each admixture mapping peak detected in our analyses. Genetic architecture refers to whether incompatibilities were inferred to involve interactions with the *X. birchmanni* mitochondria, the *X. malinche* mitochondria, or both (see Fig. 3; Supporting Information 1). Annotated MitoCarta genes detected in the admixture mapping interval are listed, as are the number of nonsynonymous substitutions in those protein sequences that differ between *X. birchmanni* and *X. malinche*, dN/dS between *X. birchmanni* and *X. malinche* for each MitoCarta gene, and the number of substitutions observed in either species that were predicted not tolerated based on SIFT analysis. We also list whether there was evidence of differential expression of MitoCarta genes between *X. birchmanni* and *X. malinche* in a previously analyzed RNAseq dataset^77^. For incompatibilities involving the *X. malinche* mitochondria, we tested for phenotypic effects on embryonic size, embryonic respiration, embryonic heart morphology and rate (Supporting Information 3), and on adult size (Methods). Associated phenotypic effects are listed in the “Phenotype” column. In the Ancestry evidence and Cline evidence columns, we note whether there is evidence of selection on the region in natural hybrid populations. Ancestry tests were performed in the Acuapa and Tlatemaco populations, and cline analyses were performed in the Río Pochula. For the ancestry evidence, ancestry at focal regions was compared to the genome-wide background. For bidirectional incompatibilities where we expect depletion of mismatched ancestry in both the Acuapa and Tlatemaco populations, a subset of columns list “Mixed,” indicating that ancestry was depleted in only one population. For cline evidence, ancestry at focal regions was compared to matched nulls, and we indicate whether the region was an outlier in minimum allele frequency (pMin) or in cline width.

## Materials and Methods

### Sample collection and curation

For the admixture mapping analyses described in this paper, we combined a previously published dataset of 359 natural hybrids from the Calnali Low hybrid population^25^ with newly collected data from 372 additional hybrids from the same population (Fig. 1C). By approximately doubling our sample size, we expected to increase our power to detect incompatible interactions with more modest selection coefficients. Samples were collected using baited minnow traps. Individuals were anesthetized in 100 mg/mL MS-222 and a small fin clip was taken from the upper caudal fin of each individual (following Stanford APLAC protocol #33071). Fish were allowed to recover in a holding tank and then returned to the site where the trap was deployed. Fin clips were stored in 95% ethanol for later DNA extraction and sequencing.

For this study, we also included analysis of samples across a geographical cline along the hybrid zone in the Río Pochula. We collected samples from 12 locations along the Río Pochula, ranging from elevations of 221 meters to 1400 meters (Fig. 1D). The number of samples per geographical location ranged from 16 to 45. Full details of sampling localities on the Río Pochula can be found in Table S1. Individuals were collected and fin clipped as described above. Finally, we collected fin clips from 805 F_2_ hybrids generated in our fish facility at Stanford to supplement data collected in our original study^25^ for lab-generated hybrids. We followed the same sample collection procedure to fin clip artificial hybrids generated in the laboratory, except that individuals were tagged with a unique elastomer fluorescent tag for later matching of individuals and their genotypes. We also reanalyzed several datasets generated from natural hybrid populations that were previously published. We summarize all the datasets analyzed in the manuscript in Table S2.

### DNA extraction and library preparation

We prepared fin clips for low-coverage whole genome sequencing as described previously^23,25,57^. Briefly, we extracted DNA from each fin clip individually in a 96-well plate format using the Agencourt DNAdvance kit (Beckman Coulter, Brea, California), following the manufacturer’s instructions (with half-reactions). Extracted DNA was quantified using a TECAN Infinite M1000 plate reader (Tecan Trading AG, Switzerland). Each sample was diluted to 10 ng/ul and libraries were prepared in a 96-well plate format. DNA was enzymatically sheared and initial adapter sets were added using the Illumina Tagment DNA TDE1 Enzyme and Buffer Kit. Following this reaction, each sample was amplified with dual-indexed primers for 12 cycles using the OneTaq HS Quick-load PCR mastermix. The samples from these resulting PCR reactions were pooled and purified using 18% SPRI magnetic beads. Libraries were sequenced on a HiSeq 4000 at Admera Health (South Plainfield, NJ, USA).

### Local ancestry inference

We relied on methods established by our group for hidden Markov model-based local ancestry inference of *X. birchmanni* x *X. malinche* hybrids across the nuclear and mitochondrial genome^58^. We followed the methods described in Moran et al.^25^ for local ancestry inference using the *ancestryinfer* pipeline^58^, except that we used updated chromosome-scale versions of both the *X. birchmanni* and *X. malinche* genomes generated using PacBio HiFi data^24^. Briefly, we defined candidate ancestry informative sites using a set of high coverage *X. birchmanni* and *X. malinche* individuals, as we had previously^25^. We mapped reads to the *X. birchmanni* reference genome using bwa-mem^59^, identified and removed duplicates with PicardTools^60^, and performed variant calling with GATK^61^, applying hard calls to filter variants that minimized mendelian errors in previous analyses^25,55^. See Moran et al.^25^ for a detailed description of the variant calling pipeline and quality thresholds.

We identified sites that were called for the reference base across *X. birchmanni* individuals and called for the alternate based in *X. malinche*. We treated these as our candidate ancestry informative sites. We next verified that these sites were ancestry informative with a large sample of low-coverage population data from each species (N=126 *X. birchmanni*; N=38 *X. malinche*). Note the *X. malinche* has ∼4X lower genetic diversity than *X. birchmanni*^55^. We calculated allele frequency in the parental populations at each candidate ancestry informative site using bcftools mpileup^62^ and removed ancestry informative markers with <98% frequency difference between the two species. This resulted in a total of 729,167 ancestry informative sites across the 24 *Xiphophorus* chromosomes, or ∼1 informative site per kb. Performance of ancestry inference using *ancestryinfer*^58^ and these informative sites was tested on individuals from the parental populations that were not used in the filtering dataset (N=48 *X. birchmanni*; N=36 *X. malinche*), as well as known F_1_ hybrids between the two species from laboratory crosses (N=52). Based on these analyses, we conclude that the ancestry inference error rate is extremely low (<0.1% per ancestry informative site). This mirrors results from previous versions of our ancestry inference pipeline^23–25^ and simulation-based tests of pipeline performance^58^.

To infer local ancestry from the Calnali Low hybrid population, we ran the *ancestryinfer* pipeline, which has been described in detail elsewhere^6,23,25,57,58^. Briefly, the *ancestryinfer* pipeline maps reads to both reference genomes using bwa-mem^59^, uses the ngsutilsj^63^ package to remove reads that do not map uniquely to *both* references and remove reads with mapping quality less than 30, uses bcftools mpileup^62^ to count the number of reads supporting each allele at each ancestry informative site, reformats the data for input into the AncestryHMM program and runs AncestryHMM^64^. We ran *ancestryinfer* setting the prior admixture proportion to 50% *X. malinche* and the estimated time since initial admixture to 40 generations based on past analyses of this population^25^. We set the expected error rate to 2% and the expected per basepair recombination rate to 2×10^-6^ cM/bp. Past work has suggested that the HMM implemented in *ancestryinfer* is relatively insensitive to prior misspecification^58^.

*ancestryinfer* outputs posterior probabilities of ancestry (homozygous parent 1, heterozygous, or homozygous parent 2) at each ancestry informative site along the chromosome. To post-process this data, we used a threshold of 0.9 to convert these posterior probabilities into “hard-calls.” For sites in an individual where a given ancestry state was supported at ≥0.9 posterior probability, the site was converted to that ancestry state. If no ancestry state at a given site was supported by a ≥0.9 posterior probability, that site was converted to NA. This dataset was used an input into admixture mapping analyses.

### Admixture mapping

We used an admixture mapping approach to identify regions across the nuclear genome that show an unexpectedly strong association with mitochondrial ancestry (Fig. 2). The Calnali Low population, hereafter the “admixture mapping population”, is one of the few natural hybrid populations between *X. birchmanni* and *X. malinche* that segregates for both mitochondrial haplotypes^25^ (Fig. 1C). To identify interactions with mitochondrial ancestry, we treated the individual’s mitochondrial haplotype (*X. birchmanni* or *X. malinche*) as the phenotype of interest (Fig. 3). Natural selection that disproportionately removes particular ancestry combinations can generate unexpectedly high correlations in ancestry between physically unlinked loci. We used a partial correlation approach^27^ to evaluate the correlation between nuclear and mitochondrial ancestry while accounting for the covariance in ancestry expected given each individual’s admixture proportion. For each focal ancestry informative site, we recorded the p-value from the correlation in mitochondrial and nuclear ancestry after accounting for genome-wide admixture proportion (using the ppcor package in R). We excluded ancestry informative sites with high levels of missing data from our analysis (more than 15% of individuals in the dataset missing). We compared the observed data with null simulations, described in the next section.

**Fig. 2.**
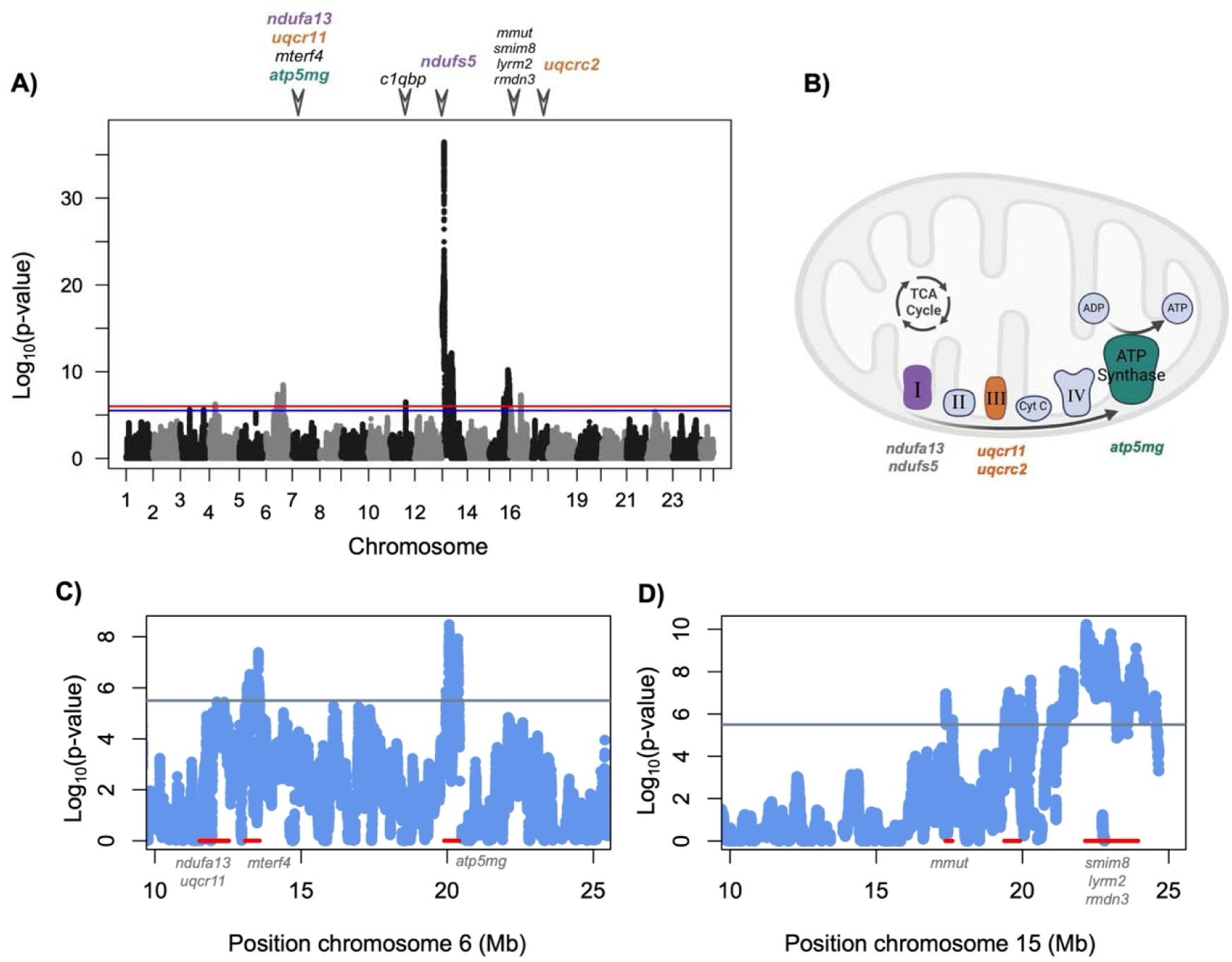
**A)** Admixture mapping results for the association between mitochondrial and nuclear ancestry across the genome. Red line represents the 5% false positive rate threshold, blue line represents the 10% false positive rate threshold. Triangles and gene names indicate MitoCarta annotated genes associated with each admixture mapping interval. Separate intervals on the same chromosome are collapsed for visualization purposes. Colored text highlights mitochondrially interacting genes that localize to particular protein complexes (see **B**). See Fig. S20 for a version of this figure plotted with a truncated y-axis for better visualization of signals close to the genome-wide significance threshold. **B**) Illustration adapted from BioRender of the mitochondrial electron transport chain. Each component of the electron transport chain is indicated and labeled; ATP synthase is synonymous with Complex V. Purple, orange, and green complexes indicate complexes that were implicated in mitonuclear incompatibilities based on our admixture mapping results. The newly mapped incompatibilities that form parts of particular mitochondrial complexes are listed below those complexes in the corresponding color. Incompatibilities previously mapped by Moran et al.^25^ are listed in gray text. **C)** Admixture mapping results for chromosome 6 highlight multiple regions that surpass the genome-wide significance threshold (gray line indicates 10% false positive rate threshold). Although these regions are linked in early generation hybrids (Fig. S21), they are not in strong linkage disequilibrium in the admixture mapping population (Fig. S4). **D)** Admixture mapping results for chromosome 15 highlight multiple regions that surpass the genome-wide significance threshold (gray line indicates 10% false positive rate threshold). These regions are also not in strong linkage disequilibrium in the admixture mapping population (Fig. S4).

**Fig. 3.**
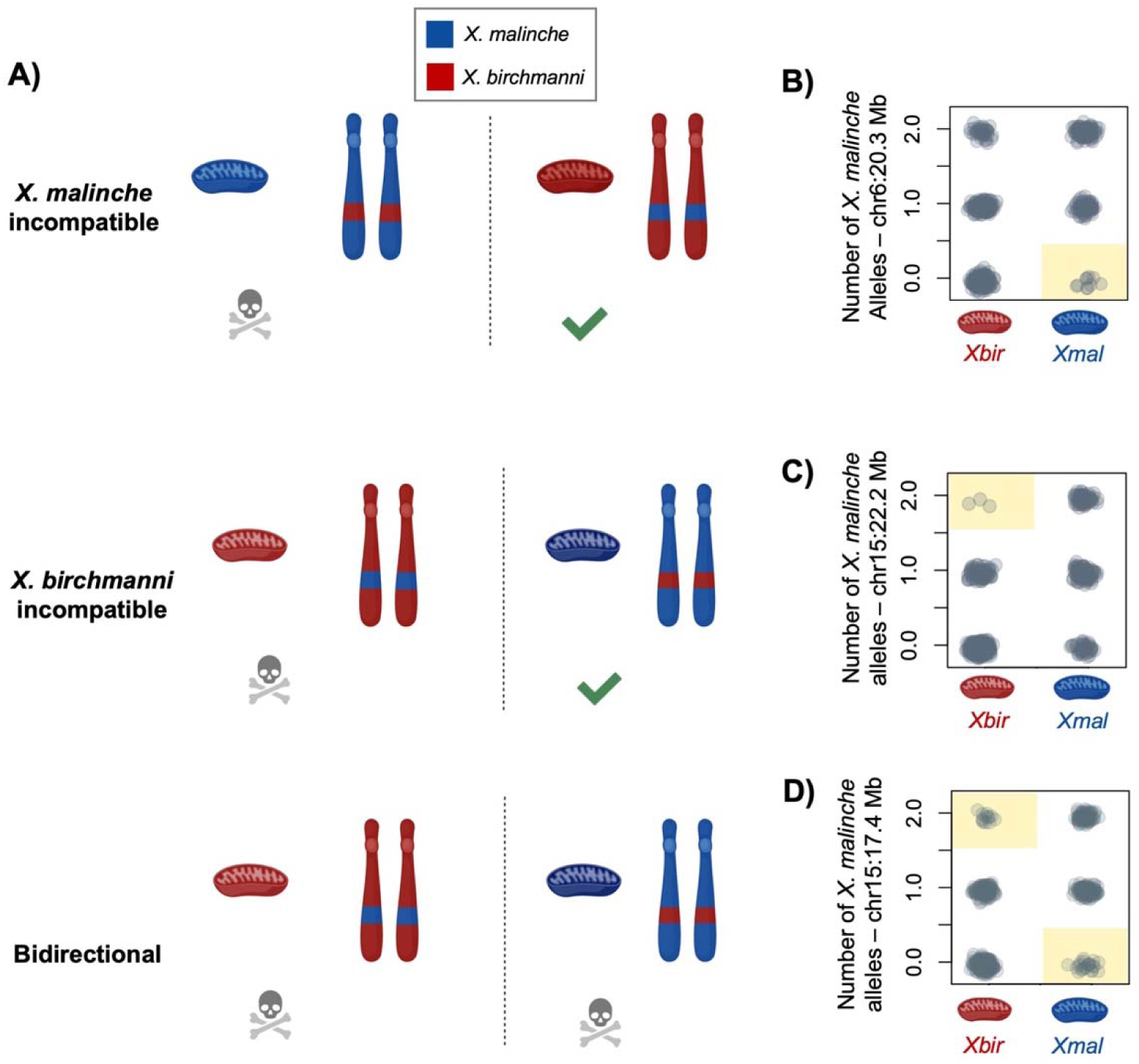
**A)** Architecture of mitonuclear incompatibilities inferred from our data. Red shading represents individuals with *X. birchmanni* haplotypes and the blue shading represents individuals with *X. malinche* haplotypes. Mitochondrial haplotypes are indicated with mitochondria illustrations and nuclear haplotypes are indicated with chromosome illustrations. *X. malinche* incompatible interactions are cases where the *X. malinche* mitochondria is incompatible with *X. birchmanni* ancestry at the nuclear locus (top). *X. birchmanni* incompatible interactions are cases where the *X. birchmanni* mitochondria is incompatible with *X. malinche* ancestry at the nuclear locus (middle). Bidirectional incompatible interactions are cases where both mitochondrial types are incompatible with heterospecific ancestry at the nuclear locus (bottom). **B)** Empirical example of a *X. malinche* incompatible interaction. Each gray point indicates one adult individual in our admixture mapping population at 20.3 Mb on chromosome 6. Red mitochondria on the x-axis indicates individuals with *X. birchmanni* (*Xbir*) mitochondrial haplotypes and blue mitochondria indicates individuals with *X. malinche* (*Xmal*) mitochondrial haplotypes. Genotype combinations that are significantly depleted based on comparisons to null simulations are highlighted with yellow shading (see Supporting Information 1). **C)** Empirical example of a *X. birchmanni* incompatible interaction at 22.2 Mb on chromosome 15, with plot information following **B**. **D)** Empirical evidence of a bidirectionally incompatible interaction at 17.4 Mb on chromosome 15, with plot information following **B**. Illustrations in this figure were produced by BioRender. For similar plots for the remaining mitonuclear incompatibilities mapped in this or previous work, see Figs. S1, S22.

### Simulations to determine the genome-wide significance threshold for admixture mapping

To determine the appropriate genome-wide significance threshold for our admixture mapping analysis, we investigated the distribution of p-values for associations between nuclear and mitochondrial ancestry when there was no true relationship between the loci. While permutations of the observed data are often used to generate these expectations, such an approach is likely inappropriate here because individuals in our population vary widely in genome wide ancestry (Fig. 1C), which will drive correlations in ancestry between any two loci simply due to population structure^27^. In order to account for this issue, we instead used observed genome-wide ancestry to simulate a mitochondrial haplotype for each individual. To do so, we used the random binomial function in R and set the probability of drawing a zero or one to the proportion of an individual’s genome derived from the *X. malinche* parental species. If we drew a zero, we set the mitochondrial haplotype to *X. birchmanni* for that individual in that simulation. If we drew a one, we set the mitochondrial haplotype to *X. malinche* for that individual in that simulation. We repeated this procedure until all individuals had a simulated mitochondrial haplotype. Next, we performed admixture mapping as we had for the real data and recorded the minimum p-value observed in that simulation. We repeated this for 500 replicate simulations, yielding a distribution of 500 minimum p-values. For our significance threshold, we used the lower 5% tail of the simulated p-values, roughly corresponding to an expected false positive rate of 5%. Based on these simulations, we set the genome-wide significance threshold to p< 1 x 10^-6^. In addition, we evaluated whether more regions were detected at a relaxed threshold corresponding to a 10% false positive rate, corresponding to p<3 x 10^-6^.

We also used a simulation-based approach to infer the likely architecture of each mitonuclear incompatibility. Specifically, we wanted to determine whether particular nuclear genotypes in our dataset were depleted in combination with the *X. malinche* mitochondria, *X. birchmanni* mitochondria, or both. This approach is described in detail in Supporting Information 1 and results for the inferred incompatibility architecture are summarized in Table 1.

### Defining the association interval and identifying candidate genes

For regions that exceeded our genome-wide significance threshold, we needed to determine how to delineate the associated region for further analysis. To be conservative, we included the entire region that fell within ± 2 of the peak Log_10_ (p-value). For example, if the peak association was 10 (i.e. p<10^-10^), we identified the region on both sides of the peak site where the significance of the association exceeded 8 (i.e. p<10^-8^).

We used the program *bedtools* to overlap each of the identified regions with previously annotated genes in the *X. birchmanni* PacBio^65^ assembly. See Dodge et al.^65^ for details on the approach used to annotate the *X. birchmanni* PacBio genome assembly (GenBank Assembly ID: GCA_036418095.1). These intervals and associated genes are reported in Table S3. Next, we used the Human MitoCarta3.0 database (https://www.broadinstitute.org/mitocarta/) to determine whether the focal gene is involved in mitochondrial pathways or localizes to the mitochondria. We used blastp to identify genes in the Human MitoCarta 3.0 database with a high match to *X. birchmanni* predicted protein sequences, using an e-value threshold of 10^-20^ (Supporting Information 2). Note that due to the teleost whole genome duplication, some MitoCarta genes matched two *X. birchmanni* protein sequences (typically delineated as *a* and *b* in the *X. birchmanni* genome annotation). We treated any identified MitoCarta genes as likely candidates for mitonuclear incompatibilities involving the focal region, given that nuclear-encoded genes involved in mitochondrial function are most likely to be involved in mitonuclear incompatibilities ^66, 38^.

To evaluate whether more MitoCarta genes overlapped with our admixture mapping intervals than expected by chance, or whether a greater proportion of peaks contained more than one MitoCarta gene compared to null expectations, we calculated the size of each of the admixture mapping intervals. We then performed permutations by randomly selecting a chromosome and drawing a start position from a uniform distribution ranging from 1 to the chromosome end. We defined the permuted admixture mapping interval for that peak by taking the random start interval and adding the length of the interval being simulated to generate the stop position. Once we had simulated locations for all the admixture mapping peaks in our dataset, we overlapped these peaks with the locations of MitoCarta genes and counted the total number of overlapping genes and the number of peaks with at least one MitoCarta gene. We repeated this procedure 1,000 times and compared the simulated and observed data.

### Segregation distortion analysis in a large dataset of F_2_ hybrids harboring the X. malinche mitochondria

In previous work, some of the first evidence we detected for mitonuclear incompatibilities was based on signals of segregation distortion in ∼950 F_2_ hybrids raised in common garden conditions^25^. We have now collected data from 1748 F_2_ hybrids, giving us increased power to detect segregation distortion along the genome. However, it is important to note that due to lower success of one cross direction^67^, we are only able to generate F_2_ hybrids with the *X. malinche* mitochondria, and thus we do not expect to see segregation distortion surrounding regions associated with the *X. birchmanni* mitochondria (e.g. interactions listed as “*X. birchmanni* incompatible” in Table 1).

To set the significance threshold for segregation distortion analysis, we performed neutral simulations of 1,450 F_2_ hybrids using the program admix’em^68^ (>99% of markers were covered in ≥1450 individuals in our empirical dataset). We simulated 24,000 markers spread across 24 chromosomes, matched in size to the 24 *Xiphophorus* chromosomes. Admix’em can take advantage of user-provided local recombination rates. We calculated average recombination rates from the *X. birchmanni* population recombination map^55^ in 5 kb intervals and used this to specify recombination priors in admix’em, assuming that each chromosome experienced an average of one crossover per meiosis. Following each simulation, we calculated average ancestry at each ancestry informative site. We repeated this procedure 100 times and used the upper and lower 2.5% quantile from these simulations (46.2 and 53.4% parent 1 ancestry, respectively) as our significance threshold.

For the real data, we identified stretches of markers that fell above or below this threshold as potential segregation distorters (Fig. 4A). We excluded ancestry informative sites with fewer than 1450 individuals covered (∼0.1% of informative sites in our dataset). We also excluded regions where distortion extended for less than 100 kb, as such regions are unexpected in a dataset of early-generation hybrids where admixture linkage disequilibrium typically extends for many megabases.

**Fig. 4.**
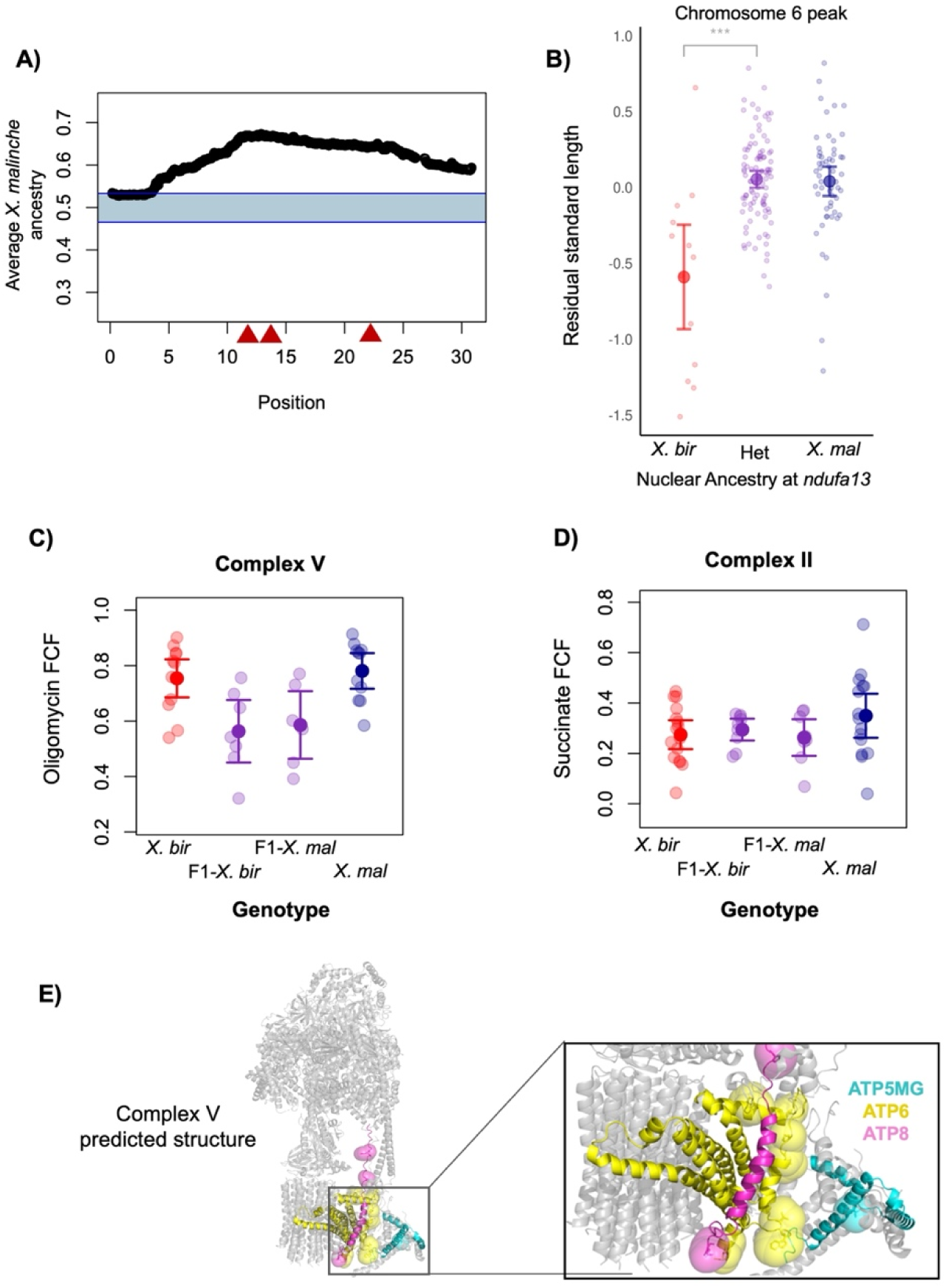
**A)** Segregation distortion along chromosome 6 for the 1748 F_2_ hybrids analyzed in this manuscript. Most of chromosome 6 departs from expectations under a scenario with no selection. Red triangles indicate the locations of admixture mapping peaks identified in the Calnali Low hybrid population (see Fig. 2). **B)** Lab raised hybrids with homozygous *X. birchmanni* ancestry on chromosome 6 and an *X. malinche* mitochondria are smaller (body size; residual standard length) on average than hybrids that are heterozygous or homozygous *X. malinche.* Large points and whiskers indicate the mean ± 2 standard errors, small points correspond to the raw data. The genotypes plotted here correspond to the admixture mapping peak at *ndufa13* (12.5 Mb), but a strong relationship between genotype and body size is observed for F_2_ hybrids across chromosome 6 (see Table S4). **C)** Complex V function is reduced in F_1_ hybrids with the *X. birchmanni* (F_1_ - *X. bir*) or *X. malinche* mitochondria (F_1_- *X. mal*) compared to pure *X. birchmanni* (*X. bir*) and *X. malinche* (*X. mal*). The Oligomycin flux control factor (Oligomycin FCF) represents the impact of inhibiting Complex V activity after stimulating Complexes I & II. Lower impact of inhibiting Complex V in hybrids indicates lower baseline activity in this protein complex, either caused directly by dysfunction in Complex V or by secondary effects of dysfunction in earlier components of the electron transport chain. **D)** Complex II function in pure *X. birchmanni* (*X. bir*), *X. malinche* (*X. mal*), and F_1_ hybrids with the *X. birchmanni* (F1 – *X. bir*) or *X. malinche* mitochondria (F1 – *X. mal)*. The Succinate flux control factor (Succinate FCF) represents the impact of activating Complex II with its substrate, succinate, after Complex I has been activated. Note that Complex II includes only nuclear-encoded proteins, so serves as a control comparison where we do not expect to observe mitonuclear incompatibilities. In **C & D**, semi-transparent points show individual data, point and whiskers show mean ± two standard errors. **E)** Predicted protein structure of Complex V which contains *ATP5MG* (cyan), *ATP6* (yellow), and *ATP8* (magenta). Large spheres represent amino-acids that differ between *X. birchmanni* and *X. malinche*.

### Analysis of size variation by genotype in laboratory-generated hybrids

Due to the potential for mitochondrial incompatibilities to impact growth, we raised F_2_ fry in controlled conditions and measured their size after 3-6 months of age. Briefly, fry were separated from parents less than one week after they were born and raised in common aquarium conditions. We tracked a total of 181 individuals from seven families. Once individuals were large enough to be individually tagged, they were marked with a unique elastomer fluorescent tag, fin clipped for genotyping, and photographed on a standard background with a ruler. Images were analyzed using the Fiji software^69^, and standard length measurements (length of the fish from the snout to the beginning of the caudal fin rays) were collected for each tagged fish. Subsequently, we performed ancestry inference as described above and selected an ancestry informative marker that tagged each region of interest on chromosome 6 (11.53-13.5 Mb and 20.25 Mb), chromosome 4 (6.84-7.14 Mb), chromosome 13 (2.1 Mb), and chromosome 15 (17.37-17.61 Mb). We then analyzed the data using a Linear Mixed Model in R, evaluating the relationship between length and genotype, including family/tank as a random variable (Fig. 4B). We implemented a Bonferroni correction to adjust p-values for multiple tests (Table S4). We also re-analyzed a dataset of morphological and physiological data from 235 F_2_ embryos^25^ as a function of genotype at the newly identified incompatibilities (Supporting Information 3).

### Analysis of mitochondrial function by genotype in laboratory-generated hybrids

Given that mapping results implicated multiple mitochondrial protein complexes in incompatible interactions, we were interested in directly measuring the performance of different components of the mitochondrial electron transport chain in hybrids and parent species. However, since several of the incompatible interactions are lethal or nearly lethal in the homozygous state, we decided to evaluate this question in F_1_ hybrids. We compared mitochondrial performance in F_1_ hybrids with either the *X. birchmanni* or *X. malinche* mitochondrial haplotype to both parental species (Fig. 4C-D). In previous studies, we were not able to assay the *X. birchmanni* mitochondria in hybrids. *Xiphophorus* species are live-bearing fish and *X. birchmanni* mothers carrying F_1_ embryos have a high rate of spontaneous abortion and maternal mortality^25^. However, some offspring are occasionally viable from this cross, and by scaling up the number of crosses attempted, we were able to generate sufficient F_1_ hybrids for physiological assays of this cross direction.

Our protocol for mitochondrial respiration measurements was identical to that described in a previous publication^25^, including preparation and isolation of the mitochondria by differential centrifugation, and titration of the mitochondria in Mir05 solution in the Oroboros O2K with the same substrates and inhibitors in the same order. The only deviation from the prior protocol was that, rather than standardizing all runs to the same quantity of mitochondrial protein, we allowed runs to vary in this quantity because this allowed us to include more samples for each genotype. Note that this does not impact the calculations of flux control factors, which are internally controlled for mitochondrial protein content. Details on the calculation of flux control factors can be found in Moran et al.^25^.

### Protein modeling

Based on our admixture mapping results, we were interested in determining the locations of nonsynonymous substitutions between species in mitochondrial Complex V (also known as the ATP synthase complex). To predict the structure of individual proteins and the overall structure of Complex V surrounding mapped mitonuclear interactions (Table 1), we loaded the protein sequences into ColabFold v1.5.5: AlphaFold2 using MMseqs2^70^ and ran the software with its default parameters. We then visualized the ColabFold protein data bank formatted structures in PyMOL and used clustal omega^61^ to identify the position of nonsynonymous substitutions in the protein sequences. We visualized the predicted Complex V structure along with nonsynonymous substitutions in *ATP8*, *ATP6*, and *ATP5MG* using PyMOL (Fig. 4E).

### Analysis of substitutions and evolutionary rates in candidate nuclear genes involved in mitonuclear incompatibilities

For all proteins of interest associated with mitonuclear incompatibilities (Table 1), we calculated rates of protein evolution between *X. birchmanni* and *X. malinche*. We extracted predicted cDNA sequences using the genome annotations of each species and aligned them to ensure they were of equivalent length. We then used the codeml function in PAML^71^ to estimate the rate of nonsynonymous substitutions per nonsynonymous site (dN) versus the rate of synonymous substitutions per synonymous site (dS) and their ratio with the codeml model option set to zero. We also used predicted amino acid sequences of *X. birchmanni, X. malinche*, and two outgroups (*X. variatus* and *X. cortezi*), to predict which lineage each amino acid substitution arose in.

For each gene with nonsynonymous substitutions, we also extracted the predicted protein sequence from all bony fishes on NCBI, aligned them with clustal omega^72^, and visually inspected alignments for errors. We removed *X. birchmanni* and *X. malinche* from this analysis but included other *Xiphophorus* species with available sequences (*Xiphophorus helleri* and *X. couchianus*). We then used SIFT^73^ to evaluate whether the substitutions observed in *X. birchmanni* or *X. malinche* were predicted to have functional effects (i.e. predicted not tolerated).

### Approximate Bayesian Computation simulations with SELAM

To estimate the strength of selection consistent with mitonuclear incompatibilities identified in our admixture mapping data, we used an approximate Bayesian computation (ABC) approach, with the forward time simulator SELAM^74^. We performed SELAM simulations jointly modeling population history and selection on mitonuclear interactions, focusing on incompatibilities that have not been previously studied^25^ and are not physically linked to other interactions (since we found jointly simulating multiple linked incompatibilities computationally intractable). Detailed methods on ABC simulations in SELAM can be found in Supporting Information 4.

### Local ancestry in natural populations

To evaluate whether mapped incompatibilities are experiencing selection in natural hybrid populations, we drew on previously published datasets for naturally occurring *X. birchmanni* x *X. malinche* hybrids. We focused on populations that occur in independent river systems from each other and from the admixture mapping population^23^ (Fig. 1B; Table S2). One of these populations, the Tlatemaco population (n=96), has fixed the *X. malinche* mitochondrial haplotype, and the second, Acuapa (n=97), has fixed the *X. birchmanni* mitochondrial haplotype. We note that other majority-*X. birchmanni* and majority *X. malinche* populations exist but are not in independent river systems from populations used in other analyses, so we do not analyze them here. We asked whether regions of low minor parent ancestry (i.e. non-mitochondrial parent ancestry) in these populations coincided with the mapped locations of mitonuclear incompatibilities. We calculated average ancestry in 10 kb windows and compared windows that overlapped with mapped mitonuclear incompatibilities to the genome-wide average (Fig. 5A). To determine if mitonuclear incompatibilities on average had lower minor parent ancestry than expected, we performed simulations randomly drawing 10 kb windows from the genome-wide distribution and calculating average minor parent ancestry to generate a null distribution which we compared to the observed data.

**Fig. 5.**
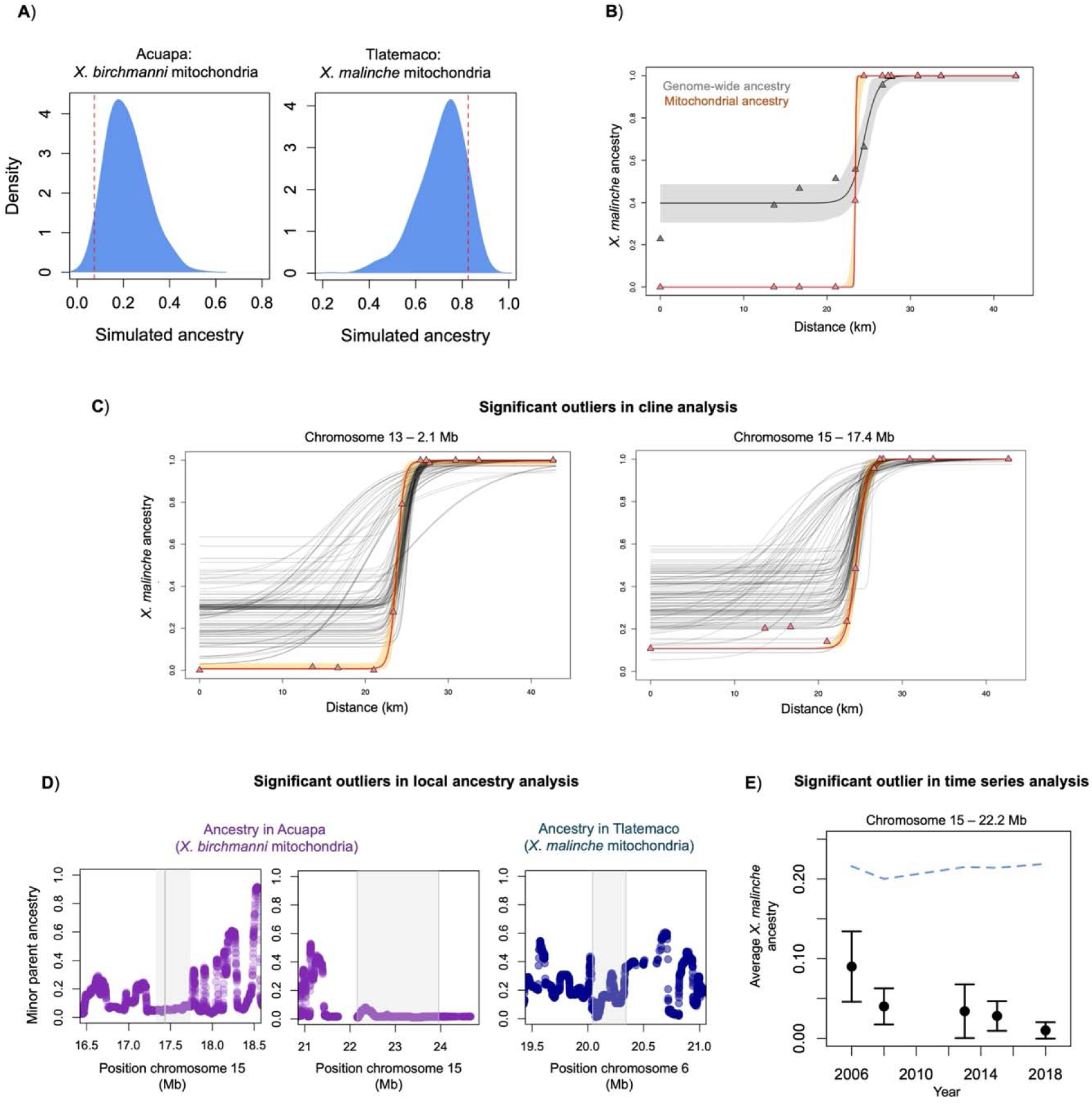
Evidence of selection on a subset of newly mapped mitonuclear incompatibility loci based on analyses of data from natural hybrid populations. **A**) Red dashed line indicates the average *X. malinche* ancestry at all newly mapped mitonuclear incompatibility loci compared to null simulations (blue distribution) in the Acuapa (left) and Tlatemaco (right) populations. The *X. birchmanni* mitochondrial haplotype is fixed in the Acuapa population and the *X. malinche* mitochondrial haplotype is fixed in the Tlatemaco population. Loci identified as mitonuclear incompatibilities had significantly less non-mitochondrial parent ancestry than the genome-wide background in the Acuapa population (see Methods; p=0.029 by simulation), but not in the Tlatemaco population (p=0.11 by simulation). **B)** Results of cline analysis considering average genome-wide ancestry (gray, cline center = 24.4, width = 3.26, *p_min_ =* 0.397, *p_max_ =* 0.999) and mitochondrial ancestry (orange, center = 23.4, width = 0.211, *p_min_ =* 0, *p_max_ =* 1) in populations from the Río Pochula. As expected, each individual had identical ancestry calls across ancestry informative markers in the mitochondria, so an arbitrary marker (10782 bp) was used for cline analysis. Triangles represent average ancestry in each population, line represents model fit inferred using the HZAR software^70^, and the envelope represents the 95% credible regions of the HZAR fit. **C)** Clinal changes in allele frequency across sampling sites in the Río Pochula at focal genomic regions on chromosome 13 (2.1 Mb) and chromosome 15 (17.4 Mb) versus 100 matched null markers (see Methods). SNP-specific clines were fit using the SNP closest to the admixture mapping peak (2,112,372 bp for chromosome 13, 17,383,659 bp for chromosome 15). Cline models were fit using the HZAR software. The red line represents the model fit by HZAR to each focal locus, triangles show average *X. malinche* ancestry in each sampled population, orange ribbons show 95% credible regions for the focal cline, and gray lines show null clines from matched control markers (see Methods). For cline results at other loci that did not significantly differ from their matched nulls, see Fig. S23. **D**) Local ancestry in Acuapa (purple) and Tlatemaco populations (blue) near a subset of admixture mapping peaks. Shown here are three cases where minor parent ancestry is especially low near the mapped mitonuclear interactions. Gray envelope indicates the associated region from admixture mapping. Results for all loci identified via admixture mapping can be found in Fig. S17. **E)** Change in minor parent ancestry over time at the chromosome 15 admixture mapping peak at 22.2 Mb in the Acuapa population. Points show mean ancestry at the focal region in each year and whiskers show ± 2 standard errors. Dashed blue line shows average minor parent ancestry genome wide in Acuapa over the same time period. For time series data for other mapped loci, which did not change significantly over our sampling period, see Fig. S18.

### Time series analysis

For one *X. birchmanni* x *X. malinche* hybrid population, the Acuapa population, we had access to samples from 2006, 2008, 2013, 2015, and 2018^75^. This provides a means to evaluate whether ancestry at incompatible loci has changed in frequency over time in this population. While we analyze all loci for completeness, we only expect a subset of these analyses to be informative. This is because our most recent samples from the Acuapa population are fixed for the *X. birchmanni* mitochondrial haplotype. Thus, we expect that analyses of incompatibilities involving the *X. birchmanni* mitochondria (i.e. “*X. birchmanni* incompatible” in Table 1) are most likely to result in a detectable signal.

For each interaction, we identified the peak ancestry informative site from our admixture mapping results (Table 1). We then intersected this peak site with our time series data from the Acuapa population, with average ancestry summarized in 10 kb windows. This resulted in an estimate for the change in *X. malinche* ancestry over time at each locus (Fig. 5E). We note that because this analysis involves the same population and some of the same samples, it is not independent of the local ancestry analysis in the Acuapa population discussed above.

### Cline analysis

As a complementary approach to investigate the role that mitonuclear incompatibilities play as barriers to gene flow between *X. birchmanni* and *X. malinche* populations in nature, we used a clinal dataset collected from the Río Pochula (Fig. 1D). This dataset spans 12 populations and an elevation gradient from ∼200 meters to 1,400 meters (Table S1; Fig. 5). Ancestry inference was performed on these datasets as described above. We next identified and removed markers that were not present in all populations across the river, leading to a total of 331,518 ancestry informative sites. We selected the marker closest to the peak signal in our admixture mapping analysis. For all regions, we were able to select a marker that fell within the focal admixture mapping region. We also calculated the average genome-wide ancestry of individuals in each population, allowing us to generate a genome-wide cline for comparison to clines at loci involved in mitonuclear incompatibilities.

To evaluate the significance of observed patterns at each locus, we first generated null datasets matching for local gene density and local recombination rate. We summarized the number of coding basepairs in 100 kb windows along the genome, as well as at the recombination rate, estimated from *X. birchmanni*^24,55^, at this same spatial scale. Next, for each focal region, we selected windows that fell within ± 20% the number of coding basepairs and inferred recombination rate. We identified all ancestry informative markers in those matched windows and randomly selected 100 markers as control markers for the focal region of interest.

To fit cline models to the focal and matched control datasets, we used the HZAR^76^ software to identify the best-fit model from two alternatives, either with cline maxima and minima as free parameters to be estimated (scaling = “free”, tails = “none” in hzar.makeCline1DFreq function) or fixed at the observed maximum and minimum allele frequencies in the data at that ancestry informative site (scaling = “fixed”, tails = “none”). From the best fit HZAR model, we extracted estimates of cline width, center, and minimum *X. malinche* ancestry, and compared these values between the focal and matched control datasets and to the genome-wide average.

## Results

### Admixture mapping reveals new mitonuclear incompatibilities

Previous analyses using smaller datasets detected three mitonuclear incompatibilities (Table 1), two of which could be traced to specific nuclear genes^25^. Here, we used a large admixture mapping population of natural hybrids (N=731) from the Calnali Low hybrid population (Fig. 1D), to map additional mitonuclear hybrid incompatibilities. In addition to confirming patterns at previously detected incompatible interactions involving *ndufa13* and *ndufs5* (Fig. S1), we identified new interactions (Fig. 2A). Specifically, the peaks on chromosome 4, chromosome 6 at 13.3 Mb, chromosome 6 at 20.3 Mb, chromosome 11 (but see below), chromosome 15 at 17.5 Mb, chromosome 15 at 19.6 Mb and chromosome 16 were not identified in our previous work^25^. The detection of new interactions is likely facilitated by our larger dataset, which more than doubles the sample size of our first study^25^, increasing power to detect genotype combinations that are underrepresented in the admixture mapping population. As expected from models of selection against hybrid incompatibilities, most regions that exceed our genome-wide significance threshold show an increased rate of matched ancestry between the mitochondrial and nuclear genome (signals on chromosomes 4, 6, 13, 15 and 16). One exception is a signal on chromosome 11, which is enriched for ancestry mismatch between the mitochondrial and nuclear genome. Since this is not a signal predicted by models of hybrid incompatibilities, we omit chromosome 11 from our analyses in the main text but discuss these results in the supplement (Supporting Information 5; Fig. S2).

Chromosomes 6 and 15 exhibit multiple distinct peaks of association with mitochondrial ancestry (Fig. 2C-D). Examining alignments of the two species generated from long read assemblies indicated that there were no small or large structural rearrangements that might be generating unusual patterns of associations in these regions (Fig. S3). We find that between each pair of peaks on the same chromosome, admixture linkage disequilibrium decays to background levels, suggesting that they are indeed distinct signals (Fig. S4; Supporting Information 6). We also confirmed that patterns of missing data and local ancestry are not expected to generate multiple peaks from a single signal (e.g. due to reduced power from locally high rates of missing data; Supporting Information 6). Together, this supports the conclusion that there are multiple mitonuclear incompatibilities on both chromosome 6 and chromosome 15. Throughout the rest of the paper, we refer to these loci by their chromosome and nuclear position, which are summarized in Table 1.

### Diverse incompatibility architecture revealed by admixture mapping

To investigate evidence for selection on particular ancestry combinations in the mitochondrial and nuclear genomes of hybrids, we compared observed genotype combinations to those expected by chance in our admixture mapping population. To appropriately account for variation in admixture proportion in the population, we used a simulation-based approach (see Supporting Information 1). We found that the majority of detected incompatibilities only involved one mitochondrial haplotype. For example, focusing on the interaction on chromosome 6 at 20.3 Mb, we see that individuals with homozygous *X. birchmanni* ancestry at the admixture mapping peak and an *X. malinche* mitochondrial haplotype are depleted from our dataset (Fig. 3B). The alternative genotype combination (homozygous *X. malinche* ancestry with an *X. birchmanni* mitochondrial haplotype) is not significantly underrepresented compared to null expectations at this region on chromosome 6. However, we do identify three cases where selection on mitonuclear ancestry mismatch appears to be bidirectional, including one previously reported case on chromosome 13^25^ (Fig. S1) and previously undetected loci on chromosome 4 and chromosome 15 (Table 1; Fig. 3C-3D). Notably, we infer that the distinct admixture mapping peaks on chromosome 15 have different genotypes under selection (Fig. 3C-3D; Table 1). For simplicity, throughout the manuscript we describe these various interactions as regions incompatible with the *X. malinche* mitochondria (“*X. malinche* incompatible”), incompatible with the *X. birchmanni* mitochondria (“*X. birchmanni* incompatible”), or as bidirectional incompatibilities (Fig. 3A). For a summary of these results, see Table 1. Overall, our identification of several bidirectional incompatibilities highlights that a subset of mitonuclear incompatibilities between *X. malinche* and *X. birchmanni* experience more complex selection than predicted by classic hybrid incompatibility models^10,15^.

### Identification, evolution, and expression of mitonuclear genes shows varied patterns

For each of the identified regions, we determined whether any of the annotated genes in these regions were present in the MitoCarta 3.0 database of known nuclear-encoded genes in mammals that localize to the mitochondria. We identified at least one MitoCarta annotated gene in all three chromosome 6 intervals (one previously reported^25^), on chromosome 13 (as previously reported^25^), on two of the three intervals on chromosome 15, and the chromosome 16 interval. Four of the newly mapped intervals had only a single MitoCarta annotation, which likely represents the causal gene of the mitonuclear interaction in these regions (chromosome 6 at 13 Mb – *mterf4*, chromosome 6 at 20.3 Mb – *atp5mg*, chromosome 16 – *uqcrc2*, and chromosome 15 at 17 Mb – *mmut*). A summary of these results can be found in Table 1. Compared to null expectations generated by permuting the admixture mapping intervals on the genome, there was no significant enrichment in the number of MitoCarta genes falling within these peaks or the number of peaks with at least one MitoCarta gene (p=0.83 and p=0.1 respectively, by simulation).

For each identified MitoCarta gene, we compared the predicted amino acid sequence between *X. birchmanni* and *X. malinche* and estimated *d*_N_/*d*_S_ between species (Table 1). We found that none of the newly identified MitoCarta genes from admixture mapping were rapidly evolving between species (Table 1). Six of the eight newly identified genes had an estimated *d*_N_/*d*_S_ <1. The remaining two genes, *atp5mg* and *rmdm3*, had nonysynonymous substitutions but lacked synonymous substitutions, resulting in undefined *d*_N_/*d*_S_ estimates. To evaluate these genes further, we compared the number of nonsynonymous substitutions in these genes to an empirical distribution of length-matched genes genome-wide. Compared to genes within 10% of their length, neither *atp5mg* nor *rmdm3* were significant outliers in the number of nonsynonymous substitutions (p=0.11 and p=0.2 by simulation respectively). By contrast, previously mapped mitonuclear incompatibilities in this system show strongly elevated rates of amino acid evolution^25^. We also queried previously collected RNAseq data^77^ to infer whether any of the genes of interest were differentially expressed between the parent species, but found limited evidence for this (Table 1, Fig. S5).

Examining the patterns of substitutions between *X. birchmanni* and *X. malinche* relative to two outgroups at each newly identified gene, we found that substitutions distinguishing species did not appear to be derived on a particular lineage (Fig. S6). This pattern is notably distinct from previous results for the mitonuclear incompatibilities involving *ndufs5* and *ndufa13,* where nonsynonymous substitutions had accumulated disproportionately on the *X. birchmanni* branch^25^. We also evaluated whether any of the observed substitutions between *X. birchmanni* and *X. malinche* involved changes that are likely to impact protein function using SIFT (Table 1). Notably, we detected several such substitutions (including those previously detected in *ndufs5* and *ndufa13*), two of which fell in mitochondrial “leader” sequences, which are short signal peptides that target the localization of the protein to the mitochondria (Table 1; Fig. S6). This indicates the presence of substitutions that are likely to alter protein function or localization within the admixture mapping regions.

### Structural modeling suggests direct physical interactions do not explain newly identified mitonuclear incompatibilities

One mechanism through which hybrid incompatibilities can arise is through a breakdown in protein-protein (or protein-DNA/RNA) interactions (see also^5,78,79^). Since protein complexes involved in mitochondrial function have been intensively studied, we identified “chimeric” protein complexes, where nuclear-encoded proteins formed larger complexes that include mitochondrial-encoded proteins (or RNAs), as particularly likely sites of mitonuclear interactions. We focused on protein complexes that had structures in the RCSB PDB database (https://www.rcsb.org/). Based on these criteria, we investigated *mterf4* and *atp5mg* in more detail. Note that the reference database structures used are not specific to *Xiphophorus* but represent solved structures for other eukaryotic species that are highly conserved over deep evolutionary distances (Fig. S7). See Moran et al.^25^ for our previous structural analyses of *ndufs5* and *ndufa13*.

One identified gene, *mterf4,* encodes a protein that interacts with mitochondrial-encoded rRNAs. We found that all nonsynonymous differences between *X. birchmanni* and *X. malinche* in this gene fell in the protein “leader” sequence. This highly conserved sequence is subsequently cleaved and thus is unlikely to impact physical interactions between *mterf4* and mitochondrial-encoded rRNAs. As a result, we did not model this protein-RNA interaction further.

Another gene, *atp5mg*, encodes one of the nuclear accessory subunits of the chimeric OXPHOS Complex V (ATP synthase) which catalyzes ATP synthesis across the mitochondrial inner membrane. We found that two of the nonsynonymous differences between *X. malinche* and *X. birchmanni* also fell in the leader sequence of this protein. After removing the leader sequence, one nonsynonymous substitution between *X. birchmanni* and *X. malinche* remained in *atp5mg*. Our modeling results indicate that *ATP5MG* is in physical contact with both mitochondrial-encoded ATP synthase proteins (*ATP6* and *ATP8*; Fig. 4E), but the nonsynonymous substitution itself is not. Due to the predicted distance [>25 Å] between nonsynonymous substitutions in *ATP5MG, ATP6,* and *ATP8*, we consider it unlikely that that there are direct physical interactions between *X. malinche* and *X. birchmanni* substitutions in these proteins. This finding contrasts with previous findings for *NDUFS5* and *NDUFA13* where multiple species-specific substitutions were predicted to be in contact between mitochondrial and nuclear proteins in Complex I^25^. However, indirect interactions could impact the function of Complex V as a whole. This possibility is especially intriguing given the proximity of these proteins to the flow of protons across the mitochondrial membrane and the c-ring rotor^80^. We compare the performance of Complex V in hybrids and the parental species below.

### Evidence for segregation distortion in lab hybrids harboring X. malinche mitochondria

All lab-generated F_2_ hybrids have the *X. malinche* mitochondria; the alternative cross has a low success rate in lab due to a high abortion and maternal mortality rate^67^. Four of the newly identified interactions involve the *X. malinche* mitochondria or are inferred to be bidirectional (Fig. 3). This suggests that ancestry distortions at these loci should be detectable in lab raised hybrids, assuming that the incompatibilities are not driven solely by sources of selection that are environmentally dependent. Our previous work using a smaller number of F_2_ hybrids showed evidence of colocalization of segregation distortion and the *ndufa13* and *ndufs5* incompatibilities^25^. Here, we use a dataset of 1748 F_2_ hybrids to revisit these trends (Table S2).

We find evidence of significant segregation distortion overlapping all of the chromosome 6 admixture mapping peaks. Specifically, we see a strong depletion of *X. birchmanni* ancestry across the majority of this chromosome (Fig. 4A). We detect similar patterns on chromosome 13 with our larger dataset (Fig. S8) as we reported previously^25^. While we detect weak but significant signals of segregation distortion on chromosomes 4 and 15, these either do not localize with the admixture mapping peaks or show unexpected directionality (Fig. S9). This could indicate that individuals with these genotype combinations are not under strong selection in a lab environment. However, these results may also be explained by low power to detect segregation distortion. Simulations suggest that even with our large sample size we may lack power to identify segregation distortion in cases where selection coefficients fall below 0.3 (Supporting Information 7).

### Incompatibilities are associated with size variation in lab-raised F_2_s

For a subset of 181 F_2_ individuals with the *X. malinche* mitochondria, we were able to raise individuals in the lab for several months and collected paired genotype and phenotype data. Given the expected importance of the mitochondria in growth^81^, we analyzed the correlation between genotype at admixture mapping peaks and standard length within groups of siblings. Specifically, we analyzed ancestry at mapped incompatibilities involving the *X. malinche* mitochondria on chromosome 4 (6.84-7.14 Mb), chromosome 6 (11.53-13.6 Mb, and 20.25 Mb), chromosome 13 (2.10-2.14 Mb) and chromosome 15 (17.37-17.61 Mb).

We found strong correlations between genotype on chromosome 6 and standard length (measured as the length from snout to caudal peduncle) in families of lab-raised F_2_ individuals. Individuals with incompatible genotypes were significantly smaller than their siblings with compatible genotypes (Fig 4B). The mean length of incompatible individuals was 2.83 cm, while individuals with heterozygous and homozygous *X. malinche* ancestry were on average larger by 0.603 cm and 0.623 cm, respectively (Table S4). Note that due to strong linkage among the three loci on chromosome 6 in early generation hybrids (Supporting Information 6), we cannot distinguish which of the three loci is driving the observed body size phenotype. We did not identify associations between standard length and genotype for loci on chromosomes 4, 13, or 15.

We previously collected respirometry and morphometric data from 235 lab-generated F_2_ embryos, which we reanalyze here, again focusing on incompatibilities involving the *X. malinche* mitochondria. In addition to effects of loci on chromosome 6 and 13 on rates of embryonic respiration and morphological defects^25^ (see Supporting Information 3; Fig. S10), we found that homozygous *X. birchmanni* ancestry at the newly identified peak on chromosome 4 contributed to smaller head width in embryos (F=7.3, p=0.0075).

### Physiological signatures of mitonuclear hybrid incompatibilities

While we do not have access to lab-generated F_2_ adults with both mitochondrial types, we were able to generate F_1_ hybrids with either the *X. birchmanni* or *X. malinche* mitochondria (see Methods). This allowed us to directly compare different features of mitochondrial function in F_1_ hybrids with either the *X. birchmanni* or *X. malinche* mitochondria to that of the parental species. We note that while F_1_ hybrids do not appear to experience reduced viability as a result of mitonuclear incompatibilities, they may still show physiological signatures of subfunctional mitochondria. In a prior study, we demonstrated that F_1_ hybrids have reduced function in the chimeric mitochondrial Complex I (NADH dehydrogenase)^25^, but not in overall mitochondrial function, demonstrating a negative effect of the incorporation of incompatible proteins that may be compensated for by other mitochondrial processes. A caveat of that previous work was that we were only able to assess function in F_1_ hybrids with the *X. malinche* mitochondria.

Here, we used F_1_ adult hybrids with both mitochondrial haplotypes to demonstrate that hybrids of both types show evidence of bidirectional dysfunction in the three chimeric mitochondrial complexes that we investigated (Complexes I, IV, and V), based on a substrate-uncoupler-inhibitor-titration protocol using the Oroboros O2K high-resolution respirometry system^25^ (Fig. 4C-D; Fig. S11-S13). We confirmed that hybrids of both types show reduced Complex I activity relative to parentals, quantified both by the activation of Complex I with ADP (ANOVA; *d.f.* = 3, *F* = 6.1122, *p* = 0.0015; Fig. S11) and its inhibition by rotenone (*d.f.* = 3, *F* = 8.2008, *p* < 0.0005; Fig. S11). This bidirectional dysfunction of Complex I is expected given that both mitochondrial types have mapped incompatible interactions with Complex I proteins. Note that the response to rotenone we detect here is stronger than detected in our previous study^25^. Both hybrid types also showed a reduced response to the inhibition of Complex V by oligomycin (*d.f.* = 3, *F* = 6.5512, *p* = 0.0014; Fig. 4), and the activity of Complex IV (cytochrome-c oxidase) as suggested by the response to ascorbic acid and TMPD (*d.f.* = 3, *F* = 5.4665, *p* = 0.0041; Fig. S12).

By contrast, for Complex II—which is often viewed as a control when investigating mitonuclear interactions since it is entirely encoded by the nuclear genome—hybrids show little evidence of reduced function (Fig. 4). The activity of Complex II (succinate dehydrogenase) when stimulated by succinate did not differ among hybrids and parentals (*d.f.* = 3, *F* = 1.2224, *p* = 0.3128; Fig. 4D). We note, however, that the inactivation of Complex II by malonate did differ between hybrids and parentals (*d.f.* = 3, *F* = 5.1205, *p* = 0.0056; Fig. S13). We saw no significant differences in responses between the two hybrid types, or between the two parental species, in any comparison (Table S5).

Together, these results suggest a clear impact of mitonuclear incompatibilities on the activation of chimeric complexes. Notably, the incompatibility at 20.3 Mb on chromosome 6, which contains *ATP5MG*, could be causing Complex V dysfunction, although the bidirectional dysfunction we observed could be attributable to incompatibilities in upstream complexes as well. The drivers of dysfunction in Complex IV are not yet evident from our mapping results, but may also result from upstream Complex I incompatibilities. Due to the design of our assay, we were not able to investigate Complex III activity, but we may expect to see some reduction in Complex III function in hybrids given that the mapped mitonuclear interaction on chromosome 16 contains *uqcrc2* (Table 1), a nuclear subunit of this chimeric complex.

### Strong selection against mitonuclear incompatibilities

Previous mapping results for mitonuclear incompatibilities involving *ndufa13* and *ndufs5* indicated that mismatched ancestry between the *X. malinche* mitochondria and *X. birchmanni* ancestry at these genes was essentially lethal, with estimated selection coefficients for ancestry mismatch at both loci exceeding 0.9^25^. Given that the additional mitonuclear incompatibilities we map here were not detectable with our previous admixture mapping population (N=359), we expect *a priori* that these newly identified mitonuclear incompatibilities should be under weaker selection.

We tested this prediction using an approximate Bayesian computation approach implemented in SELAM (see Methods^74^) to estimate the strength of selection acting on incompatibilities involving loci on chromosome 4 and 16. We recovered well-resolved posterior distributions for selection and dominance coefficients for all four of the modelled incompatibilities (Fig. S14-S15). As expected, our results are consistent with weaker but still severe selection on the mitonuclear interactions (maximum *a posteriori* estimates of *s* = 0.59 and 0.72 for chromosome 4 mismatched with the *malinche* or *birchmanni* mtDNA respectively, and *s* = 0.67 and 0.75 for chromosome 16). The credible intervals for these incompatibilities were between *s* = 0.22–0.98 and *s* = 0.27-0.97 for chromosome 4, and *s* = 0.09 – 0.97 and *s* = 0.22 – 0.99 for chromosome 16, in keeping with the results of our power analysis (Fig. S14 & S15; Supporting Information 8). Three of the four interactions were inferred to be partially recessive (Fig. S14 & S15).

### Cline analysis indicates selection acting on some incompatibilities in natural populations

To evaluate evidence for selection on mitonuclear incompatibilities in nature, we analyzed clinal ancestry patterns in the Río Pochula, where we had access to samples spanning 12 sites along the river. These sites ranged from *X. birchmanni-*typical elevations of ∼200 meters to *X. malinche-*typical elevations of up to 1400 meters. Compared to matched control loci, three mapped regions were significant outliers based on either the minimum *X. malinche* allele frequency or cline width (Table 1). Each of the incompatibilities that were identified as cline outliers were either under bidirectional selection or involve only the *X. birchmanni* mitochondria (Table 1; Fig. 3). Because of the structure of migration in *X. birchmanni* x *X. malinche* hybrid zones, we expect migration to predominantly occur from upstream *X. malinche* populations to downstream *X. birchmanni* populations (Fig. 1B). Thus, it is not surprising that incompatibilities involving the *X. birchmanni* mitochondrial haplotype are the most detectable using cline approaches (Supporting Information 9; Fig. S16). We note that three interactions involving the *X. birchmanni* mitochondria were not significant outliers based on cline analysis, which could reflect a lack of power to detect these interactions or that selection on them is context-dependent.

### Patterns of local ancestry at incompatibility loci in hybrid populations

As a complementary approach to study selection in natural populations, we turned to previously collected data from two natural hybrid populations that formed ∼100 generations before the time of sampling^23,57^. These populations occur in different river systems from both the Río Pochula populations and from the admixture mapping population (Fig. 1B) and thus can be viewed as independent datasets for studying selection on mitonuclear incompatibilities in nature.

For incompatibilities involving the *X. birchmanni* mitochondria, we re-analyzed population genomic data collected from 97 hybrids from the Acuapa population^25,57^, which has fixed for the *X. birchmanni* mitochondria (Fig. 1B). For incompatibilities involving the *X. malinche* mitochondria, we re-analyzed genomic data collected from 96 hybrids from the Tlatemaco population, which has fixed the *X. malinche* mitochondria (Fig. 1B). Overall, we found evidence that minor parent ancestry (i.e. non-mitochondrial parent ancestry) was less common than expected in regions surrounding mapped mitonuclear incompatibilities (Fig. 5). However, we find moderate levels of minor parent ancestry surrounding some incompatibilities (Fig. S17). This could again indicate that selection on some of the mitonuclear incompatibilities is context dependent or that selection is too weak or variable to drive effective purging of incompatibilities in all natural populations.

We also took advantage of time series data previously collected from the Acuapa population that spans approximately 25 generations of evolution in the hybrid population. Since the Acuapa population is estimated to have formed approximately 100 generations before the present^23,57^, we may expect incompatibilities that are under strong selection to have already been purged by the time sampling began (e.g. *ndufs5*; see^57^). Here we focus on interactions involving the *X. birchmanni* mitochondria, since this mitochondrial haplotype is fixed in present-day samples from Acuapa. However, we report results for all loci in Fig. S18.

Of the three regions for which we expect to see directional selection against *X. malinche* ancestry in the nuclear genome in the Acuapa population (Table 1; chromosome 15 at 17.4 Mb, chromosome 15 at 22.2 Mb and chromosome 16 at 12.8 Mb), we see evidence for a change in ancestry over time in one region (Fig. 5; chromosome 15 at 22.2 Mb), although the relationship is marginally significant (t= −3.2, p=0.048). However, all three regions have relatively low *X. malinche* ancestry at the start of sampling and maintain this pattern over time (Fig. S18), consistent with selection in prior generations impacting the starting *X. malinche* ancestry frequency in our samples from 2006.

## Discussion

The genetic architecture of reproductive isolation between closely related species is a foundational question in evolutionary biology and is intrinsically linked to the question of how new species arise. While models of how hybrid incompatibilities may evolve were proposed nearly a century ago^1,2^, empirically identifying hybrid incompatibilities, studying their genetic architecture, and their impacts on patterns of genetic exchange between species has been challenging. The technical difficulties of mapping incompatibilities, including low power of existing methods, requirements for large numbers of hybrids, and poor mapping resolution, have stymied empirical progress in this area. Here, we leverage natural hybrid populations between *X. birchmanni* and *X. malinche* to map six new mitonuclear incompatibilities, resulting in a total of nine mitonuclear incompatibilities identified in this system. We also build on our previous work^25^ by applying new approaches for studying selection on these regions, including cline analysis and time series analysis.

Our results reveal a complex landscape of selection on ancestry mismatch between the mitochondrial and nuclear genomes in *X. birchmanni* x *X. malinche* hybrids. Notably, simulations indicate that we only have power to detect mitonuclear incompatibilities under moderate to strong selection, hinting that the true number of mitonuclear incompatibilities that are physiologically relevant to hybrids may be even larger. Moreover, because we focus only on mapping interactions between the mitochondrial and nuclear genome, many more incompatibilities may exist between these species genome-wide (consistent with some previous work^27,56^). Since only ∼1500 genes are known to localize to and interact with the mitochondria^82^ we would predict *a priori* that nuclear-nuclear incompatibilities should be more common. However, it is also possible that the large number of mitonuclear incompatibilities is attributable to the unusual biology of the mitochondrial genome. In many vertebrates, the mitochondria experiences >10X higher nucleotide substitution rate relative to the nuclear genome^39^, and due to physical interactions between mitochondrial encoded and mitochondrially localizing proteins encoded in the nuclear genome, high rates of protein coevolution are common^66,38^. Consistent with these observations in other species, pairwise divergence between *X. birchmanni* and *X. malinche* in the nuclear genome is ∼0.4% whereas mitochondrial divergence is 4.9%. Other mechanisms, such as sexual conflict driven by uniparental inheritance^17^, could also be responsible for a “large-mitochondrial” effect in the evolution of hybrid incompatibilities. An exciting future direction is disentangling how the number of mitonuclear incompatibilities scales with genetic divergence between species and whether patterns inferred here generalize to nuclear-nuclear interactions.

The large number of newly mapped mitonuclear incompatibilities identified here allows us to investigate the architecture of selection on these interactions and begins to provide hints about the architecture of reproductive isolation more generally. First, we detect some incompatibilities that are “asymmetric,” meaning that only one mitochondrial type is incompatible with mismatched ancestry in the nuclear genome. These types of incompatibilities are those envisioned by classic models in evolutionary biology^1,2,13,15^ (see Supporting Information 10). However, we also identify several incompatibilities that are “symmetric” or bidirectional, meaning that selection acts against mismatched mitochondrial-nuclear ancestry in both directions. Our findings suggest that bidirectional incompatibilities could be relatively common (though we note that they are also easier to map; Supporting Information 8), and it may be useful to revisit classic models in light of these empirical results. More recently proposed models for the evolution of hybrid incompatibilities, such as coevolutionary models^83^ and developmental systems drift models may more readily explain the emergence of symmetrical incompatibilities^25,35^.

Theory predicts that the architecture of selection on genes involved in hybrid incompatibilities will impact their efficacy as barriers to genetic exchange between species^15,83^. Asymmetric incompatibilities can revert to a compatible ancestral genotype and thus can be ineffective at preventing gene flow between species^18^ and may even introgress between species^22^. However, bidirectional incompatibilities are blocked from reversion to an ancestral genotype by low fitness intermediates, and thus should be more effective barriers to introgression^83^. Regardless of architecture, very few empirical studies to date have evaluated the efficacy of known incompatibilities as barriers to gene flow in nature^21–24^. Thus, after mapping mitonuclear incompatibilities in one river system, we explored evidence of selection on these regions in independently formed hybrid populations in different rivers. On average, we find that loci inferred to be involved in mitonuclear incompatibilities have depleted non-mitochondrial parent ancestry in natural hybrid populations (Fig. 5), but this pattern is stronger around bidirectional incompatibilities. Two out of three of the loci that we infer to be under bidirectional selection are outliers in clinal analyses, and all three loci have low levels of minor parent ancestry in samples from natural hybrid populations (Fig. 5, S17). These results are consistent with theoretical and simulation studies which predict that bidirectional incompatibilities are more likely to resist gene flow between species^15,83,84^. We note, however, that there are several possible interpretations of variation in ancestry at other mitonuclear incompatibilities in natural populations (Table 1; Fig. S17), including environmental dependence of selection, which may be expected given the large role of the mitochondria in homeostasis and organismal physiology^85,86^.

Our admixture mapping approach allowed us to detect several linked hybrid incompatibilities co-occurring on the same chromosome. In early generation crosses, signals from linked incompatibilities may be obscured by an insufficient number of crossovers. In natural hybrid populations, many generations of recombination can unmask these interactions. Indeed, in our previous work we detected the signal of segregation distortion on chromosome 6^25^ but did not consider the possibility that this signal was driven by multiple linked incompatibilities until several distinct signals were detected in the higher powered analysis included in this manuscript (Fig. 2). A series of analyses indicate that these results are unlikely to be an artifact of admixture linkage disequilibrium or variation in power along the chromosomes (Supporting Information 6). This increased resolution to map incompatibilities on the same chromosome allowed us to detect a new incompatible interaction between the *X. malinche* mitochondria and *X. birchmanni* ancestry near 20.3 Mb on chromosome 6. This peak contains a single MitoCarta gene, *atp5mg*. *atp5mg* forms part of Complex V, which combines mitochondrial and nuclear proteins in physical proximity and is essential for catalyzing ATP synthesis (Fig. 4E). This protein has several nonsynonymous substitutions between species, including at a conserved residue (Table 1; Fig. S19), and is differentially expressed between *X. birchmanni* and *X. malinche* (Fig. S5).

In addition, we found evidence of reduced sensitivity to Complex V inhibition in hybrids (Fig. 4), consistent with decreased Complex V function. Complex V is the final protein complex in the mitochondrial electron transport chain and is responsible for the production of ATP. Because of its position in the electron transport chain, reduced performance of Complex V could be consistent with issues in the function of this specific protein complex, or domino effects generated by reduced function of earlier components of the electron transport chain (e.g. Complex I). Regardless of the precise cause, these results reinforce our prior work connecting ancestry mismatch to dysfunctional mitochondrial function^25^, and suggest that this relationship may extend across multiple chimeric protein complexes in hybrids between *X. birchmanni* and *X. malinche* (e.g. Fig. 4; Fig. S11-S12).

Beyond these clear physiological signals of altered mitochondrial function in hybrids, we also document impacts on organism-level phenotypes driven by mitonuclear incompatibilities (Fig. 4). Our previous work studying mitonuclear incompatibilities between *X. birchmanni* and *X. malinche* identified several phenotypes associated with genetic incompatibilities, including abnormal embryonic development, abnormal heart development, and reduced physiological function of mitochondrial Complex I. Here, we raised a large number of F_2_ individuals in the lab and found a strong effect of genotypes across chromosome 6 on size. While individuals with the chromosome 6 incompatibilities suffer dramatically higher mortality (Fig. 4 ^25^), we found that individuals that survive on average are much smaller than their compatible siblings. Future work will be necessary to disentangle which of the chromosome 6 incompatibilities drives these growth defects or if there are synergistic effects, but together our results underscore strong phenotypic effects of several of the mapped mitonuclear interactions.

Our previous work identified strong DMIs that resulted in nearly complete hybrid lethality^25^. Our findings in the present study underscore an important role of mitonuclear interactions in the evolution of incompatibilities and paints a more complex picture of the ways in which they act in practice, including the identification of several linked incompatibilities and detection of interactions that have a less severe impact on hybrid viability. A key question raised by our work is whether the mitochondrion is unique in its web of interactions impacting hybrid fitness, or whether nuclear-nuclear incompatibilities follow similar patterns. Since the majority of studies only have power to detect hybrid incompatibilities with the strongest effects on fitness^27^, we consider this an open question in the field. Our results highlight the urgent need for more sensitive approaches to map hybrid incompatibilities as well as studies that examine their efficacy as barriers to gene flow in natural populations, allowing the field to move to a more complete understanding of the architecture of reproductive isolation between species.

## Acknowledgements

We thank members of the Schumer Lab for comments on earlier versions of this manuscript. This work was supported by NIH grant R35GM133774 to MS, HFSP grant Y81 to MS, NIH grant R35GM142836 to JH, EDGE - NSF IOS-2421661 to JH and MS, NSF IOS-1755327 to GGR, NIH R01GM115523 to PA, an NSF Graduate Research Fellowship to NR (20232146755), a CONACyT Fellowship to GR, and a Knight-Hennessy Scholars Fellowship and NSF Graduate Research Fellowship (2019273798) to BMM. We thank the Mexican Government for permission to collect fish (Permit No. PPF/DGOPA-002/19). Stanford University and the Stanford Research Computing Center provided computational support for this project.

## Data Accessibility and Benefit-Sharing

### Data Accessibility

All raw data will be deposited on NCBI sequence read archive (SRA XXXX). All processed data and phenotypic data from hybrids will be deposited on Dryad (Dryad doi XXXX). Code is available at https://github.com/Schumerlab.

### Benefit-Sharing

Data collection for this manuscript was performed in accordance with the Nagoya protocol on access and benefit sharing. Benefits from this research include sharing of data and results in public databases and partnerships with the Centro de Investigaciones Científicas de las Huastecas “Aguazarca”, A.C. to support research and outreach near our collection sites in Hidalgo, Mexico.

## Author contributions

Designed research: NVR, BMM, GIJ, PA, YB, JCH, GGR, MS. Performed research: NVR, BMM, MJRB, GIJ, TG, ENKI, DLP, SB, JJB, AS, YB. Analyzed data: NVR, BMM, MJRB, GIJ, AS, YB, MS. Wrote the paper: MS, NVR, MJRB, BMM.

