## Supplement for "Admixture mapping reveals evidence for multiple mitonuclear incompatibilities in swordtail fish hybrids"

**Supporting Information 1. Inferring the architecture of selection from admixture mapping data**

We were interested in inferring the architecture of selection on putative mitonuclear incompatibilities detected by admixture mapping. Specifically, we wanted to determine whether particular genotypes were depleted in combination with the *X. malinche* mitochondria, *X. birchmanni* mitochondria, or both. In controlled crosses, evidence of departures from mendelian expectations can be used to detect evidence of selection removing particular genotype combinations (or alternative mechanisms such as meiotic drive). However, in the context of natural populations, variation in admixture proportions and the presence of population structure makes it challenging to directly interpret such patterns in the data. Thus, to evaluate the likely architecture of each interaction, we used a simulation-based approach.

For each focal region, we performed simulations using the observed genotypes of each individual as well as their genome-wide admixture proportion. For a single simulation, we used the individual’s genome-wide admixture proportion to simulate their mitochondrial haplotype with a weighted binomial. Qualitatively, this produced covariation in (simulated) mitochondrial and nuclear ancestry similar to that observed in the real data. For each simulation, we stored the genotype matrix, and repeated the procedure 10,000 times. We then compared the observed genotype combinations at the focal locus to the simulated distribution. If the observed value fell below the 1% confidence interval of the simulated distribution, we considered this evidence that the particular genotype combination was underrepresented in our dataset compared to null expectations. We repeated this procedure for each focal region.

For example, for a given simulation of the incompatibility on chromosome 6 at 20.25 Mb, we simulated mitochondrial haplotypes as described above and used the observed nuclear genotypes at this locus. We then asked how many individuals with a simulated *X. malinche* mitochondrial haplotype were homozygous *X. birchmanni* at this site on chromosome 6 and recorded this number. We repeated this procedure 10,000 times and found that based on null expectations, 99% of simulations recovered ≥30 individuals with an *X. malinche* mitochondrial haplotype that were homozygous for *X. birchmanni* ancestry at 20.25 Mb on chromosome 6. By contrast, in our empirical data, only 10 individuals were observed with this genotype combination. This indicates that this genotype combination is underrepresented in our real data, even accounting for ancestry variation in the admixture mapping population. However, the number of individuals with an *X. birchmanni* mitochondrial haplotype that were homozygous *X. malinche* at 20.25 Mb on chromosome 6 overlapped with the simulated distribution. Forty-five individuals with this genotype combination were observed in the real data, which falls in the lower 20% quantile of the simulated data (95% confidence intervals: 38-64). We interpret these results as indicating that selection on interactions between 20.25 Mb on chromosome 6 and the mitochondrial genome are primarily driven by interactions with the *X. malinche* mitochondrial haplotype.

**Supporting Information 2. BLAST search for sequences homologous to MitoCarta proteins**

When investigating whether MitoCarta proteins were present in our admixture mapping intervals, we initially used *bedtools intersect* to overlap the intervals with the *X. birchmanni* genome annotation. However, in one of our admixture mapping regions on chromosome 4, we do not identify any annotated MitoCarta proteins. Moreover, in the admixture mapping region on chromosome 16, we identify one MitoCarta protein (Uqcrc2) but this protein lacks nonsynonymous changes between *X. birchmanni* and *X. malinche* and is not differentially expressed between species based on available RNAseq datasets^1^. This raised the possibility that our annotations were incomplete, and that we might be overlooking interacting mitochondrial genes in these regions.

To address this concern, we switched strategies to perform a targeted search for MitoCarta proteins genome-wide in *X. birchmanni*. We used blastp^2^ and the human MitoCarta protein database^3^ to search for hits in the *X. birchmanni* reference genome with an e-value threshold of 1^-20^. We evaluated these blast results genome-wide and subset regions that overlapped with our admixture mapping peaks. We found strong concordance overall between the genome annotation for *X. birchmanni* and blast search results. Moreover, we did not identify any previously undetected MitoCarta annotations in our admixture mapping regions using this approach.

**Supporting Information 3. Analysis of previously collected developmental data in F_2_ hybrids**

Past research from our group collected paired whole genome ancestry and developmental and respiration data from 235 F_2_ embryos with *X. malinche* mitochondrial haplotypes^4^. This initial work focused on relationships between developmental and respiration phenotypes in embryos and genotypes at *ndufa13* and *ndufs5*. Here, we reanalyzed the data to determine whether developmental and respiration phenotypes covaried with genotypes at newly identified mitonuclear incompatibilities. Since all hybrids in this dataset had the *X. malinche* mitochondria, we focused only on interactions that involved the *X. malinche* mitochondria or were bidirectional (Fig. 3; Table 1). We used the lme4 package in R to model embryo phenotype as a function of embryo genotype at the focal locus, median embryo length within the brood (a proxy for developmental stage, see^4^), and included brood identity as a random effect. Phenotypic data of interest for each embryo included individual length, head width, heart rate, sinu-atrium width, yolk diameter, and oxygen consumption. We implemented a Bonferroni correction to adjust p-values for multiple tests.

In past work we found distinct effects of ancestry mismatch between the *X. malinche* mitochondria and *X. birchmanni* ancestry at *ndufa13* and *ndufs5* (on chromosome 6 and 13 respectively) on rates of embryonic respiration and morphological defects^4^. In the main text we describe a newly detected phenotypic association with embryonic head width involving chromosome 4. Here, we discuss in more detail analyses pertaining to the mitonuclear interactions on chromosome 6. We previously found that *X. birchmanni* ancestry at *ndufa13* on chromosome 6 contributed to a number of embryonic (and adult) heart defects^4^. While the chromosome 6 intervals are not in linkage disequilibrium in natural hybrid populations, they are in strong linkage disequilibrium in lab crosses (Fig. S21), where F_2_ individuals have an average of 1.4 ancestry transitions per chromosome. This makes it difficult to distinguish their phenotypic effects, and in naïve tests treating each locus as unlinked, all three loci appear to have a strong impact on sinu-atrium width and heart rate (Table S4; Fig. S10). We evaluated whether there were sufficient recombination events in this region to evaluate only embryos that differed in genotype at the focal intervals along chromosome 6. However, we found that there were only 16 recombination events along this stretch of chromosome 6 in our sample of 235 F_2_ hybrid embryos. We thus concluded that we lacked sufficient data to distinguish between the effects of these loci in F_2_ hybrid embryos.

**Supporting Information 4. Approximate Bayesian computation inference of strength of selection**

*Simulations in SELAM for approximate Bayesian computation analyses*

We were interested in determining the likely strength of selection consistent with the empirical data at newly identified hybrid incompatibility loci. To do so, we used an approximate Bayesian computation (ABC) approach with simulations implemented in the forward time simulator SELAM^5^. SELAM allows users to model both population demographic history and the strength and architecture of selection via user-provided parameters. SELAM then models admixture and selection in a Wright-Fisher framework, and records ancestry for each individual along the genome. The ‘N’ parameter allows users to simulate mitochondrial inheritance as well as selection on mitonuclear interactions.

We focused on simulations modeling the chromosome 4 and chromosome 16 regions for several reasons. First, selection on these interactions have not been previously studied ⎯ past work focused on chromosomes 6 and 13^4^. Second, in the new admixture mapping dataset, mapped interactions on chromosome 6 and chromosome 15 are suggestive of multiple incompatibilities in physical proximity (see Results; Fig. 3B-D), complicating attempts to simulate selection on these regions.

For each mitonuclear interaction, we randomly drew demographic parameters based on previous estimates for the admixture mapping hybrid population^4,6^. We sampled from a uniform distribution for the number of generations since initial admixture (20-80) and hybrid population size (200-2,000). We sampled the initial admixture proportion from a normal distribution with an average of 0.35 *X. birchmanni* ancestry and a standard deviation of 0.02. We also simulated high rates of migration (*m*=3%) from parental populations based on empirical data from the admixture mapping population (see below).

To model selection on mitonuclear incompatibilities on chromosome 4 and chromosome 16, we specified the architecture of selection for each genomic region in separate simulations. For each simulation, we drew selection coefficients and dominance coefficients from uniform prior distributions (0-1). We drew two selection coefficients and dominance coefficients for each simulation, one for interactions with the *X. birchmanni* mitochondrial haplotype and one for interactions with the *X. malinche* mitochondrial haplotype. Following selection, we recorded the two-locus genotype of each individual and calculated the frequency of each genotype combination in the simulated population.

To determine whether to accept or reject a simulation, we compared simulated values to the observed data. We resampled the observed genotype combinations on chromosome 4 and 16 with replacement 1,000 times and recalculated the two-locus genotype frequencies for each replicate. If the SELAM simulation results fell within the 99% confidence intervals of the resampled distribution, we accepted the simulation. We repeated the SELAM simulations until we had accepted 500 parameter sets for the chromosome 4 and chromosome 16 incompatibilities.

*Discussion of migration rates simulated in approximate Bayesian computation analyses*

In the main text, we describe the results of approximate Bayesian computation (ABC) analyses to infer the strength of selection on newly mapped mitonuclear hybrid incompatibilities identified on chromosome 4 and 16. For these simulations, we implement a high migration rate from both parental species, motivated by empirical observations from this population.

Our admixture mapping population is the Calnali Low hybrid population. Unlike the majority of hybrid populations we have studied^4,7,8^, the Calnali Low hybrid population segregates for both mitochondrial types (Fig. 1). Both empirical data (see below) and simulations suggest that this is maintained by high levels of migration from source populations. Specifically, in simulations of selection on mitonuclear interactions without migration, hybrid populations rapidly fix for the mitochondrial haplotype of one of the parental species, and for compatible genotype combinations at the interacting nuclear locus. Indeed, when we performed ABC simulations modeling the Calnali Low population without migration, we were unable to accept any simulations out of 1 million parameter sets.

Available empirical data suggests that maintenance of both mitochondrial haplotypes is likely driven by high rates of migration. Based on sequencing data collected from a large sample from the Calnali Low hybrid population in 2022, 3% of individuals appeared likely to be recent migrants based on admixture proportion (i.e. >90% of ancestry genome-wide derived from one of the parental species). Thus, we model a migration rate of 3% in our simulations for ABC. However, we note that modifying the migration rate to other biologically realistic levels could impact our inference of selection and dominance coefficients.

**Supporting Information 5. Discussion of unexpected signal on chromosome 11**

The majority of signals that we detect with our admixture mapping approach are enriched for conspecific genotype combinations (e.g. Fig. S1). A positive correlation in ancestry between loci involved in incompatibilities is a general expectation of most models of incompatibility selection^9^. One exception to this pattern is the admixture mapping peak we detect at chromosome 11. Here the partial correlation between nuclear genotype and mitochondrial haplotype is negative (p<4 x 10^-7^; R = -0.21).

We explored this unexpected result by visualizing genotype combinations at this region (Fig. S2) and used simulations to determine which genotype combinations were over or under-represented (see Supporting Information 1). We observed a significant underrepresentation of individuals with *X. malinche* mitochondria and homozygous *X. birchmanni* ancestry at the chromosome 11 admixture mapping peak (p<0.001 by simulation). However, we observed an *overrepresentation* of the alternate genotype combination: individuals with *X. birchmanni* mitochondria that are homozygous *X. malinche* at the chromosome 11 admixture mapping peak (p<0.001 by simulation). Thus, the data could be consistent with strong asymmetrical selection against heterospecific ancestry in individuals harboring the *X. malinche* mitochondria. Other possible explanations for this pattern could involve more complex dynamics of selection or a false positive (given the expectation from our simulations that ~5% of signals may be false positives).

Given uncertainty about the drivers of this negative correlation in ancestry at the chromosome 11 peak, we refrain from discussing these results in the main text. We note however, that we searched for annotated MitoCarta genes within the chromosome 11 interval (18.29-18.66 Mb). We identified one MitoCarta gene in this interval, *C1qbp*. As in the main text, we investigated whether there were nonsynonymous or gene expression changes in these genes between *X. birchmanni* and *X. malinche*. We found no nonsynonymous substitutions that differentiated species, and no evidence of expression differences between species in previously collected datasets for the brain and liver^1^.

**Supporting Information 6. Decay in admixture linkage disequilibrium between admixture mapping peaks on the same chromosome**

In the main text, we discuss admixture mapping results where we find evidence for multiple peaks on chromosomes 6 and 15. To evaluate whether these multiple peaks represented truly distinct signals, we performed a number of analyses. We were most concerned that high admixture linkage disequilibrium (LD) in hybrid populations could result in selection on mitonuclear interactions in one region falsely driving the inference of selection on a nearby region. To evaluate this, we used plink^10^ to quantify the decay in admixture LD (average R^2^) over physical distance between regions within chromosome 6 and within chromosome 15. To do so, we identified the peak marker in a focal interval and evaluated decay in admixture LD over distance relative to the location of the other admixture mapping peaks. We found that in general, admixture LD between these regions decayed to the genome-wide background levels (Fig. S4).

To further evaluate possible technical causes of these signals, we generated alignments between the *X. birchmanni* and *X. malinche* versions of chromosome 6 and 15 from our Pacbio HiFi assemblies and examined them for evidence of small structural rearrangements in this region (Fig. S3). We also used RepeatMasker annotation files for the *X. birchmanni* genome to investigate whether there was a higher density of repetitive elements in these regions of the genome, which might be susceptible to increased mismapping, but found no evidence of such patterns.

Finally, we qualitatively evaluated whether patterns of missing data or skews in local ancestry in the regions between the admixture mapping peaks might decrease power and lead us to call multiple peaks instead of one larger peak spanning the whole region. We plotted the frequency of missing data (i.e. number of individuals covered per ancestry informative site) versus the location of admixture mapping peaks, as well as the average ancestry along the chromosome in individuals included in the admixture mapping dataset. We observed no correlation between drops in data quality or skews in ancestry and the locations of peaks in our admixture mapping dataset.

**Supporting Information 7. Simulations of expected power to detect segregation distortion**

In our F_2_ dataset from lab-raised individuals, we detect significant segregation distortion overlapping with mitonuclear incompatibilities on chromosomes 6 and 13. Because all lab hybrids have a *X. malinche* derived mitochondria, we do not expect to be able to detect segregation distortion at three of the incompatibilities we mapped that only interact with the *X. birchmanni* mitochondria (chromosome 15 at 19.6 Mb and 22.2 Mb and the chromosome 16 interval). However, we would expect *a priori* to detect segregation distortion at the chromosome 4 incompatibility and the chromosome 15 incompatibility at 17.4 Mb, which are inferred to involve both mitochondrial haplotypes (Supporting Information 1; Table 1). Despite this expectation, we do not observe expected patterns of segregation distortion at these regions.

The lack of a signal in lab hybrids could indicate that these incompatibilities are environmentally dependent, or that we lack power to identify them through segregation distortion analyses. Although our sample size of lab hybrids is large (~1750 individuals), F_2_ hybrids have experienced many fewer generations of selection than natural hybrids, making it unclear how we expect power to compare across the mapping scenarios. Thus, we performed simple simulations in R to evaluate our expected power to detect segregation distortion. Since all individuals in our dataset are F_2_ hybrids, based on Mendelian expectations 25% of individuals should be homozygous *X. malinche*, 50% heterozygous for ancestry, and 25% homozygous *X. birchmanni*. We performed simulations at a fixed selection coefficient (e.g. s=0.1) and drew the dominance coefficient from a uniform distribution ranging from 0-1. We used the selection and dominance coefficients to modify the expected frequency of each of the three genotypes post-selection, and simulated sampling of 1450 individuals (matching the lower 0.1% of individuals sampled in our F_2_ dataset to be conservative). We next calculated and recorded the allele frequency post selection and repeated this procedure 1,000 times per selection coefficient. We recorded the proportion of simulations for which the simulated allele frequency fell outside of the confidence intervals used in our segregation distortion analysis (see main text). We treated this proportion as our power to detect selection in artificial hybrids at that selection coefficient.

Our results indicate that we expect to have good power to detect selection on mitonuclear incompatibilities when *s*≥0.3 in our F_2_ mapping population. However, this means that we may have marginal power to detect some incompatibilities detected by admixture mapping in our lab hybrids. For example, the posterior distribution of *s* for the chromosome 4 incompatibility from ABC approaches ranges from 0.15-1 (95% confidence intervals of 0.22-0.98). This leaves us unclear whether the lack of segregation distortion localizing with admixture mapping peaks on chromosome 4 (or other chromosomes) is attributable to insufficient power or other factors, such as environmentally mediated selection.

**Supporting Information 8. Power to detect incompatibilities with different selection coefficients with an admixture mapping approach**

Previous mapping results for mitonuclear incompatibilities in this system identified incompatibilities with selection coefficients approaching 1 and past simulations indicate that we have good power to detect these strong interactions in our mapping population^4^. To evaluate our power to use admixture mapping to map loci with less extreme selection coefficients, similar to those inferred from ABC inference for chromosome 4 and chromosome 16 (see main text), we performed additional simulations of selection in our admixture mapping population using SELAM. We used the same simulation conditions for admixture proportions, migration rate, population size, dominance coefficient, and generations since initial admixture as we did for ABC simulations, but systematically varied *s_1_* and *s_2_* across simulations. For each set of simulations with a given *s_1_* and *s_2_*, we performed 100 replicates. From the SELAM output, we determined each individual’s genotype at the nuclear locus and each individual’s mitochondrial ancestry. We calculated genome-wide admixture proportion using 4 chromosomes, each 1 Morgan in length, that experienced no selection. We randomly subset 731 individuals from the simulated population to match the number of individuals used in our admixture mapping analyses.

This process resulted in nuclear genotypes at the selected locus, mitochondrial haplotypes, and genome-wide admixture proportions for 731 individuals from each simulation. This allowed us to calculate the partial correlation in mitochondrial and nuclear ancestry after accounting for genome-wide ancestry, as we had in the real data. We repeated this for each of the 100 replicate simulations for a given *s_1_* and *s_2_* and recorded the proportion of simulations in which the observed p-value was less than our equal to the genome-wide 5% or 10% false positive threshold for the real data.

For symmetrical incompatibilities, we found that we had excellent power for selection coefficients of 0.75, modest power for selection coefficients of 0.5, and weak power for selection coefficients of 0.25 (Fig. S24). For asymmetrical incompatibilities, we had weak power even at strong selection coefficients. In other words, although we detect many mitonuclear interactions with our large admixture mapping dataset, we only expect to have power to identify mitonuclear interactions under stronger selection. Thus, the true number of mitonuclear interactions impacting fitness in *X. birchmanni* x *X. malinche* hybrid populations remains unclear.

**Supporting Information 9. Building expectations for cline results with simulations of hybrid populations**

In the main text, we analyze patterns of clinal variation in one river, the Río Pochula, at loci involved in mitonuclear incompatibilities and matched regions from the genome-wide background. While researchers have generally assumed that loci involved in hybrid incompatibilities will have sharper clines than regions in the genome-wide background, we were interested in exploring this in simulations taking the architecture of selection on hybrid incompatibilities into account. Moreover, we wanted to incorporate biologically realistic parameters for *X. birchmanni* x *X. malinche* hybrid populations, such as asymmetry in migration. Specifically, migration tends to occur from upriver populations to downriver populations, resulting in a flow of *X. malinche* alleles down the river^11^, with substantially less gene flow in the alternative direction.

To explore these dynamics, we initiated simulations with 12 linearly arranged demes of *X. birchmanni*. We also modeled a deme zero, made up of pure *X. malinche* individuals. We simulated unidirectional migration between neighboring downstream demes (i.e. deme zero, migrates to deme one, deme one migrates to deme two, and so on). We set the population size in each deme to 1,000 and allowed the simulation to run for 200 generations. In separate simulations, we modeled three distinct types of incompatibilities: 1) incompatibilities when the *X. malinche* mitochondria was combined with homozygous *X. birchmanni* ancestry at the nuclear locus, 2) incompatibilities when the *X. birchmanni* mitochondria was combined with homozygous *X. malinche* ancestry at the nuclear locus, and 3) bidirectional incompatibilities with selection in both directions of ancestry mismatch. In Fig. S16 we present results for a lethal recessive mitochondrial by nuclear incompatibility. However, results were similar for sublethal incompatibilities. In each generation, we adjusted the expected frequency of each two-locus genotype as expected by selection and migration (for migration rates of 1% and 3% per generation depending on simulation), sampled parents from these expected frequencies, and then modeled maternal and paternal segregation and transmission to create the next generation. We conducted thirty replicate simulations for each parameter combination, and visualized ancestry at the mitochondrial and nuclear loci as a function of deme and incompatibility type.

We found that scenarios with bidirectional selection resulted in the most reduced introgression of nuclear loci over space in our simulations (Fig. S16). Scenarios with selection against interactions between the *X. birchmanni* mitochondria and *X. malinche* ancestry in the nuclear genome also resulted in lower rates of nuclear introgression compared to scenarios with selection against interactions involving the *X. malinche* mitochondria (Fig. S16). These observations from simulations fit qualitatively with our results from cline analysis, where interactions involving only the *X. malinche* mitochondria were not found as outliers in the cline analysis, whereas several interactions that were bidirectional or involved the *X. birchmanni* mitochondria were (Table 1; Fig. 5).

**Supporting Information 10.** **Expected genetic architecture of hybrid incompatibilities based on classic models**

Initial models of the evolution of hybrid incompatibilities envisioned a simple scenario where lineages harbored the same ancestral genotype (e.g. *aa-bb* at locus 1 – locus 2). Following the split of these lineages, lineage 1 may have a new mutation arise and fix at the first locus, becoming *AA-bb*, whereas lineage 2 could have a mutation arise and fix at the second locus, becoming *aa-BB*. While *A* and *B* alleles function properly in their genomic background, when introduced to each other in hybrids, they may not function properly, resulting in selection against *AA-BB* genotypes. However, the alternative heterospecific genotype combination, *aa-bb*, existed in the ancestor and thus is not expected to experience selection. This scenario can result in asymmetric selection on the two possible heterospecific genotype combinations (*AA-BB* and *aa-bb*) in hybrid incompatibilities, consistent with the results that we see for several loci in Table 1 and Fig. 3. Since it is often not straightforward to determine which heterospecific genotype combination is the ancestral genotype, the presence of asymmetric selection is often viewed as supporting this model. Note that these predictions hold even if both mutations occur in only one lineage as opposed to one mutation in each lineage.

These classic models assume that recurrent substitutions have not occurred at either locus, as might be expected under a model of coevolution of two loci or in certain scenarios of genetic conflict. In scenarios where multiple substitutions have occurred, both heterospecific genotype combinations may be incompatible, leading to what we refer to as “bidirectional” or “symmetric” incompatibilities in the main text. We identify three loci that fit this model in the current study (Table 1 and Fig. 3). Indeed, our results support the conclusion that significant turnover has occurred at the amino acid level between the mitochondrial genomes of the two species, and in some cases this is echoed in the interacting partners in the nuclear genome (e.g. *ndufs5*; Table 1). Thus, our results highlight exciting complexity in the genetic architecture of hybrid incompatibilities, with implications for models of how hybrid incompatibilities arise.

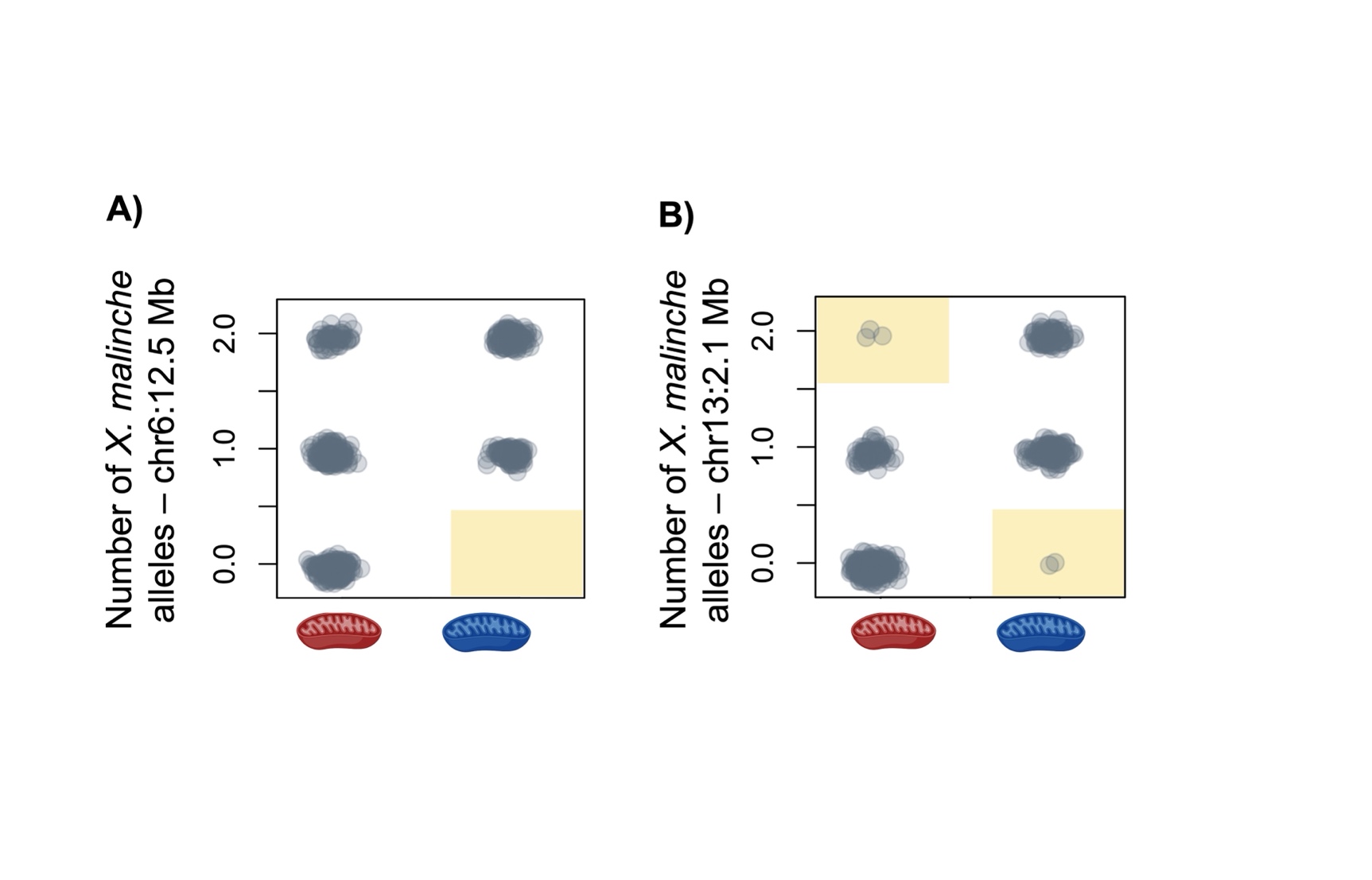

**Fig. S1.** Empirical results at previously mapped mitonuclear incompatibilities at 12.5 Mb on chromosome 6 near *ndufa13* and 2.1 Mb on chromosome 13 near *ndufs5* in our expanded admixture mapping dataset. Each gray point indicates one adult individual in our admixture mapping population as a function of their mitochondrial haplotype. Red indicates individuals with *X. birchmanni* mitochondrial haplotypes and blue indicates individuals with *X. malinche* mitochondrial haplotypes. Genotype combinations that are significantly depleted based on comparisons to null simulations (Supporting Information 1) are highlighted in yellow. These results mirror previously reported results for both regions in Moran et al.^4^.

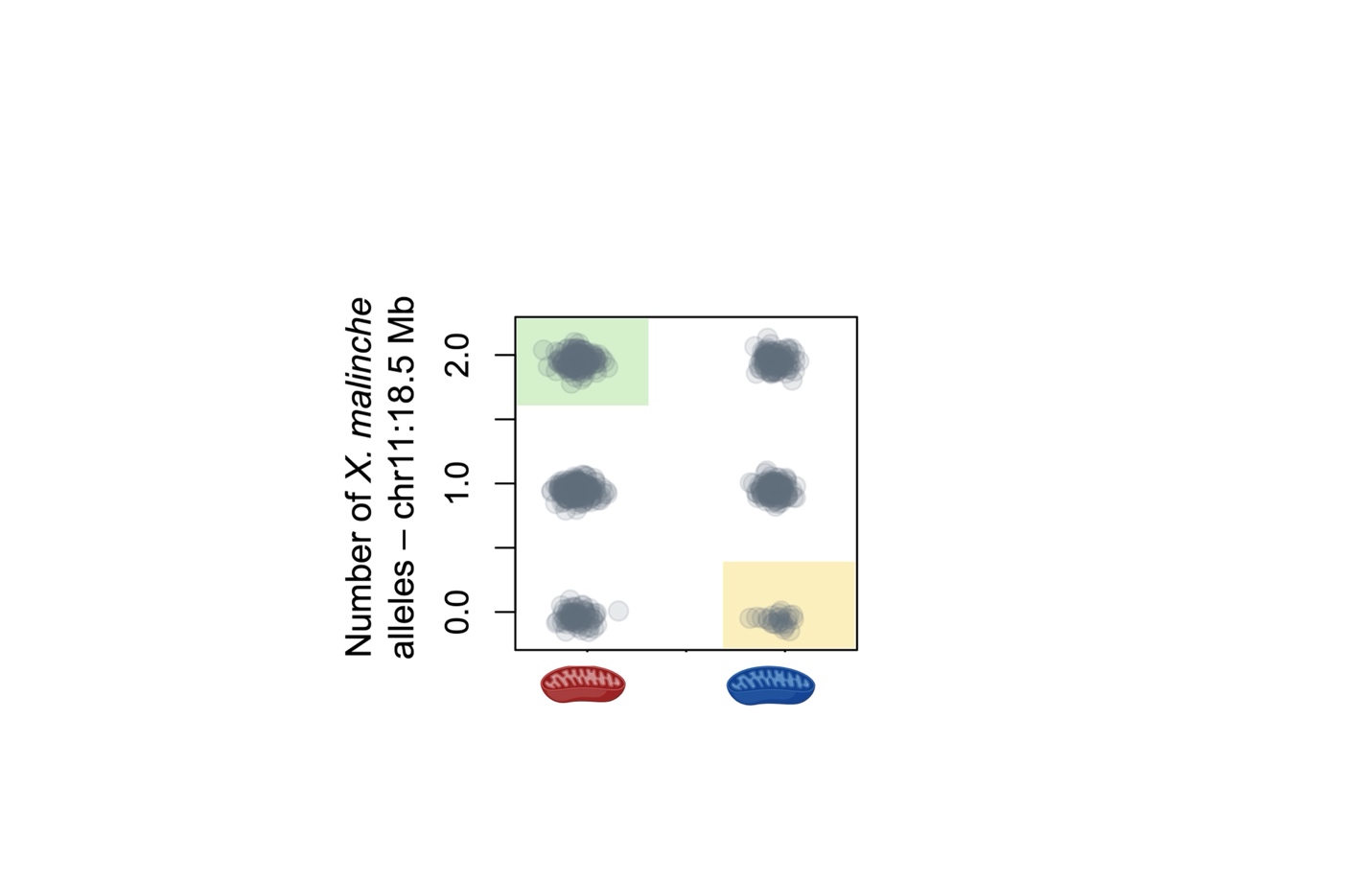

**Fig. S2.** Observed genotype combinations at the chromosome 11 peak in our admixture mapping dataset. Each gray point indicates one adult individual in our admixture mapping population as a function of their mitochondrial haplotype. Red mitochondria on the x-axis indicates individuals with the *X. birchmanni* mitochondrial haplotypes and blue mitochondria indicates individuals with *X. malinche* mitochondrial haplotypes. Genotype combinations that are significantly depleted based on comparison to null simulations are highlighted in yellow (see Supporting Information 1). Genotype combinations that are significantly enriched based on comparisons to null simulations are highlighted in green.

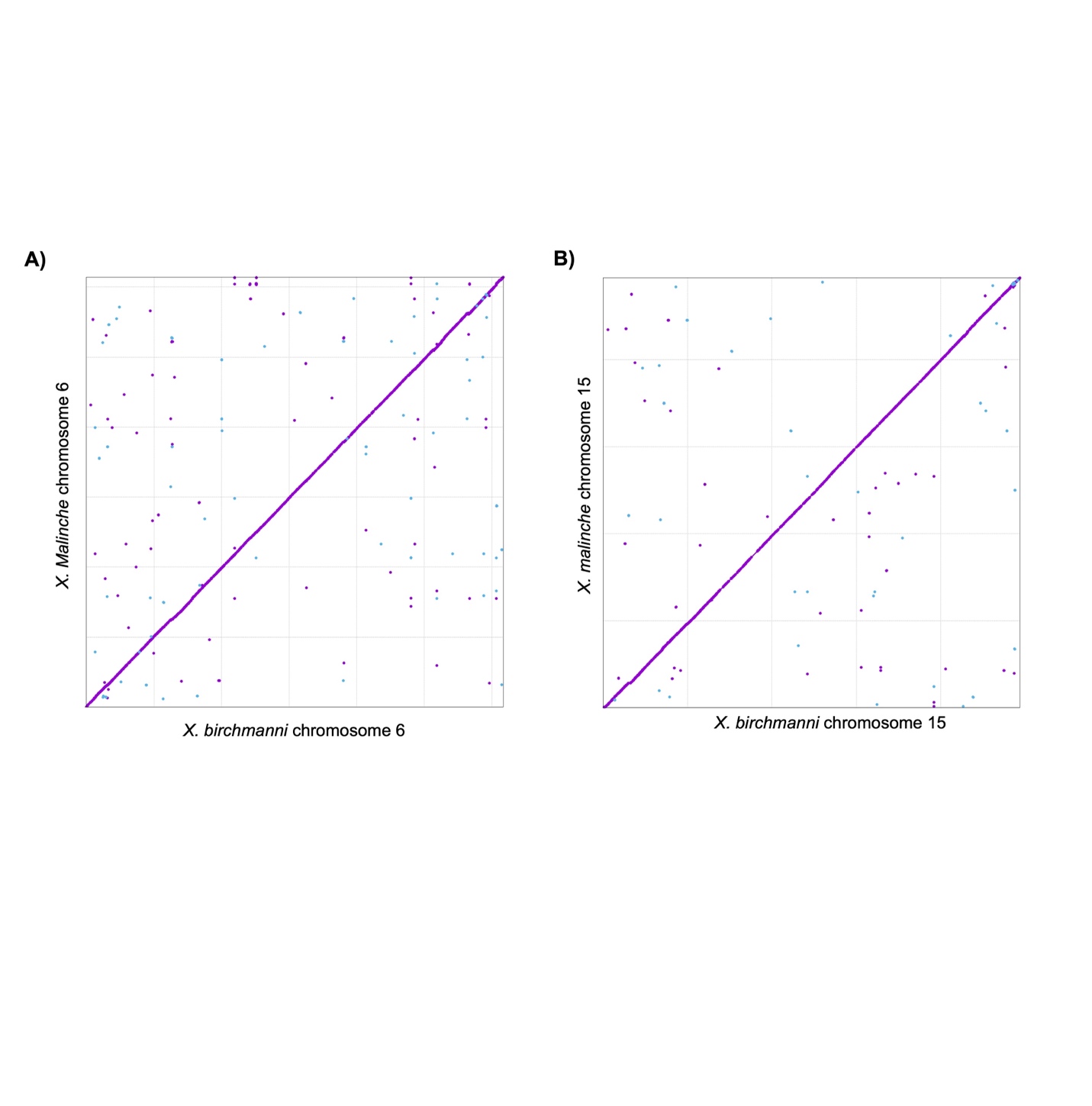

**Fig. S3.** Mummer alignments of **A)** chromosome 6 and **B)** chromosome 15 between PacBio HiFi assemblies of *X. birchmanni* and *X. malinche*. These alignments highlight complete synteny along both chromosomes, with no structural arrangements detected near admixture mapping peaks on either chromosome.

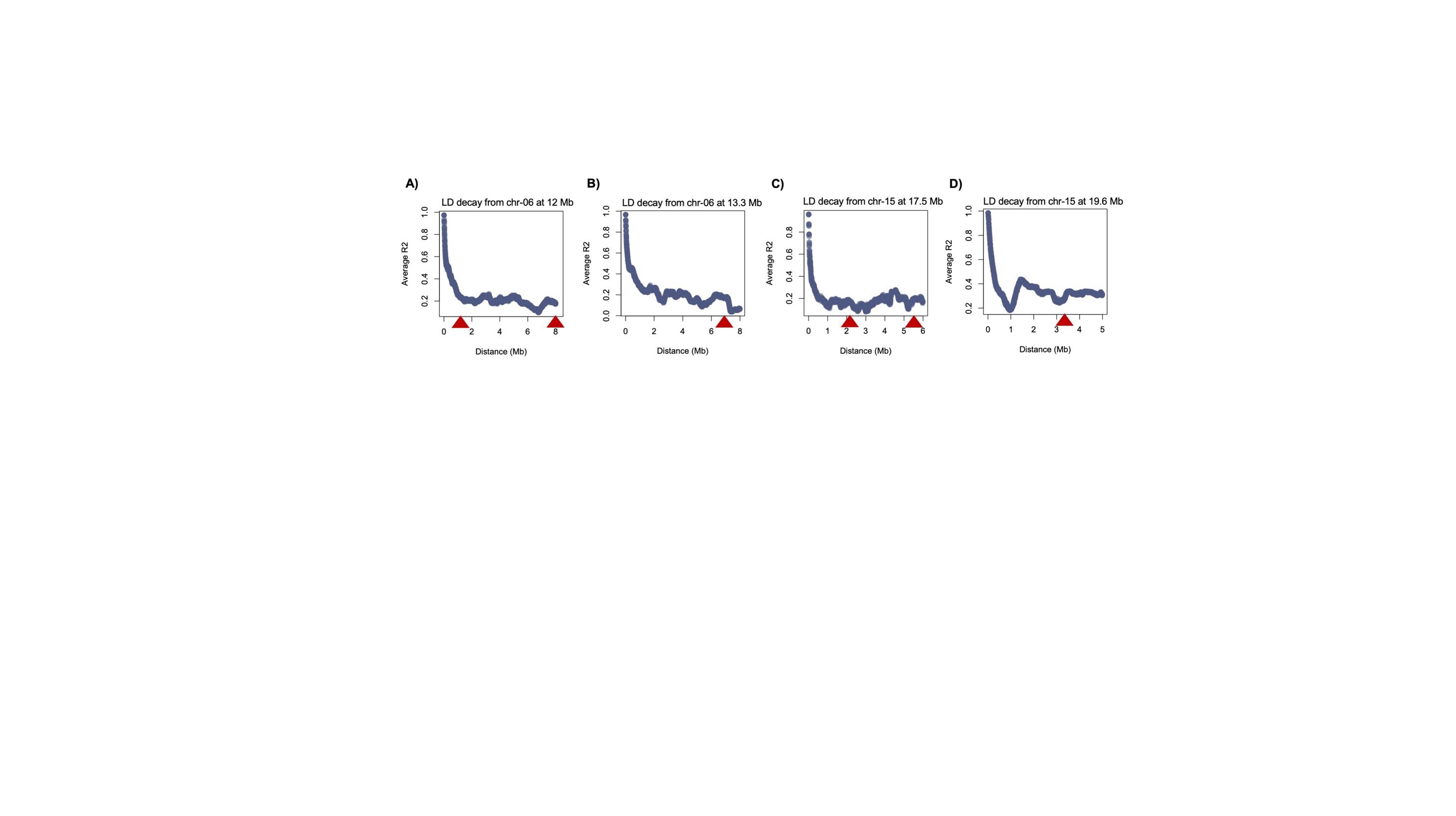

**Fig. S4.** Admixture mapping results presented in the main text suggest that there are multiple mitonuclear interactions involving nuclear loci on chromosome 6 and chromosome 15 (Fig. 2). To confirm that these signals are independent, we analyzed the decay in admixture linkage disequilibrium between each peak that occurred on the same chromosome. **A**) Decay in admixture LD, measured with R^2^, between the peak on chromosome 6 at 12 Mb and sites over the next 8 Mb. Red triangles indicate the approximate locations of signals from 13.1-13.6 Mb and 20.0-20.4 Mb. **B**) Decay in admixture LD between the peak on chromosome 6 at 13.4 Mb over the next 8 Mb. Red triangle indicates the approximate location of the peak at 20.0-20.4 Mb. **C**) Decay in admixture LD between the peak on chromosome 15 at 17.5 Mb and sites over the next 6 Mb. Red triangles indicate the approximate locations of the admixture mapping intervals from 19.4-19.9 Mb and 22.1-23.9 Mb. **D**) Decay in admixture LD between the peak on chromosome 15 at 19.6 Mb and sites over the next 5 Mb. Red triangles indicate the approximate location of the admixture mapping interval at 22.1-23.9 Mb. In each plot points show average R^2^ summarized in sliding 10 kb windows with a step size of 2 kb. Shorter total distances are plotted in **C** & **D** because the analysis reaches the end of the chromosome.

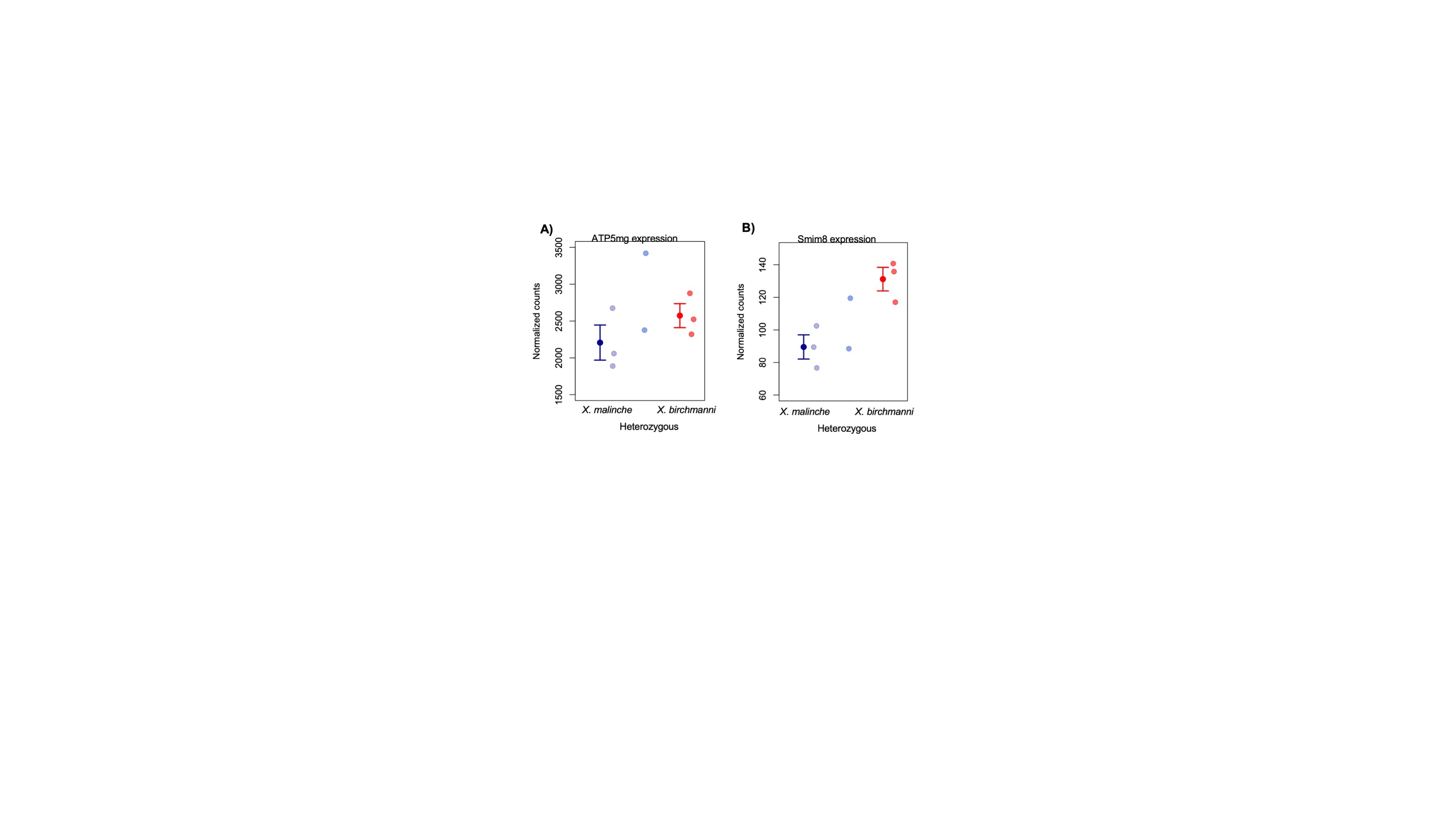

**Fig. S5.** Expression results for differentially expressed MitoCarta genes identified in admixture mapping peaks. **A)** *atp5mg* also shows evidence of differential expression in brain tissue between *X. malinche* (dark blue) and *X. birchmanni* (red). Expression in two F_1_ hybrids is also shown. Data is replotted from ambient temperature conditions in Payne et al. 2022^1^. Each semi-transparent point shows the data from one individual. The larger points show mean expression in the group and whiskers show two standard errors. Mean and standard errors are not plotted for F_1_ hybrids due to a small number of sampled individuals. **B)** Aside from *atp5mg*, one other MitoCarta gene that overlaps with our admixture mapping peaks, *smim8*, shows evidence of differential expression between the *X. malinche* and *X. birchmanni* parental species. Each semi-transparent point shows the data from one individual. The larger points show mean expression in the group and whiskers show two standard errors. Mean and standard errors are not plotted for F_1_ hybrids due to a small number of sampled individuals. Data is replotted from Payne et al.^1^ as above.

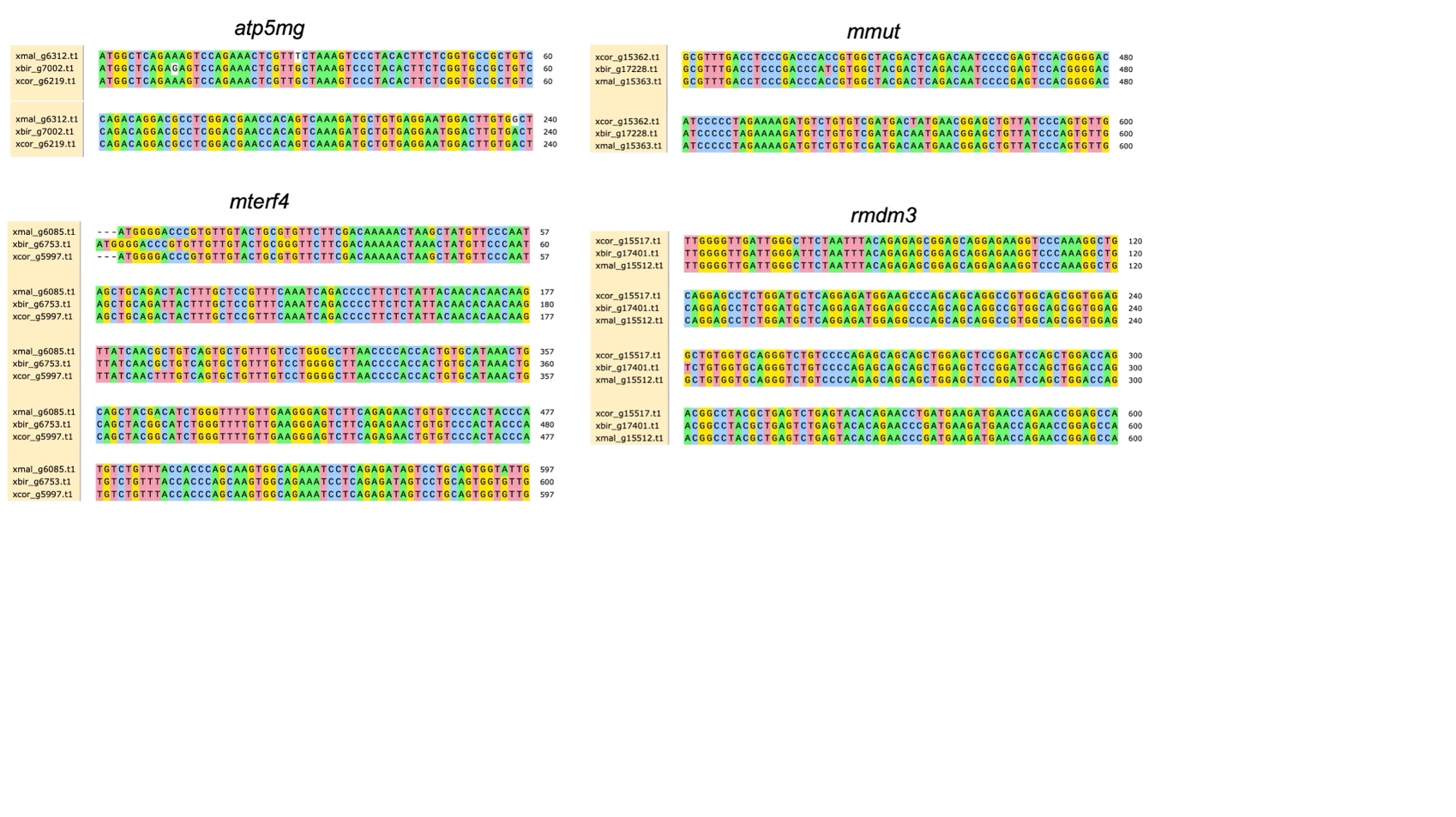

**Fig. S6.** Patterns of substitutions at newly mapped MitoCarta genes falling in admixture mapping peaks: *atp5mg*, *mterf4*, *mmut*, *rmdn3*. Shown here are alignments between annotated genes in *X. malinche* (xmal), *X. birchmanni* (xbir), and an outgroup, *X. cortezi* (xcor). Only partial alignments including variable sites are shown. Alignments were generated using clustal omega and visualized using SnapGene.

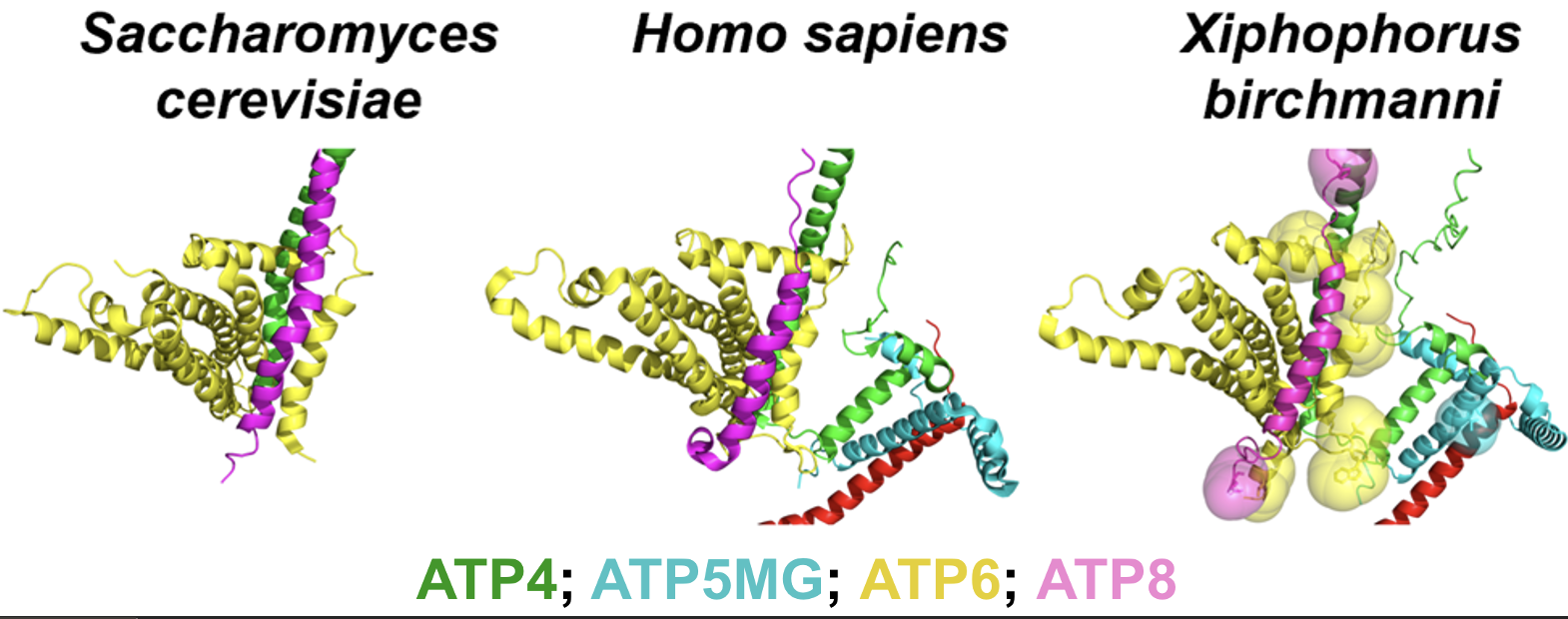
**Fig. S7.** Protein Data Bank Cryo-EM structures of ATP synthase (Complex V) from *Homo sapiens*^6^ and *Saccharomyces cerevisae* ^7^ and the predicted protein structure of this complex in *Xiphophorus*. In the predicted structure of *Xiphophorus*, the large spheres indicate amino-acid substitutions that differ between the *X. birchmanni* and *X. malinche* lineages.

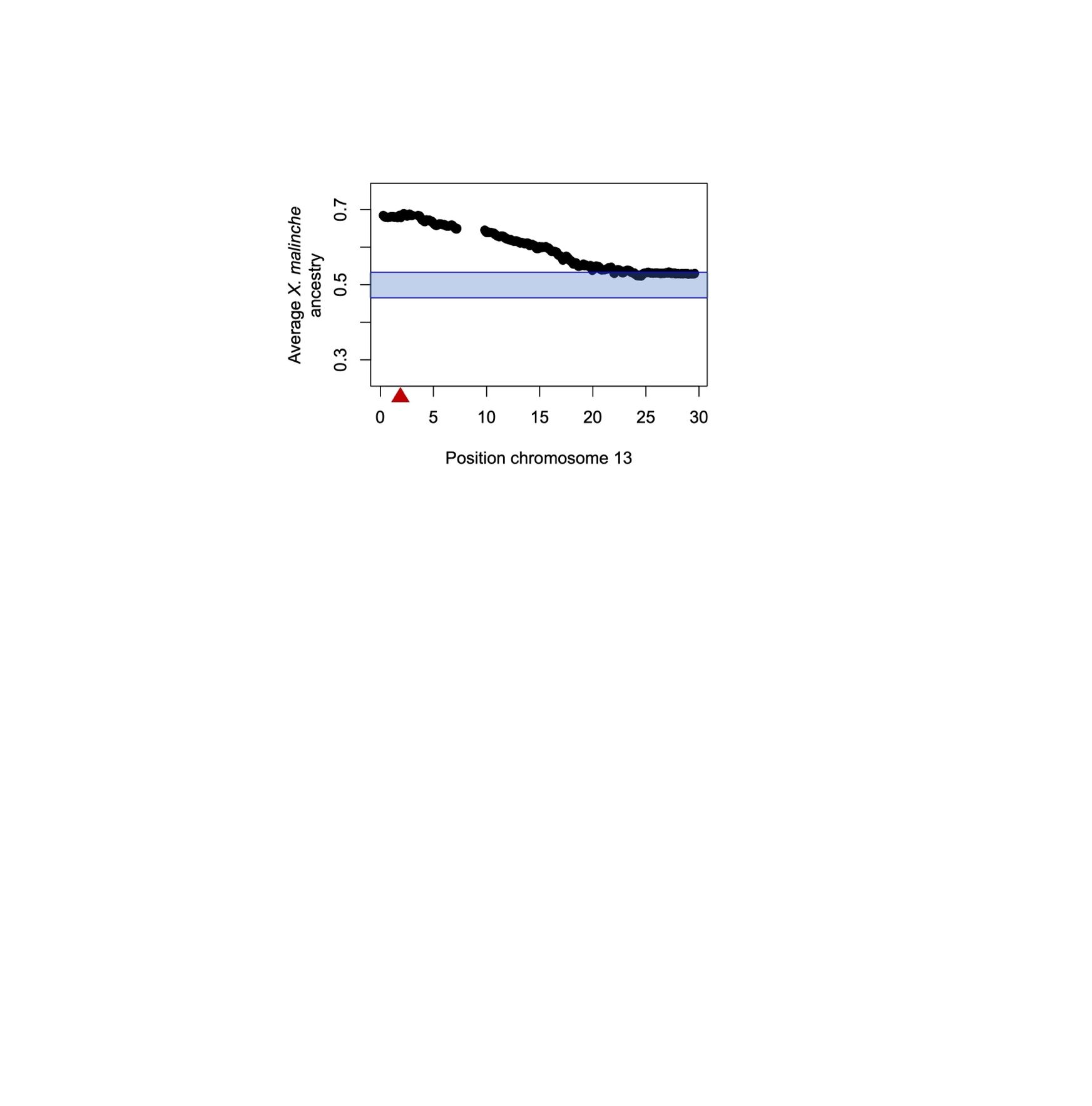

**Fig. S8.** Segregation distortion on chromosome 13 based on our larger F_2_ dataset. Red triangle indicates the location of *ndufs5* which causes lethal embryonic arrest when hybrids are homozygous *X. birchmanni* at this locus and harbor an *X. malinche* mitochondria. Blue lines and envelope indicate the 99% quantiles of expected ancestry in F_2_ hybrids from simulations matching our cross design.

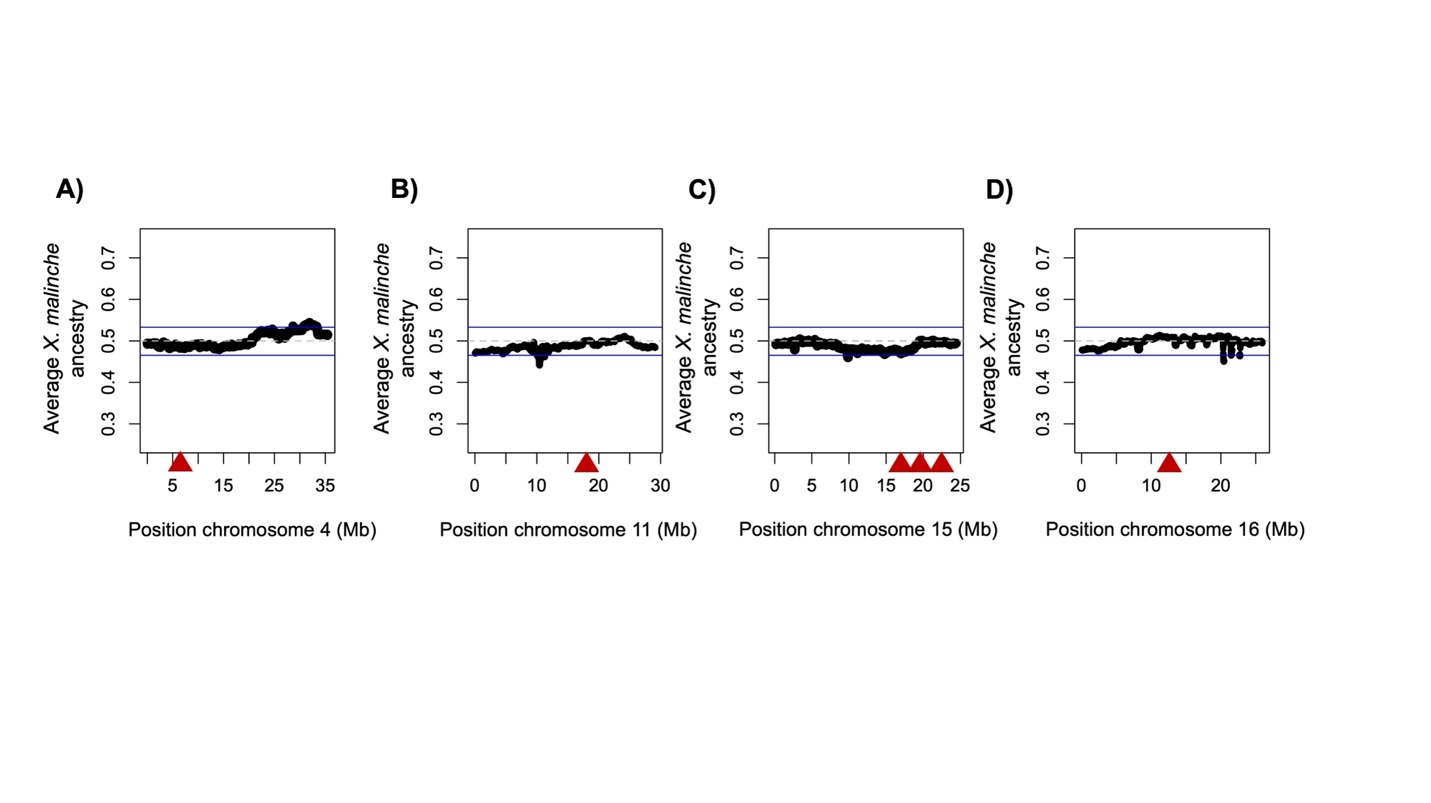

**Fig. S9.** Summary of average ancestry in F_2_ hybrids across additional chromosomes with evidence of mitonuclear incompatibilities from the admixture mapping dataset. Points show average ancestry at each ancestry informative site across F_2_ hybrids on chromosome 4 (**A**), chromosome 11 (**B**), chromosome 15 (**C**), or chromosome 16 (**D**). Dashed gray line shows expected average ancestry for this cross of 50%, blue lines show the genome-wide significance threshold for departures from this expected ancestry based on simulations (see main text). Red triangles show approximate locations of mitonuclear interactions identified through our admixture mapping approach.

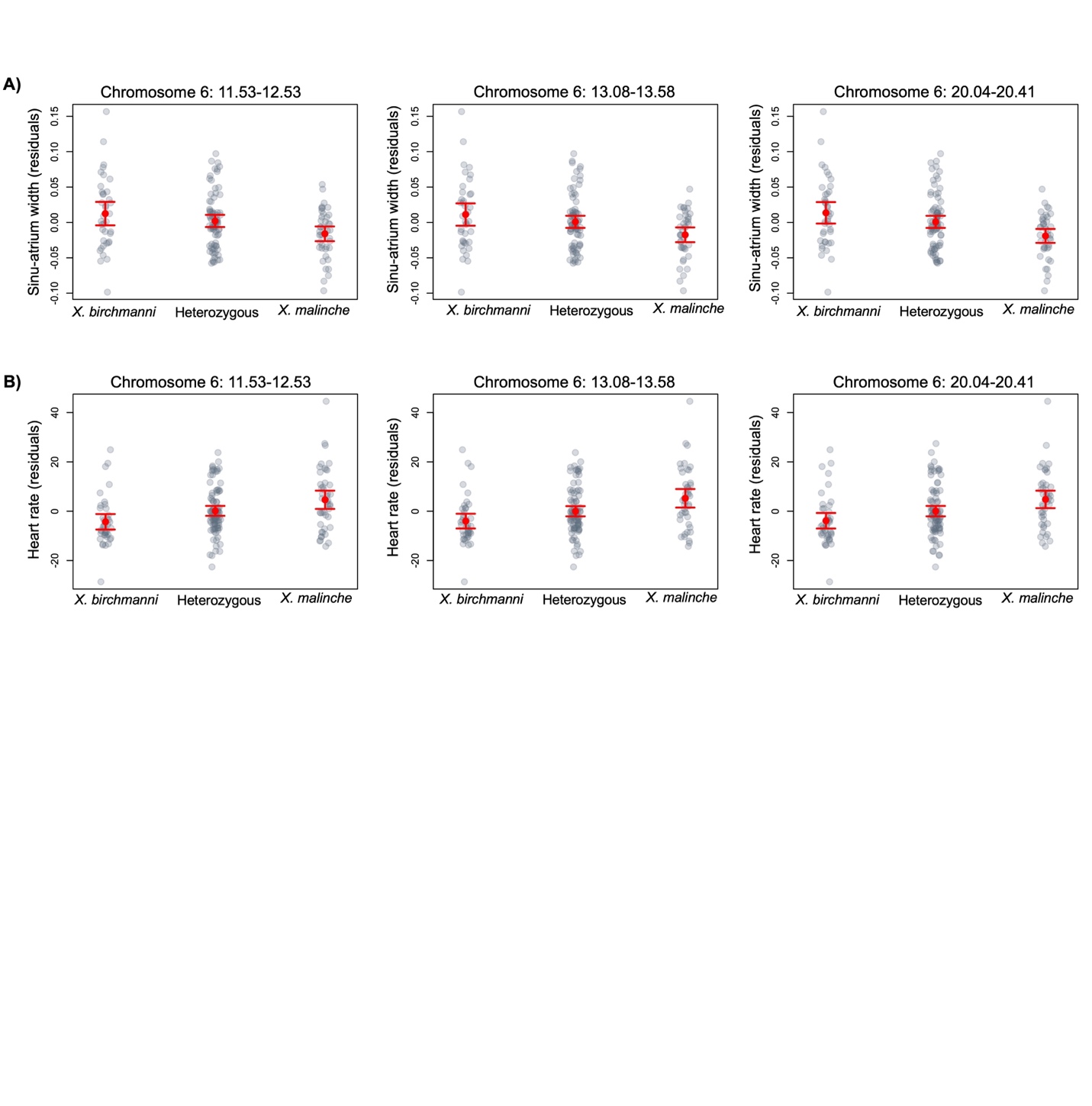

**Fig. S10.** Associations with embryonic heart phenotypes in F_2_ hybrids by locus on chromosome 6. **A)** Residuals of sinu-atrium width after accounting for embryo length and mother id (as a random variable) in a linear model at the three different admixture mapping peaks on chromosome 6. Red point shows the mean and whiskers show ± 2 standard errors. Gray points show individual data points. Note that statistical analysis was performed with a linear mixed model, residuals are for visualization only. **B)** Residuals of embryonic heart rate after accounting for embryo length and mother id (as a random variable) in a linear model at the three different admixture mapping peaks on chromosome 6. Red point shows the mean and whiskers show ± 2 standard errors. Gray points show individual data points. Note that statistical analysis was performed with a linear mixed model, residuals are for visualization only. Because of high admixture LD in artificial hybrids (Fig. S21), we cannot determine which of the chromosome 6 regions is driving the phenotypic patterns we observe in F_2_ hybrids with our current dataset.

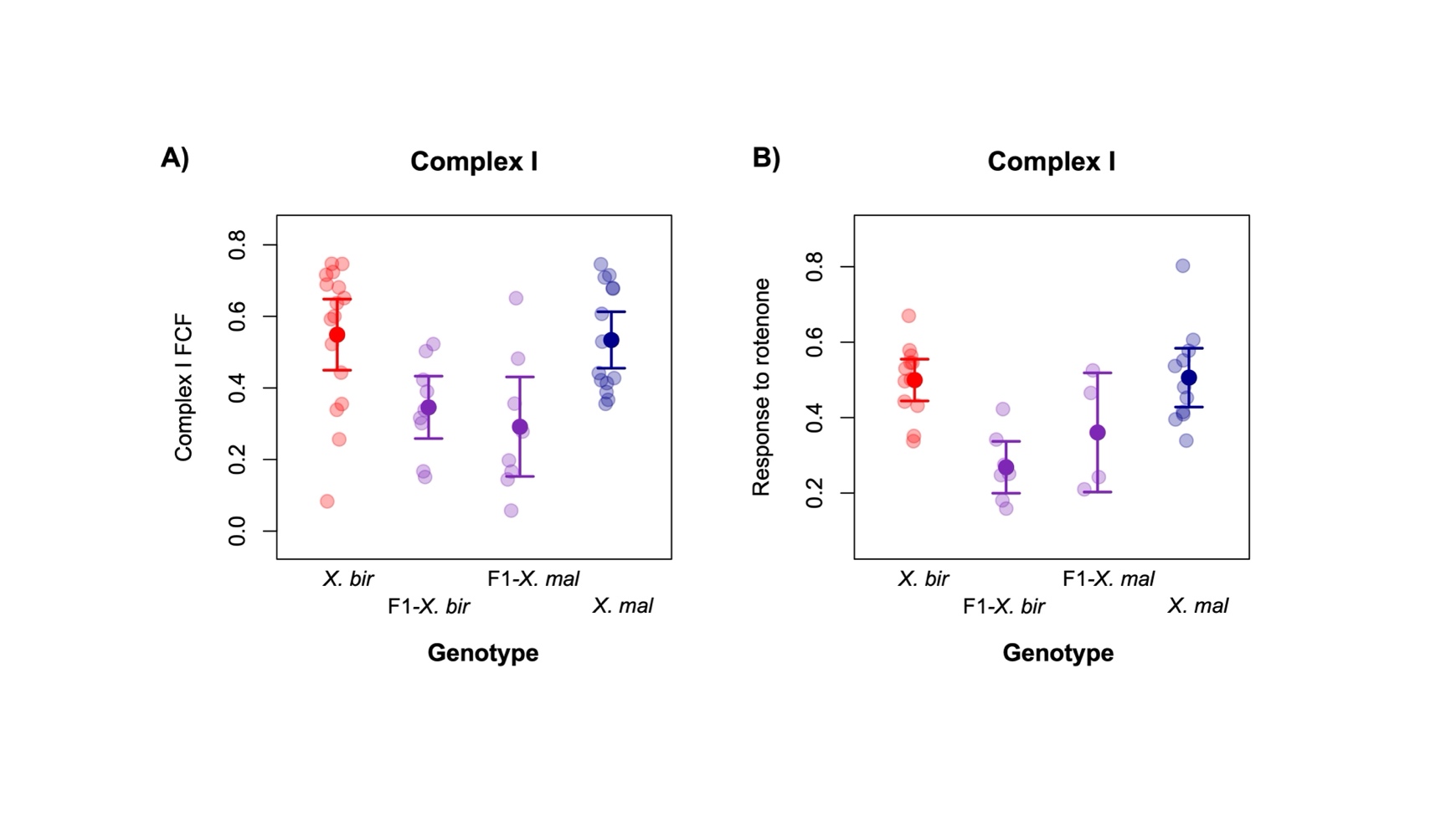

**Fig. S11.** Results of mitochondrial physiology assays using the Ouroboros O2K to interrogate Complex I function in pure *X. birchmanni* (*X. bir*), *X. malinche* (*X. mal*), and F_1_ hybrids with the *X. birchmanni* (F1 – *X. bir*) or *X. malinche* mitochondria (F1 – *X. mal)*. **A**) Complex I Flux Control Factor (Complex I FCF) measures the proportion of respiration attributable to oxidative phosphorylation in the absence of Complex II substrate. **B**) Rotenone FCF measures the proportional decrease in respiration with the inhibition of Complex I by rotenone, starting from a state with both Complex I and Complex II active, the inner mitochondrial membrane permeabilized, and Complex V inhibited. Semi-transparent points show individual data, points and whiskers show mean ± two standard errors.

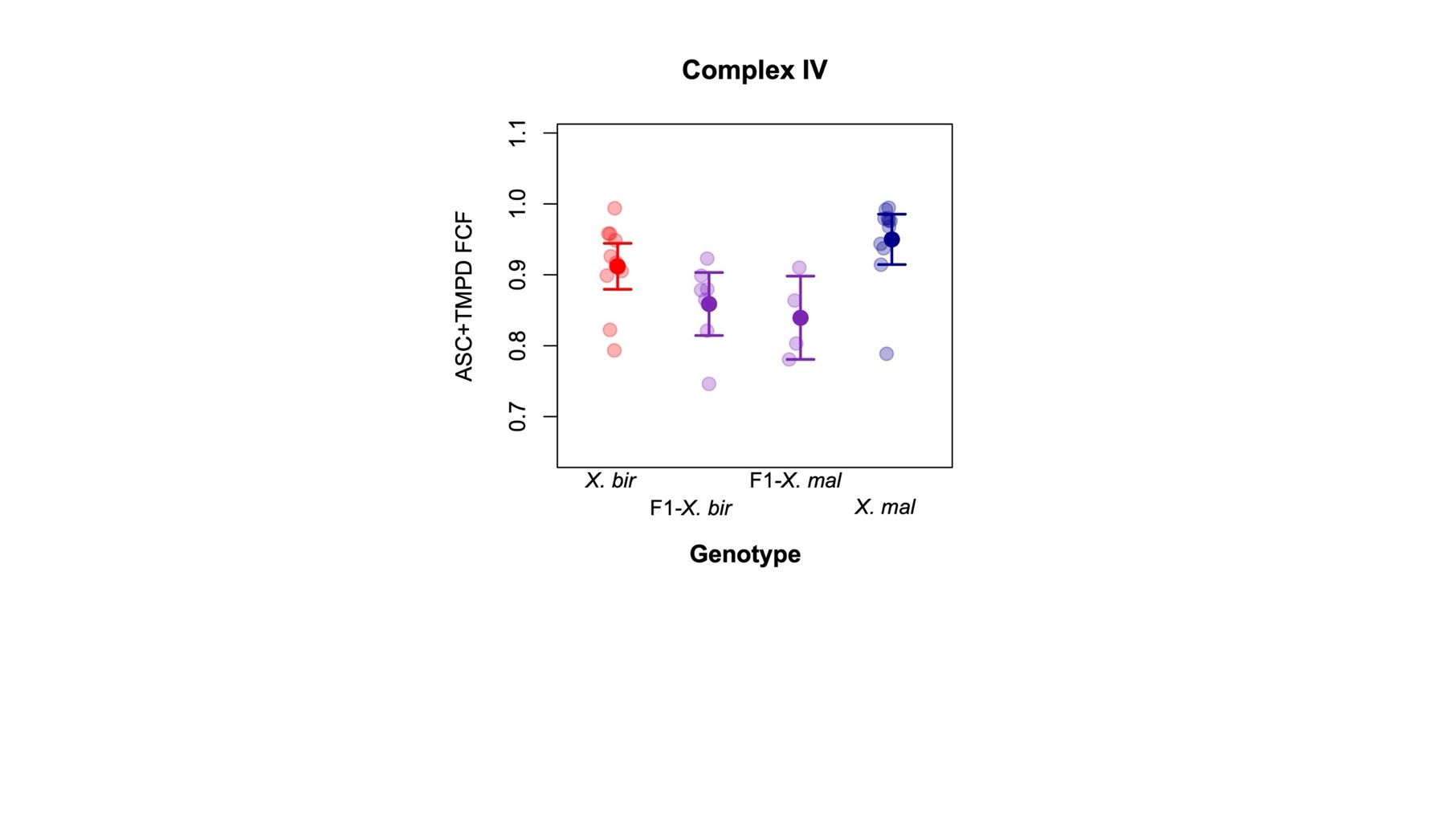

**Fig. S12.** Results of mitochondrial physiology assays using the Ouroboros O2K to interrogate Complex IV function in pure *X. birchmanni* (*X. bir*), *X. malinche* (*X. mal*), and F_1_ hybrids with the *X. birchmanni* (F1 – *X. bir*) or *X. malinche* mitochondria (F1 – *X. mal)*. Plot shows the maximum capacity of Complex IV to reduce oxygen when supplied with excess electron donors in the form of ascorbate and tetramethyl-p-phenylenediamine (ASC+TMPD FCF). Semi-transparent points show individual data, points and whiskers show mean ± two standard errors.

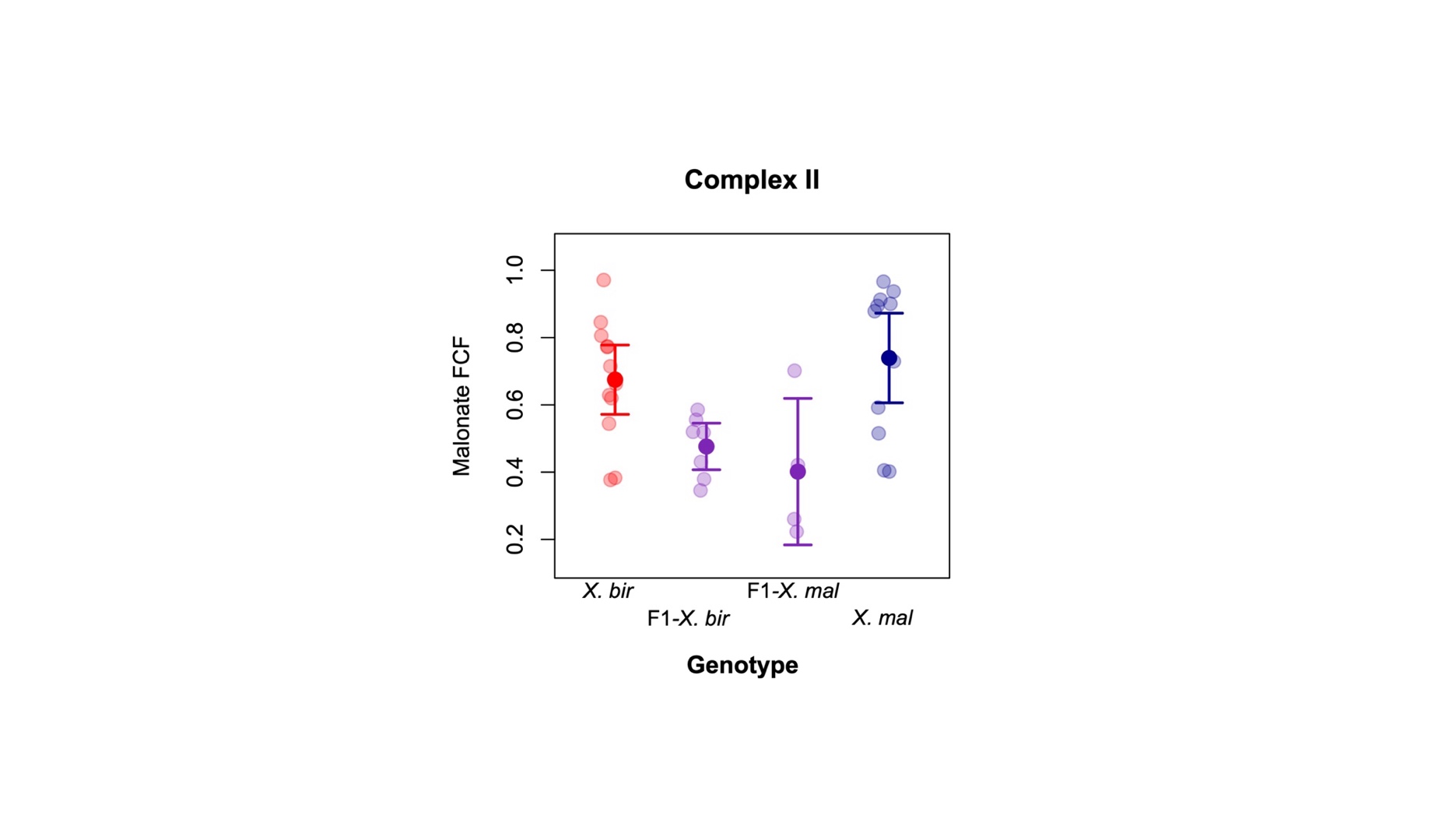

**Fig. S13.** Results of mitochondrial physiology assays using the Ouroboros O2K to interrogate Complex II function in pure *X. birchmanni* (*X. bir*), *X. malinche* (*X. mal*), and F_1_ hybrids with the *X. birchmanni* (F1 – *X. bir*) or *X. malinche* mitochondria (F1 – *X. mal)* based on response to malonate. The malonate flux control factor (Malonate FCF) measures the proportional decrease in respiration with the inhibition of Complex II, starting from a state with the inner mitochondrial membrane permeabilized, and both Complex I and Complex V inhibited. Semi-transparent points show individual data, point and whiskers show mean ± two standard errors. Note discordance between these results for Complex II function and those presented in Fig. 4.

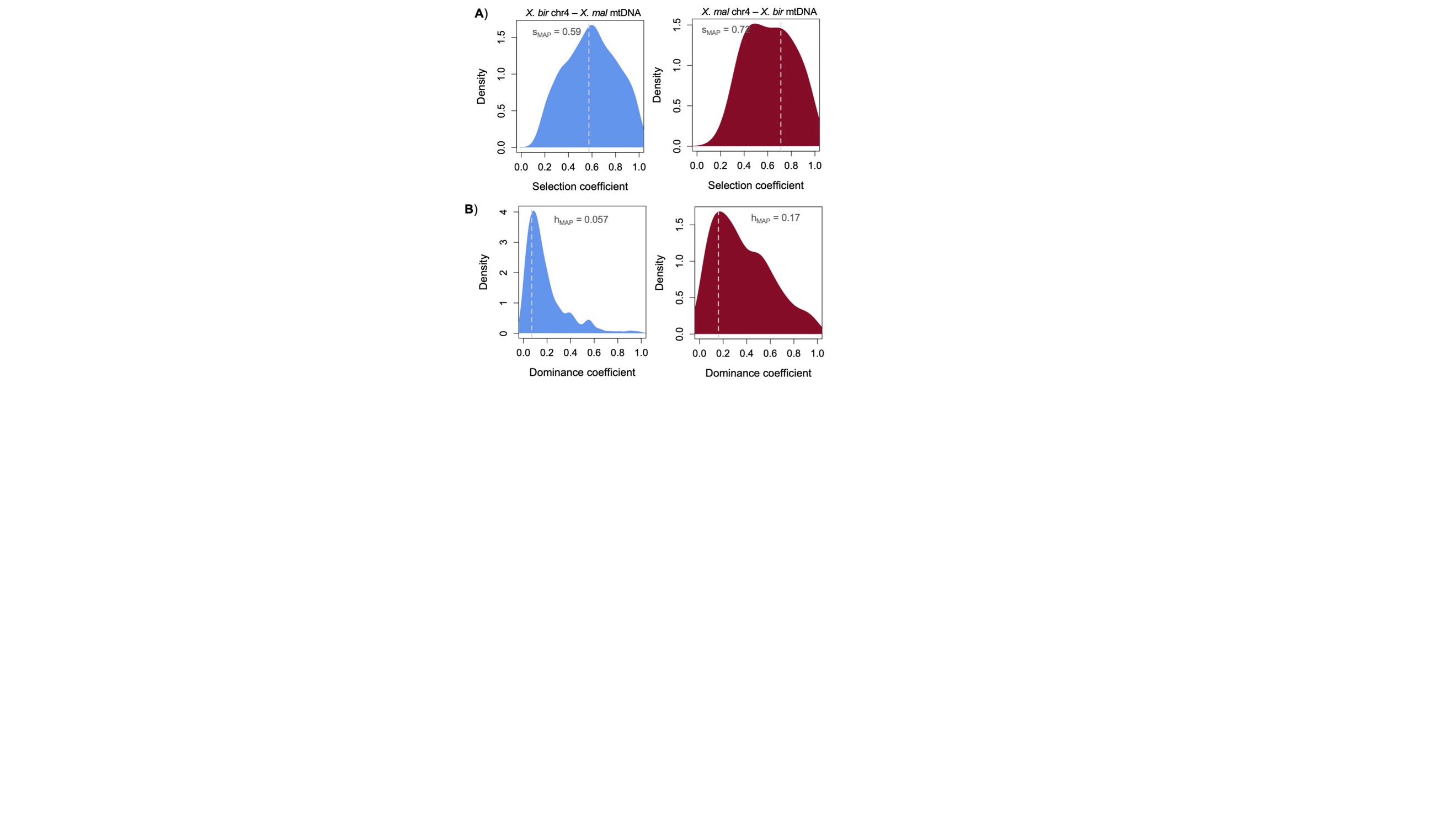

**Fig. S14.** Results of approximate Bayesian computation simulations estimating the strength of selection against mismatched ancestry between the mitochondria and the newly identified locus on chromosome 4. **A**) We infer substantial selection on both mismatch with the *X. birchmanni* mitochondria (blue distribution; *X. bir* chr4 – *X. mal* mtDNA) and the *X. malinche* mitochondria (red distribution; *X. mal* chr4 – *X. bir* mtDNA). Distribution shows all 500 accepted simulations, dashed line shows the maximum a posteriori estimate (MAP), which is also listed on the plot. **B**) We also recover well-resolved posterior distributions for the dominance coefficient in interactions with both the *X. birchmanni* mitochondria (blue distribution) and the *X. malinche* mitochondria (red distribution). Distribution shows all accepted simulations, dashed line shows the MAP estimate which is also listed on the plot. Based on these accepted simulations, we infer that both interactions involving this region on chromosome 4 are largely recessive.

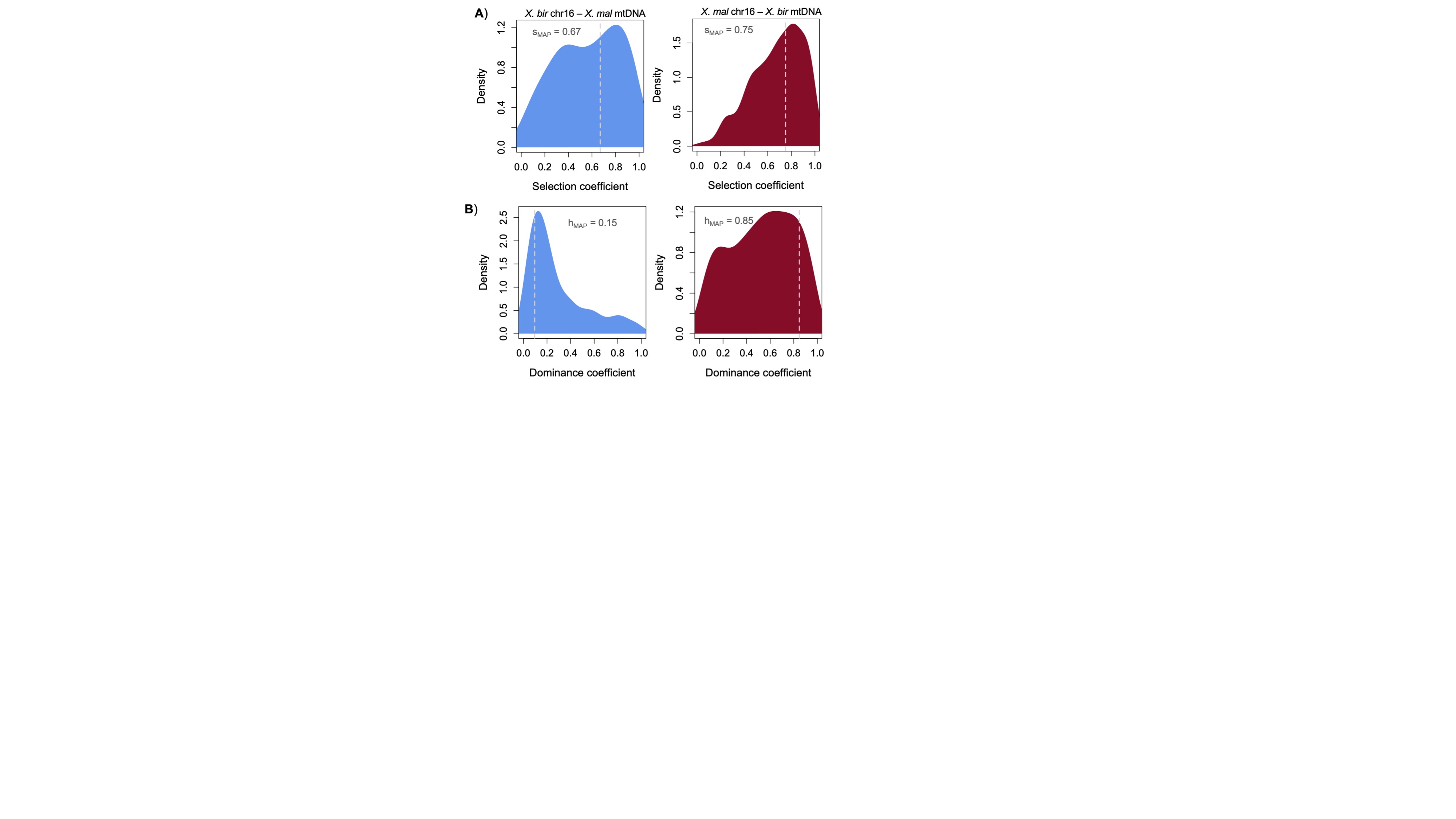

**Fig. S15.** Results of approximate Bayesian computation simulations estimating the strength of selection against mismatched ancestry between the mitochondria and the newly identified locus on chromosome 16. **A**) We infer substantial selection on both mismatch with the *X. birchmanni* mitochondria (blue distribution; *X. bir* chr16 – *X. mal* mtDNA) and the *X. malinche* mitochondria (red distribution; *X. mal* chr16 – *X. bir* mtDNA). Distribution shows all accepted simulations, dashed line shows the maximum a posteriori estimate (MAP), which is also listed on the plot. **B**) We also recover well-resolved posterior distributions for the dominance coefficient in interactions with both the *X. birchmanni* mitochondria (blue distribution) and the *X. malinche* mitochondria (red distribution). Distribution shows all accepted simulations, dashed line shows the MAP, which is also listed on the plot. Based on these accepted simulations, we infer that both interactions involving this region on chromosome 16 are largely recessive. We note that the architecture inferred here for chromosome 16 differs from simpler simulations discussed in Supporting Information 1.

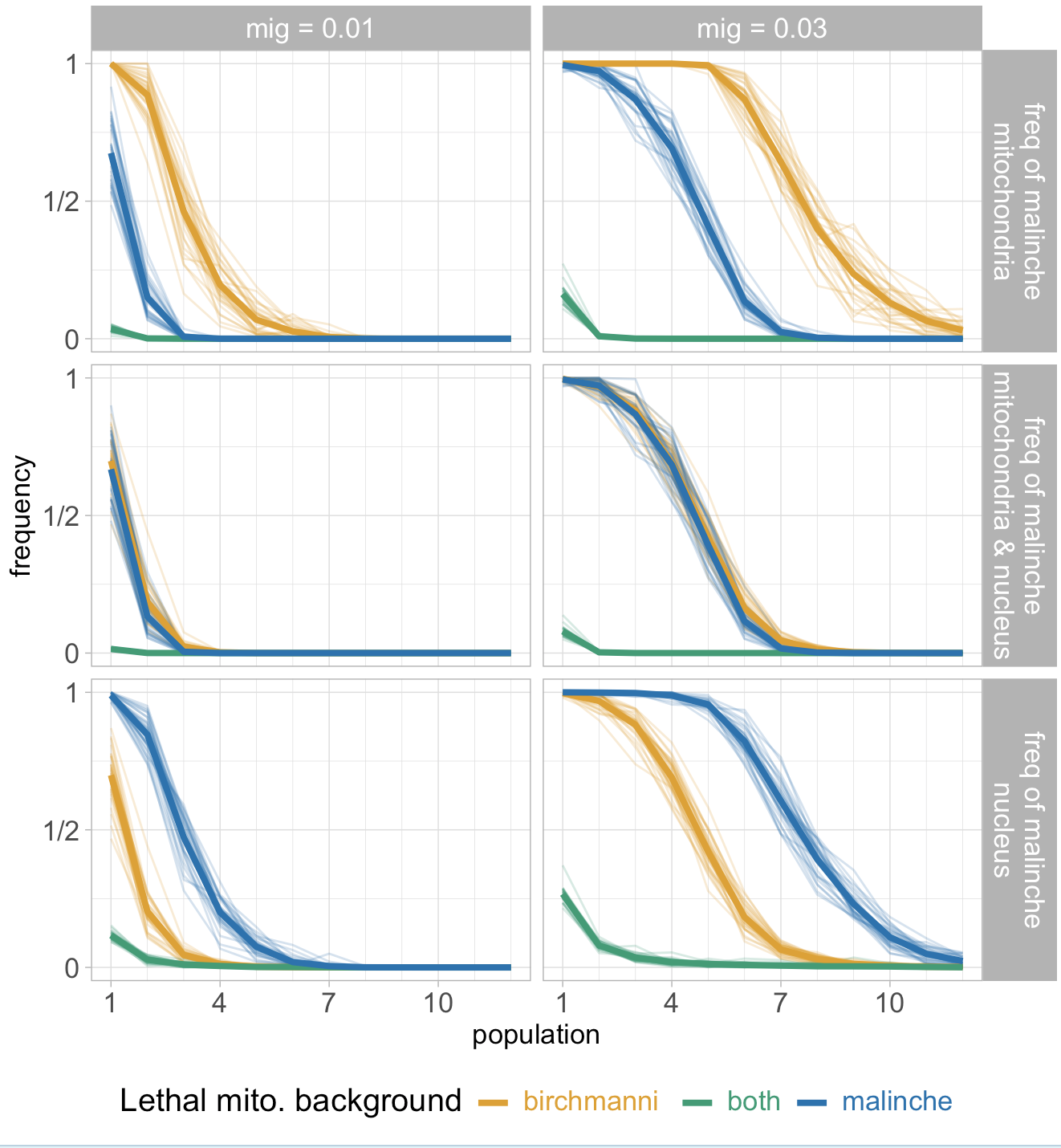

**Fig. S16.** Frequency of the *malinche* mitochondria (top row), the joint *malinche* nuclear/mitochondrial haplotype (middle row), and the *malinche* nuclear allele for three different incompatibility types. The three incompatibility types were 1) cases where homozygotes for the *malinche* nuclear allele were inviable in a *birchmanni* mitochondrial background (dark orange), 2) where either homozygote is lethal when combined with the alternate mitochondrial background (green), or 3) where homozygotes for the *birchmanni* nuclear allele are inviable when combined with the *malinche* mitochondrial background (blue). We assumed unidirectional migration rates of 1% (left column) or 3% percent (right column) from population zero (100% *malinche*) to population one, population one to population two, and so on, for two hundred generations. Heavy lines show means of thirty replicates; each replicate is indicated by light lines.

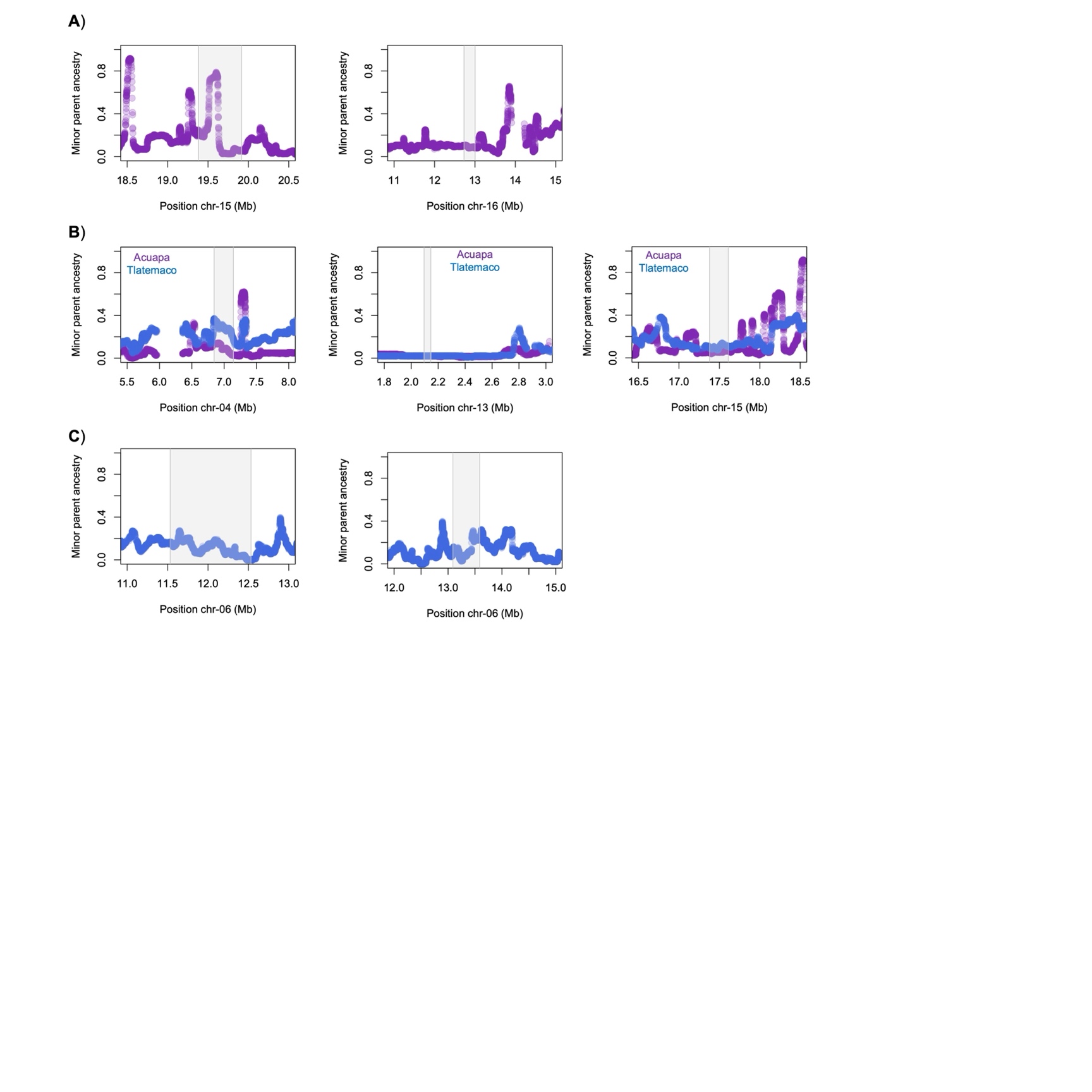

**Fig. S17.** Local ancestry at all mapped mitonuclear incompatibilities not shown in Fig. 5 in the Tlatemaco (blue) and Acuapa (purple) hybrid populations. Gray envelope indicates the associated region from admixture mapping. **A** depicts regions where we expect to see selection against minor parent ancestry in the Acuapa population given the inferred architecture of selection (i.e. the interaction involves the *X. birchmanni* mitochondria). **B** depicts regions where we expect to see selection against minor parent ancestry in both populations given the inferred architecture of selection. **C** depicts regions where we expect to see selection against minor parent ancestry in the Tlatemaco population given the inferred architecture of selection (i.e. the interaction involves the *X. malinche* mitochondria).

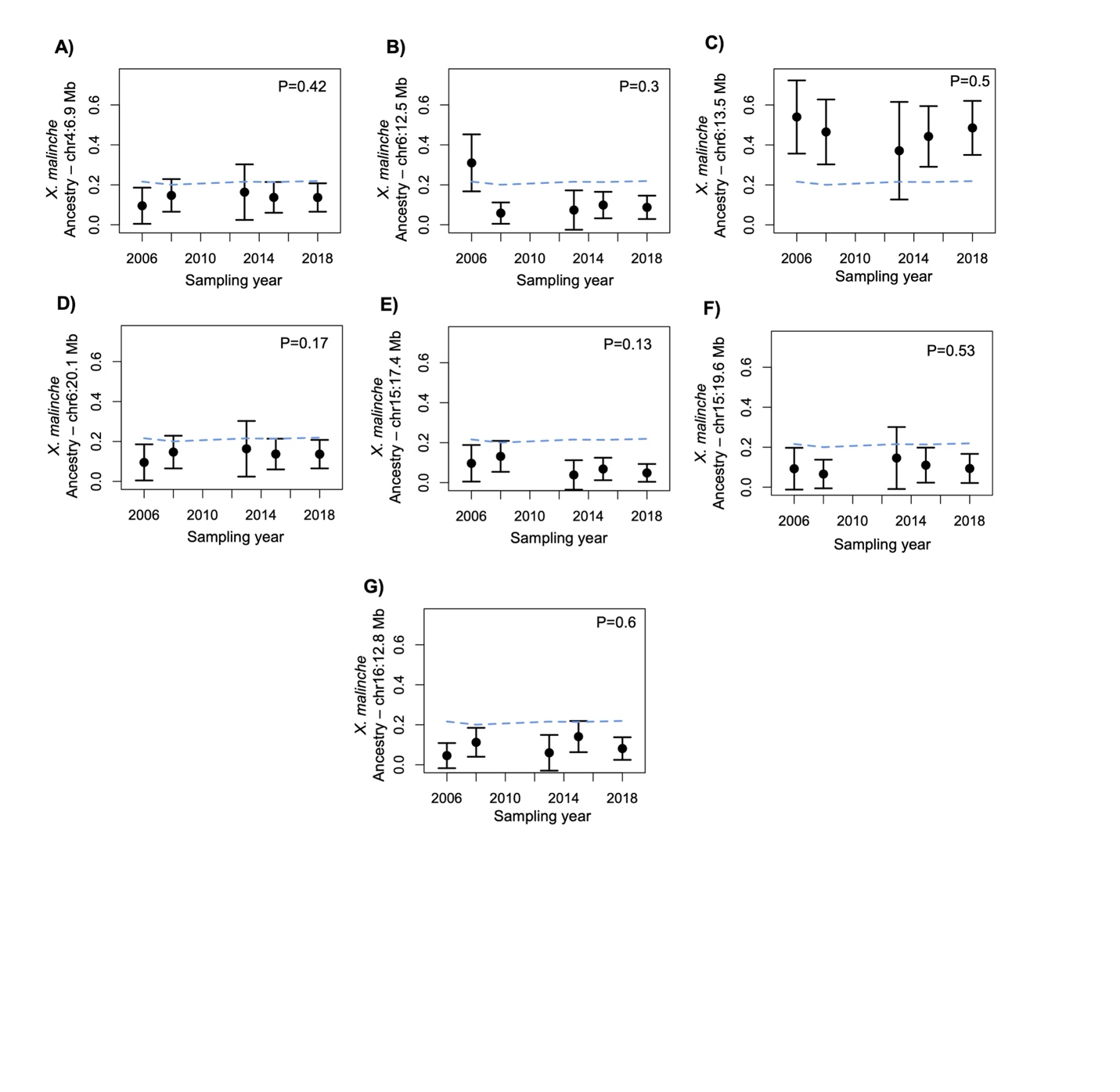

**Fig. S18.** Change in minor parent ancestry over time in the Acuapa population at additional admixture mapping loci (**A-G**). Points show mean ancestry at focal region in each year and whiskers show ± 2 standard errors. Dashed blue line shows average minor parent ancestry genome wide in Acuapa over the same time period. The inset text shows the p-value for the relationship between ancestry and sampling year based on a linear model. Note that depending on the genetic architecture of the mitochondrial interaction (see Table 1) and the strength of selection against the interaction, we may not expect to see significant changes in frequency. Specifically, we do not expect to detect interactions that primarily involve the *X. malinche* mitochondria (B, C, & D) since the *X. birchmanni* mitochondrial haplotype is dominant in the Acuapa population. Moreover, incompatibilities under strong selection are expected to be depleted in *X. malinche* nuclear ancestry by the beginning of our sampling, reducing our power to detect changes in ancestry during the focal time period.

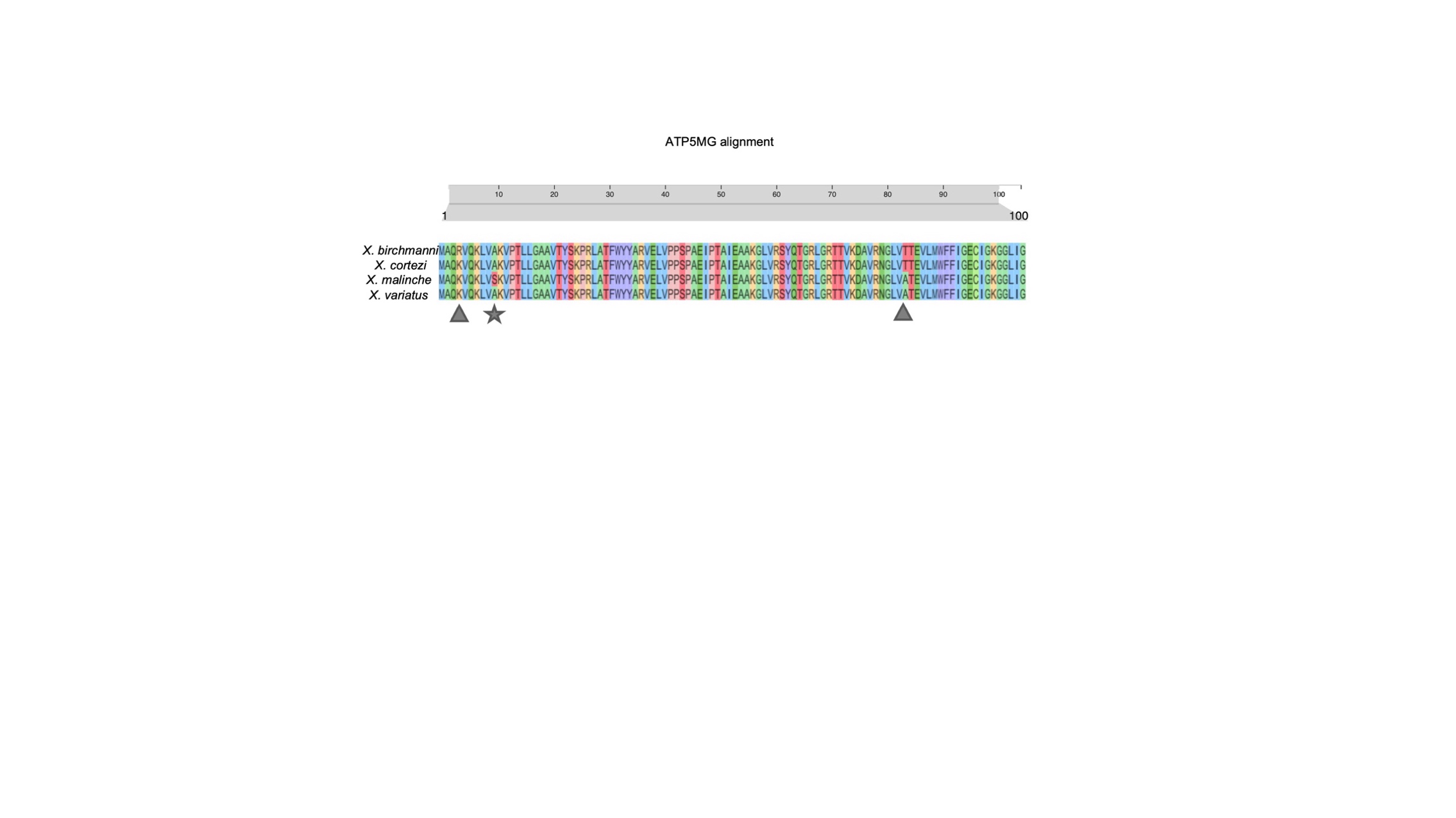

**Fig. S19.** Clustal omega alignment of predicted amino acid sequence of *ATP5MG* between *X. birchmanni* and *X. malinche,* and two outgroups, *X. cortezi* and *X. variatus*. Gray shapes highlight the locations of substitutions that differ between *X. birchmanni* and *X. malinche*. Gray star highlights the location of a substitution that is predicted not tolerated by SIFT.

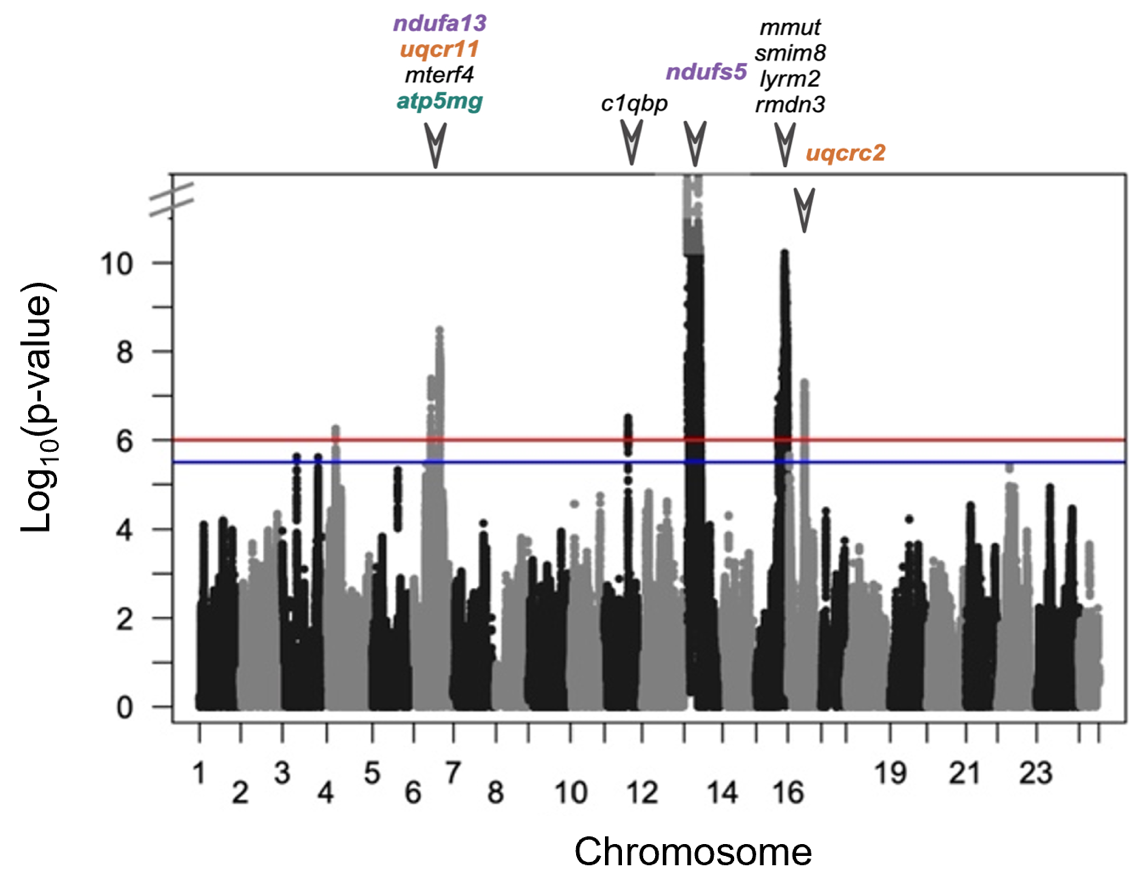

**Fig. S20.** Plot of admixture mapping results for association between mitochondrial and nuclear ancestry across the genome with the y-axis truncated for visualization of signals close to the genome-wide significance threshold (i.e. all peaks except the one identified on chromosome 13). Red line represents the 5% false positive rate threshold, blue line represents the 10% false positive rate threshold. Triangles and gene names indicate MitoCarta annotated genes associated with each admixture mapping interval. Separate intervals on the same chromosome are collapsed here for visualization purposes. Colored text highlights mitochondrially interacting genes that localize to particular protein complexes (see Fig. 2B for color coding).

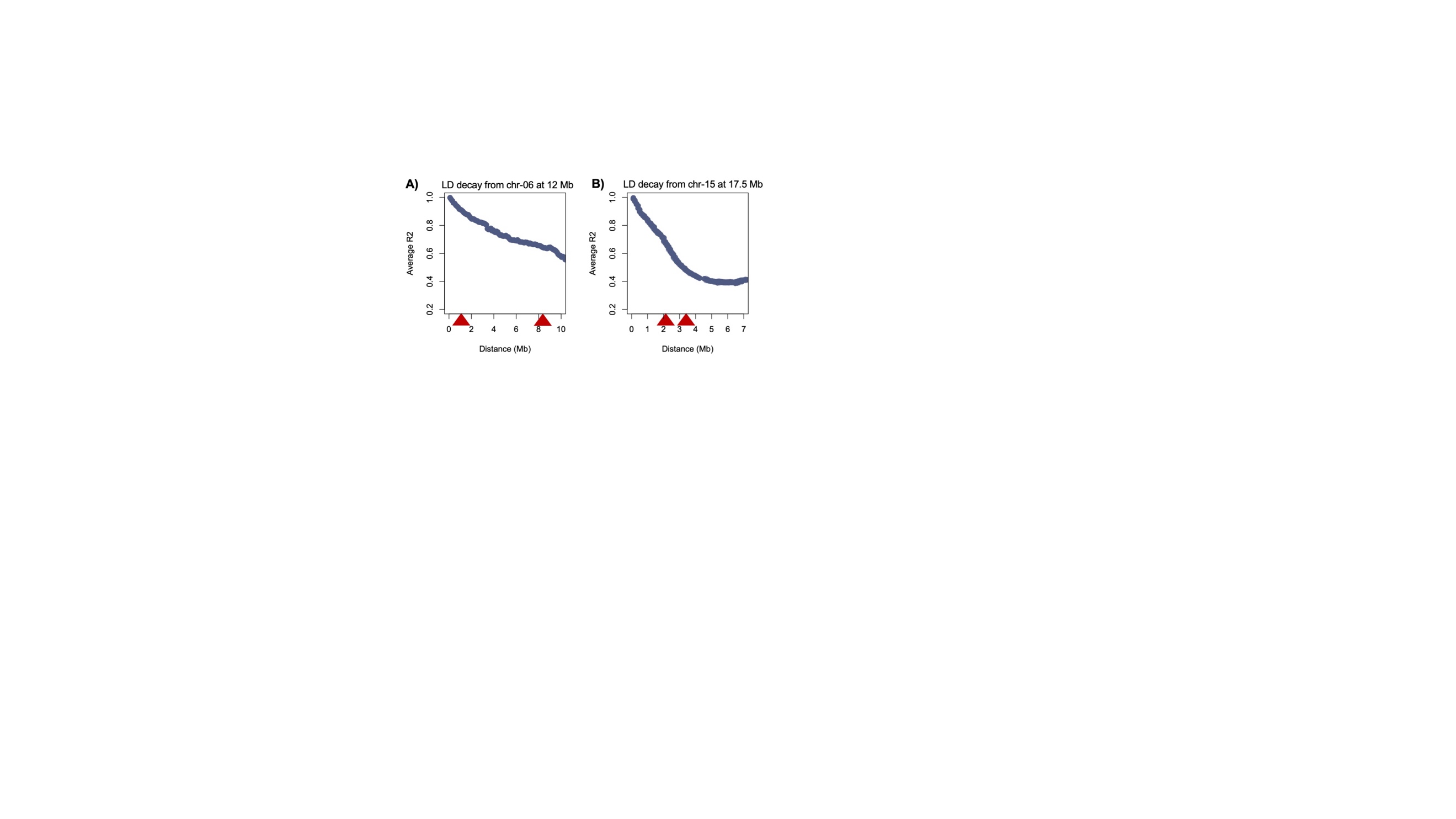

**Fig. S21.** Decay in admixture LD over physical distance in F_2_ artificial hybrids focusing on regions of interest on chromosome 6 and chromosome 15. **A)** Decay in admixture LD between the peak on chromosome 6 at 12 Mb and sites over the next 10 Mb. Red triangles indicate the approximate locations of signals from 13.1-13.6 Mb and 20.0-20.4 Mb. **B**) Decay in admixture LD between the peak on chromosome 15 at 17.5 Mb and sites over the next 7 Mb. Red triangles indicate the approximate locations of signals from 19.4-19.9 Mb and 22.1-23.9 Mb. Slow decay of admixture LD in artificial hybrids means that we cannot separate signals that may be driven by loci that are physically linked in early generation hybrids. In each plot, points show average R^2^ summarized in sliding 10 kb windows with a step size of 2 kb.

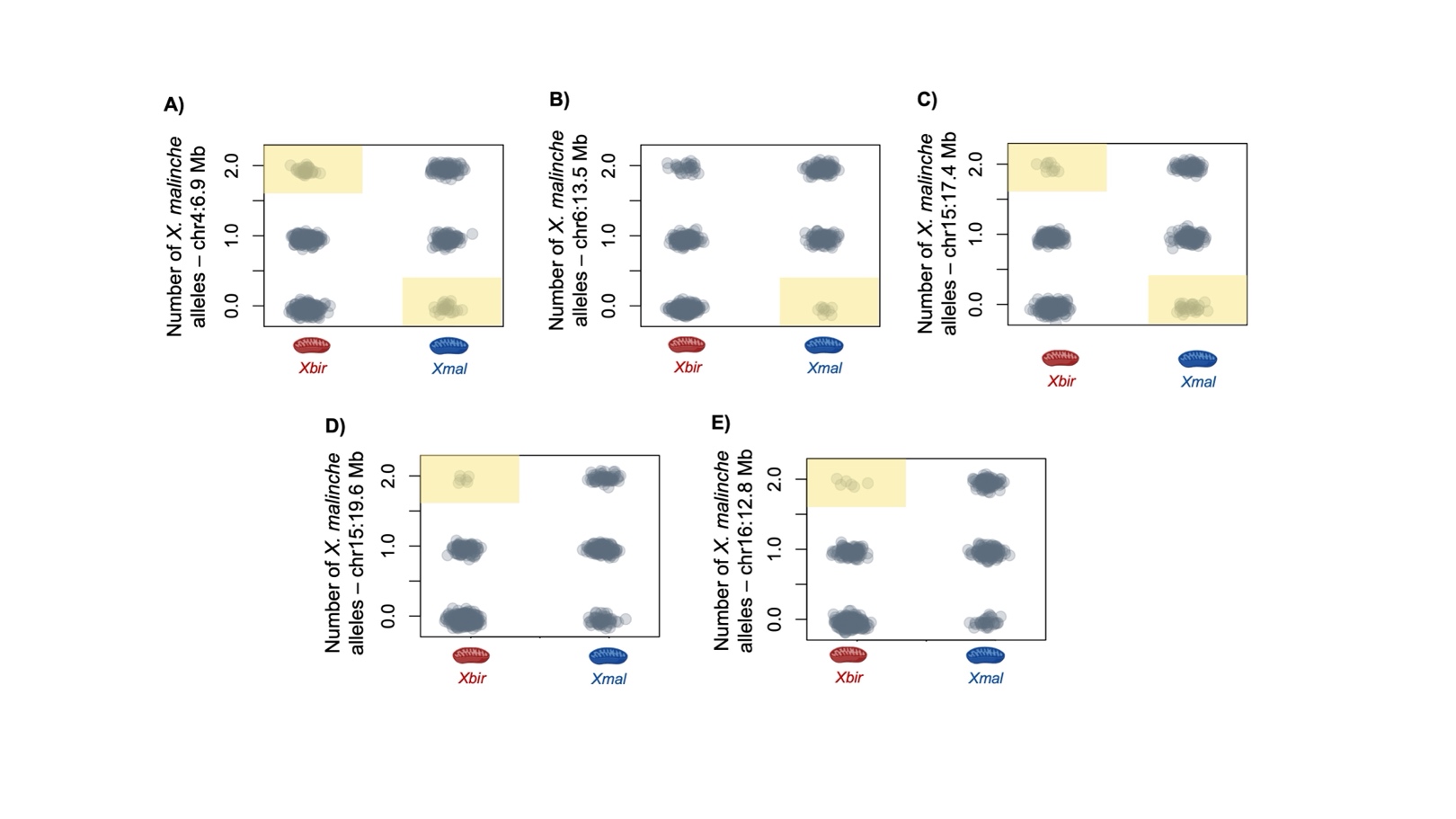

**Fig. S22.** Observed genotype combinations at mitonuclear interactions identified by admixture mapping dataset (**A-E**). For results for all other regions identified by admixture mapping, see Fig. 3 and Fig. S1. Each gray point indicates one adult individual in our admixture mapping population as a function of their mitochondrial haplotype. Red mitochondria on the x-axis indicates individuals with the *X. birchmanni* mitochondrial haplotypes (*Xbir*) and blue mitochondria indicates individuals with *X. malinche* mitochondrial haplotypes (*Xmal*). Genotype combinations that are significantly depleted based on comparisons to null simulations are highlighted in yellow (see Supporting Information 1).

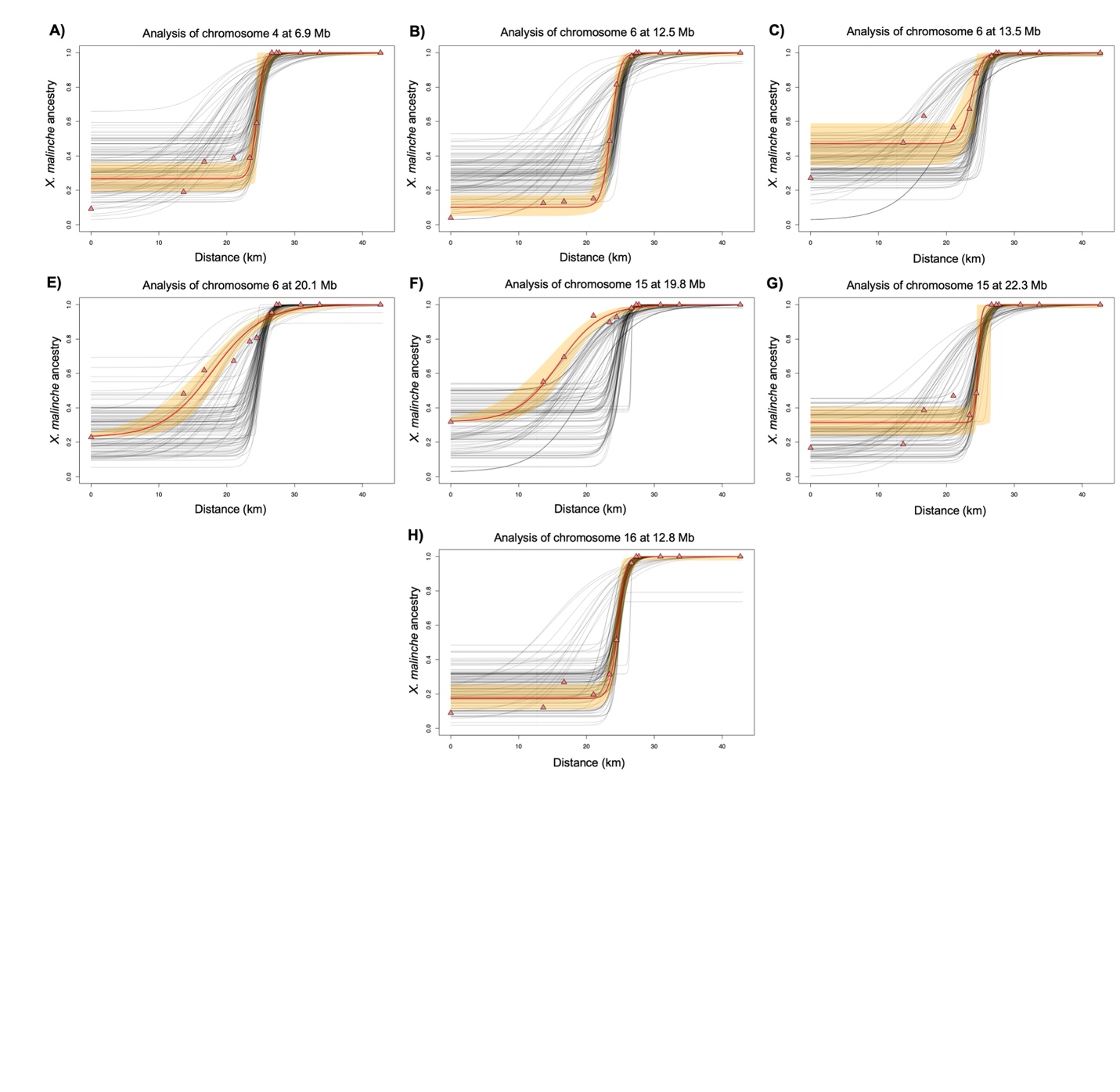

**Fig. S23.** Clinal changes in allele frequency across sampling sites in the Río Pochula for additional admixture mapping regions not shown in Fig. 5. SNP-specific clines were fit based on the marker closest to each admixture mapping peak. Cline models were fit using the HZAR software, the red line represents the model fit by HZAR to each focal locus, triangles show average ancestry in each population for the focal locus. Orange ribbons show 95% credible regions for the focal cline, and gray lines show model fit for 100 matched null markers (see Methods).

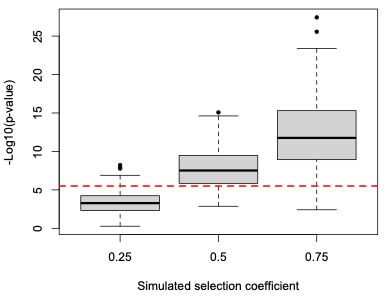

**Fig. S24.** Results of simulations evaluating power in our admixture mapping analyses. Simulations shown here model symmetrical selection on mitonuclear interactions using SELAM (see Supporting Information 8) at selection coefficients ranging from 0.25 to 0.75 with a dominance coefficient of 0.5. The red line indicates our genome-wide significant threshold for associations between mitochondrial and nuclear ancestry (10% false positive rate threshold). Note that while we expect to have good power to detect mitonuclear incompatibilities that are under extremely strong selection (s≥0.5), even with a sample size of 731 individuals we expect to have weak power to detect selection coefficients of 0.25. This suggests that we could be missing many mitonuclear incompatibilities with our mapping approach, depending on the true distribution of selection coefficients.

**Table S1.** Sampling sites and population information for locations on the Río Pochula used in geographical cline analysis. Additional information on samples included in cline analysis is available in Table S6.

| **Population name** | **Elevation** | **Average *X. malinche* ancestry genome-wide** | **Number of samples sequenced** |
| --- | --- | --- | --- |
| CULH | 1400 | 0.998 | 26 |
| TEXC | 860 | 0.998 | 23 |
| CSRL | 765 | 0.998 | 16 |
| POCH | 552 | 0.994 | 39 |
| PHLW | 542 | 0.995 | 19 |
| OCOT | 522 | 0.954 | 26 |
| ISLT | 442 | 0.661 | 36 |
| PICO | 438 | 0.556 | 39 |
| TULA | 405 | 0.512 | 33 |
| VCHO | 375 | 0.465 | 45 |
| ATMP | 330 | 0.386 | 37 |
| AREN | 221 | 0.227 | 39 |

**Table S2.** Summary of datasets and sample sizes used in the analyses presented in the main text.

| **Dataset** | **Main analysis** | **Data source** | **Data type** |
| --- | --- | --- | --- |
| *Admixture mapping dataset* | Admixture mapping of mitonuclear incompatibilities | Natural hybrids from the Calnali Low hybrid population: 359 previously collected, 372 collected for this study (731 total) | Low coverage whole-genome sequencing for local ancestry inference |
| *Segregation distortion analysis in F_2_ hybrids* | Identifying regions of the genome that significantly deviate from expected 50:50 admixture proportions in artificial hybrids with the *X. malinche* mitochondria | F_2_ lab-generated hybrids: 943 previously collected, 805 collected for this study (1748 total) | Low coverage whole-genome sequencing for local ancestry inference |
| *Size variation in F_2_ hybrids* | Testing for an effect of genotype at mitonuclear incompatibilities involving the *X. malinche* mitochondria on adult size | F_2_ lab-generated hybrids: 181 individuals from seven families | Low coverage whole-genome sequencing for local ancestry inference; measurements of adult standard length |
| *Embryo morphology & physiology* | Testing for an effect of genotype at mitonuclear incompatibilities involving the *X. malinche* mitochondria on embryo phenotypes | Reanalysis of 250 F_2_ lab-generated hybrid embryos from Moran et al. 2024 | Low coverage whole-genome sequencing for local ancestry inference; heart morphology; heart rate; embryo size; yolk size; respiration rate. |
| *Samples from natural hybrid populations with fixed mitochondria* | Examining evidence for variation in local ancestry in natural hybrid populations consistent with selection against mitonuclear incompatibilities | Reanalysis of data previously collected from the Acuapa population (*X. birchmanni* mitochondria; N=97) on the Río Huazalingo and the Tlatemaco population (*X. malinche* mitochondria; N=96) on the Río Claro | Previously published local ancestry data from Langdon et al. 2022; 2024 |
| *Clinal sampling from the Río Pochula* | Examining evidence for selection against mitonuclear incompatibilities based on clinal changes in local ancestry | Natural hybrids from 12 sites along the Río Pochula; total of 378 individuals sampled. | Low coverage whole-genome sequencing for local ancestry inference |

*Provided as an attached excel file:*

**Table S3.** All genes in each admixture mapping peak interval.

**Table S4**. Results of analysis with a linear mixed model to investigate the relationship between standard length and ancestry on chromosome 6 in lab-raised hybrids. Adjusted p-values are based on a Bonferroni correction for the total number of tests. Note that due to strong linkage between these regions in early generation hybrids (Fig. S21), we cannot distinguish which region on chromosome 6 impacts standard length in our data, so report statistics for analyses of each region separately. The likely gene associated with each region is noted parenthetically (see Table 1).

| Genotype | p-value | Adjusted p-value |
| --- | --- | --- |
| Position: 12.51 Mb (*ndufa13*) | | |
| *X. birchmanni* | 4.12e-09 | 1.24e-08 |
| Heterozygous | 1.73e-09 | 5.18e-09 |
| *X. malinche* | 1.33e-08 | 3.98e-08 |
| Position: 13.30 Mb (*mterf4*) | | |
| *X. birchmanni* | 3.90e-07 | 1.17e-06 |
| Heterozygous | 2.36e-07 | 7.07e-07 |
| *X. malinche* | 6.69e-07 | 2.01e-06 |
| Position: 20.25 Mb (*atp5mg*) | | |
| *X. birchmanni* | 7.94e-05 | 0.00024 |
| Heterozygous | 4.98e-05 | 0.00015 |
| *X. malinche* | 1.41e-04 | 0.00042 |

**Table S5.** Results of Welch’s t-test (two-sided) for differences in mitochondrial flux control factors (FCFs) between the two hybrid and the two parental groups.

|  | **FCF** | ***d.f.*** | ***t*-value** | ***p*-value** |
| --- | --- | --- | --- | --- |
| **Hybrids** | Complex I activity | 11.963 | 0.66048 | 0.5215 |
|  | Rotenone-CI inhibition | 4.1725 | -1.0731 | 0.3413 |
|  | Complex II activity | 11.554 | 0.74479 | 0.4713 |
|  | Malonate-CII inhibition | 3.621 | 0.65521 | 0.5516 |
|  | Oligomycin-CV inhibition | 10.688 | -0.27437 | 0.789 |
|  | Complex IV activity | 6.3624 | 0.52791 | 0.6155 |
| **Parentals** | Complex I activity | 27.333 | 0.23287 | 0.8176 |
|  | Rotenone-CI inhibition | 18.354 | -0.1386 | 0.8912 |
|  | Complex II activity | 24.432 | -1.4426 | 0.1618 |
|  | Malonate-CII inhibition | 19.289 | -0.76542 | 0.4533 |
|  | Oligomycin-CV inhibition | 20.992 | -0.57761 | 0.5697 |
|  | Complex IV activity | 20.613 | -1.5811 | 0.1291 |

*Provided as an attached excel file*

**Table S6.** Individual sample information for clinal samples collected from the Río Pochula. Month of collection is indicated by roman numerals and sex if known is indicated. M - male, F - female, U - juvenile of unknown sex.

**Supporting Information References**
